# A Zero-Inflated Hierarchical Generalized Transformation Model to Address Non-Normality in Spatially-Informed Cell-Type Deconvolution

**DOI:** 10.1101/2024.06.24.600480

**Authors:** Hunter J. Melton, Jonathan R. Bradley, Chong Wu

**Affiliations:** Department of Biomedical Data Science, Geisel School of Medicine at Dartmouth, Hanover, New Hampshire, U.S.A.; Department of Statistics, The University of Missouri, Columbia, Missouri, U.S.A.; Department of Biostatistics, University of Texas MD Anderson Cancer Center, Houston, Texas, U.S.A.

**Keywords:** Bayesian, spatial transcriptomics, uncertainty quantification, zero-inflation

## Abstract

Oral squamous cell carcinomas (OSCC), the predominant head and neck cancer, pose significant challenges due to late-stage diagnoses and low five-year survival rates. Spatial transcriptomics offers a promising avenue to decipher the genetic intricacies of OSCC tumor microenvironments. In spatial transcriptomics, Cell-type deconvolution is a crucial inferential goal; however, current methods fail to consider the high zero-inflation present in OSCC data. To address this, we develop a novel zero-inflated version of the hierarchical generalized transformation model (ZI-HGT) and apply it to the Conditional AutoRegressive Deconvolution (CARD) for cell-type deconvolution. The ZI-HGT serves as an auxiliary Bayesian technique for CARD, reconciling the highly zero-inflated OSCC spatial transcriptomics data with CARD’s normality assumption. The combined ZI-HGT + CARD framework achieves enhanced cell-type deconvolution accuracy and quantifies uncertainty in the estimated cell-type proportions. We demonstrate the superior performance through simulations and analysis of the OSCC data. Furthermore, our approach enables the determination of the locations of the diverse fibroblast population in the tumor microenvironment, critical for understanding tumor growth and immunosuppression in OSCC.

## 1 Introduction

Oral squamous cell carcinomas (OSCC) are the most common head and neck cancer, comprising over 90% of oral cavity mucosal malignancies (Markopoulos, 2012). OSCC progression is often undetected until advanced stages, contributing to five-year survival rates of approximately 50% (Markopoulos, 2012). Spatial transcriptomics (ST) technologies, which provide spatial localization of gene expression (Moses and Pachter, 2022), offer new opportunities to characterize tumor cellular and molecular landscapes. The recent OSCC ST dataset by (Arora et al., 2023) enables detailed investigation of the tumor microenvironment (TME) and spatial distribution of cell types, providing novel perspectives on tumorigenesis and cellular interactions (Moses and Pachter, 2022).

However, the application of ST in OSCC research encounters notable hurdles with spatial barcoding-based methods, such as the 10X Visium platform used to capture the OSCC ST data (Arora et al., 2023). These methods collect gene expression measurements from tissue locations containing multiple cells of diverse types (Ståhl et al., 2016), effectively quantifying average expression levels across cells at each location. Consequently, cell-type deconvolution, which probabilistically infers cell type proportions at each location, is crucial for determining spatial cell type distributions and enabling subsequent analyses (Ma and Zhou, 2022).

Despite these advances, existing approaches often overlook zero inflation endemic to ST data (Li et al., 2021; Zhao et al., 2022). For example, CARD (Ma and Zhou, 2022) assumes the spatially resolved gene expression data are normally distributed, disregarding zero-inflation and ties, which may result in reduced performance due to mismatched data and modeling assumptions. While Zhao et al. (2022) argue that zero-inflation can be addressed by controlling for cell-type composition or by analyzing spatially homogeneous tissue with standard count models such as a Poisson or negative binomial, their analysis focuses on scenarios where spatial heterogeneity is controlled rather than modeled. In contrast, methods like CARD that assume normality of the raw or minimally processed data may benefit from transformations that address zero-inflation and ties simultaneously with spatial modeling.

Zero-inflation is substantial in the 12 OSCC samples (Arora et al., 2023), where sparsity ranges from 86%-91%, a pattern common across ST datasets (Moncada et al., 2020). While the high sparsity could be directly addressed by a zero-inflated Poisson model (Lambert, 1992) or zero-inflated Negative Binomial model (Greene, 1994), these are challenging and time-consuming given the spatial context and data size—over 15 million points per sample. Deterministic transformations, such as log(1+*ɛ*+*x*), may be applied to ST data prior to analysis, though their impact is minimal (Web Figure 3). Consequently, few spatial transcriptomic methods address zero-inflation (Li et al., 2021), and no cell-type deconvolution methods do so directly. Moreover, uncertainty quantification (UQ) in cell-type deconvolution often relies on model-based assumptions that may not align with the true distribution of the data. Fully Bayesian models enable posterior credible intervals for UQ but require computationally intensive MCMC methods that renders them impractical for high-dimensional ST datasets. In classical statistics, when data diverge from model assumptions (often normality), deterministic transformations, such as Box-Cox (Box and Cox, 1964) or Yeo-Johnson power transformations, (Yeo and Johnson, 2000) are employed to align data with the assumptions pre-analysis. However, classical deterministic transformations struggle with data containing many ties, as ties remain after transformation, resulting in persistent non-normality. Contrastingly, the hierarchical generalized transformation (HGT) (Bradley et al., 2020; Bradley, 2022; Bradley et al., 2023) introduces a probabilistic, rather than deterministic, transformation to data processing. The HGT is a new class of noisy transformations that breaks ties by adding a small amount of noise to the data, which is modeled via a Bayesian transformation. The HGT, however, has yet to be developed for zero-inflated data. Here, we introduce the Zero-Inflated Hierarchical Generalized Transformation (ZI-HGT), a novel adaptation designed to address zero inflation and ties in ST datasets. By integrating a type of zero-inflated Poisson model into the HGT framework, the ZI-HGT generates posterior replicates from the noisy transformation, mitigating zeros and ties to ensure transformed data better aligns with model assumptions (i.e., normality in CARD). The posterior replicates of the transformation are then used in place of the original data. The combination of zero-inflated modeling with HGT and CARD produces a Bayesian hierarchical model that enables UQ and is implementable on high-dimensional (over 15 million data points), multivariate spatial datasets, a scale at which current fully Bayesian alternatives for zero-inflated multivariate spatial data become computationally intractable (Musgrove et al., 2019).

The rest of the manuscript is organized as follows: Section 2 discusses the motivating OSCC data. Preliminary concepts and methodology are discussed in Sections 3 and 4, respectively. An empirical simulation study is presented in Section 5, which elucidates the viability of the model. Section 6 applies the model to OSCC data, presenting findings, and Section 7 provides conclusions and discussions. Figure 1 contains a schematic overview of the method.

**Figure 1:**
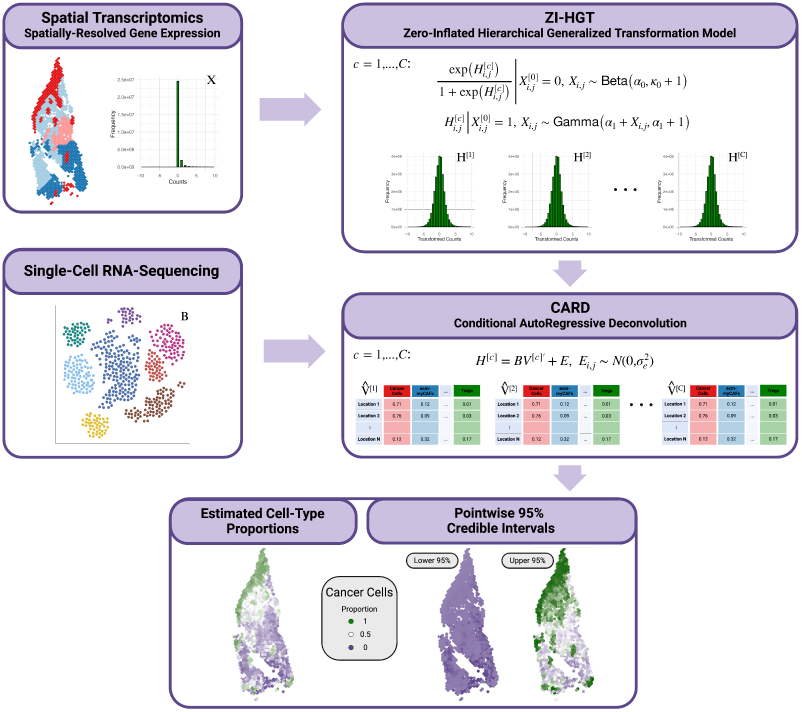
ZI-HGT + CARD Schematic Outline. The raw spatial transcriptomic data *X* is inputted into the ZI-HGT, generating *C* posterior replicates (*H*^[1]^,…, *H*^[^*^C^*^]^), which are interpreted as transformations of the original data. These transformations better fit the assumptions of CARD, namely that the data are normally distributed. CARD is then applied to each transformed dataset and a corresponding single-cell RNA sequencing (scRNA-seq) dataset *B*, yielding cell-type proportion estimates (*V̂*^[1]^,…, *V̂*^[^*^C^*^]^). These estimates are amalgamated into a single estimate *V̂*.

## 2 Data and Motivation

We analyze oral squamous cell carcinoma spatial transcriptomic (ST) data (Arora et al., 2023), consisting of 12 samples surgically resected from 10 patients’ oral cavities. Nine tissue layers were annotated by a pathologist: squamous cell carcinoma (SCC), lymphocyte-positive stroma, lymphocyte-negative stroma, normal mucosa, glandular stroma, muscle, keratin, and artery/vein, and an artifact layer. ST was conducted using the 10x Genomics Visium platform, which samples approximately 10 cells per location. Across all samples, reads were obtained for an average of 16, 763 unique transcripts over 2, 073 spatial locations. Additionally, our analysis incorporates a reference single-cell RNA-sequencing (scRNA-seq) dataset (GSE103322) (Puram et al., 2017). Data collection and preparation are detailed in Web Appendix A. As an eaxmple, we consider OSCC ST Sample 1 and note that the other samples are similar (Web Table 1). Sample 1 contains read counts for 15,844 transcripts across 1,131 spatial locations. Web Table 2 presents gene expression read count frequencies (see Web Tables 3-13 for other samples). We highlight two key features of the dataset. First, zero inflation is extremely high: 16,384,736 of 17,919,564 (91.4%) read counts are zero. Second, the count data contains many ties. These characteristics underscore the limitations of current analytical approaches like CARD, which assumes normally distributed ST data, an assumption rendered untenable by zero-inflation and abundant ties. To address these challenges, we introduce a novel zero-inflated HGT (a noisy transformation aimed at breaking ties and addressing zero-inflation) in the following section to address the non-normality of the data before conducting cell-type deconvolution with CARD.

## 3 Preliminaries

### 3.1 Notation

Let **X** be the *G* × *N* gene expression matrix from an OSCC sample (Arora et al., 2023), containing transcript reads for *G* cell-type informative genes measured at *N* spatial locations. Denote the (*i, j*)-th element *X_i,j_*. Let **H**(**X**), or **H** for brevity, be the noisy transformation of **X**, **X**^[0]^ be a set of indicators for positive gene expression in **X** (i.e., 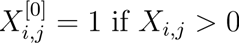), and **X**^(^*^B^*^)^ be a data augmentation variable that facilitates the implementation of a zero-inflated model without expensive Markov chain Monte Carlo (MCMC) sampling. We employ corresponding scRNA-seq data (Puram et al., 2017) as a deconvolution reference and select informative genes for distinguishing between cell types. Let **B** be the *G* × *K* single-cell gene expression matrix (the reference basis matrix), where *G* is again the number of cell-type informative genes and *K* is the number of cell types. Lastly, let **V** be the *N* × *K* cell-type proportions matrix, where the rows of **V** contain the proportions of the *K* cell types for each spatial location. In the OSCC ST data, *G* and *N* may vary slightly from sample-to-sample, but *K* = 14. Futhermore, we introduce the following shorthand notation for probability distributions. “Point Mass” is a short-hand for the point mass distribution, and “TruncPoisson” is a short-hand for the zero-truncated Poisson distribution.

### 3.2 Review of CARD

CARD (Ma and Zhou, 2022) is an empirical Bayesian model that applies the following non-negative matrix factorization model for **X**:

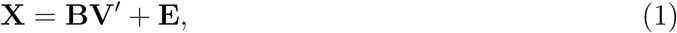

where **E** is the *G* × *N* residual error matrix and 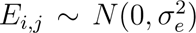. *B* contains the average gene expression level in each cell type from the scRNA-seq reference data for all “cell-type informative” genes, classified as such if their mean expression level in any cell type is at least 1.25 log-fold higher than the average across all other cell types (see Web Appendix H for details). To consider spatial colocalization of cell types, a conditional autoregressive (CAR) (Besag, 1974) model serves as the prior for columns of the cell-type proportions matrix **V**. For spatial location *i* and cell type *k*, the full-conditional distribution for *V_i,k_* is

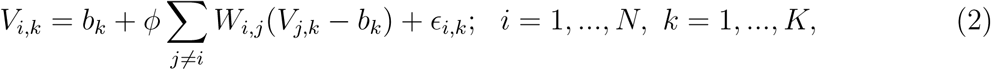

where *V_i,k_* denotes the proportion of cells at location *i* belonging to cell type *k*, *b_k_* is the cell-type intercept representing the average cell-type composition for cell type *k* across all locations, and *W_i,j_* refers to the weight for learning the cell-type composition at location *i* using the information at location *j*. The parameter *ϕ* represents the spatial autocorrelation, determining the strength of the spatial correlation in the cell-type composition, and *ɛ_i,k_* represents the residual error such that 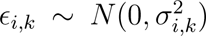. The CAR model represents the proportion of cells at location *i* in cell type *k V_i,k_* as a weighted sum of cell type *k* proportions at the other locations, enabling spatial sharing of cell-type composition information.

Notably, Equation (1) assumes normality of the) spatially resolved gene expression **X**, which conflicts with the high sparsity and prevalent ties in the OSCC data, as highlighted in Section 2. CARD incorporates location-level normalization, dividing each read count in *X* by the sum of the read counts at that location, in an attempt to address potential technical artifacts. While this procedure alleviates some ties, it does not impact zeros, so high sparsity (minimum 86% across the 12 OSCC samples) remains (Web Figure 43).

Rather than sample from the posterior predictive distribution (Equation (9) of the supplementary material of Ma and Zhou (2022)), CARD applies an algorithm that iterates through **V***_k_* and the hyperparameters to minimize the negative log-likelihood. This computationally efficient approach constructs point estimates of the cell-type proportions but precludes UQ.

### 3.3 Review of HGT

Models like CARD assume Gaussian data, which is unrealistic given the prevalence of ties in the OSCC dataset. Adding noise to the data produces a new (transformed) dataset without ties, rendering the Gaussian assumption more reasonable. To accomplish this, the HGT adds small amounts of noise to count data to break ties that fully accounts for added uncertainty from the noisy transformation. The HGT implements a fully Bayesian model that overfits the original data (Bradley, 2022), with posterior predictive replicates interpreted as the noisy transformation. By “overfitting,” the posterior replicates are close (in absolute difference) on average to a deterministic transformation of the data on some scale. The motivation for overfitting is that the added noise should be “small” enough to break ties without distorting the underlying signal or introducing substantial bias into downstream analyses.

Recall that **H** = **H**(**X**) denotes the noisy transformation of the original data **X**, and *H_i,j_* = {**H**}*_i,j_*. The current use of the HGT assumes *X_i,j_|H_i,j_* ∼ Poisson (*H_i,j_*) and *H_i,j_* ∼ Gamma (*α, κ*) so that the posterior distribution *H_i,j_|X_i,j_* ∼ Gamma (*α* + *X_i,j_, κ* + 1). When, for example, *α* ≈ *κ*, the posterior replicates of *H_i,j_|X_i,j_* have mean *Z_i,j_* that overfits the original data *X_i,j_*. This model “overfits” as the posterior summary recover the original data values. CARD is then applied to transformation **H** in place of **X**. Consider the following illustration. Let *α* = *κ* = 0.1, and let *X_i,j_* = 1. Then, as *H_i,j_|X_i,j_* ∼ Gamma (*α* + *X_i,j_, κ* + 1), 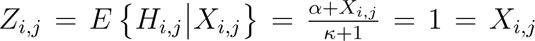, so the transformation mean equals the original data. More generally, *Z_i,j_* = *E*{*H_i,j_|X_i,j_*} → *X_i,j_* as *α, κ* → 0 or *X_i,j_* → ∞.

In the HGT, the added noise is parameterized as presented previously rather than a simpler transformation for two key reasons. First, as discussed in Bradley et al. (2020), HGT leverages signal-to-noise cross-covariance for better prediction (see Web Appendix D for details). Second, the conjugacy of the transformation enables direct sampling of the transformation posterior, significantly reducing computational costs. In fact, (Bradley, 2022) and (Bradley et al., 2023) compared the HGT framework to traditional Bayesian hierarchical models in various non-Gaussian settings, finding minimal differences in prediction and estimation but massive computational gains with the HGT. However, the presented HGT is suboptimal, as it fails to address the known zero-inflation in the OSCC ST data. Simulations suggest that ignoring zero-inflation in an HGT yields minimal improvement (Web Figure 3).

## 4 Methodology

### 4.1 Zero-inflated hierarchical generalized transformation (ZI-HGT)

When applying the ZI-HGT to CARD (Ma and Zhou, 2022), we address two features of the OSCC data: high zero inflation and ties. We consider two observed datasets: the original OSCC ST data **X** and **X**^[0]^. The *G* × *N* matrix **X**^[0]^ has (*i, j*)-th element 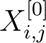 representing an indicator of positive gene expression for gene *i* at location *j*, i.e., 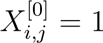 for positive gene expression and 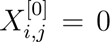 otherwise. We introduce 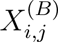, where the superscript “B” stands for Bernoulli and 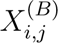 will facilitate the implementation of a Bayesian zero-inflated model with a posterior distribution that is easy to directly sample from. We assume the hierarchical model:

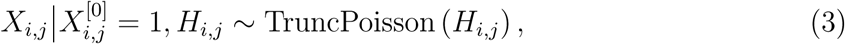

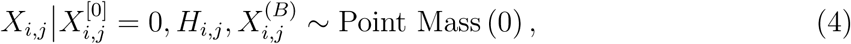

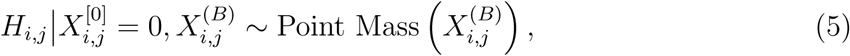

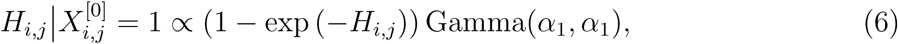

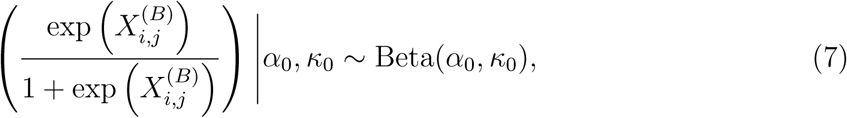

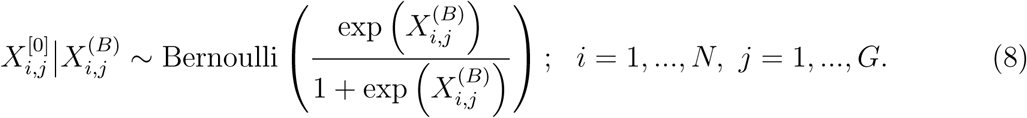

“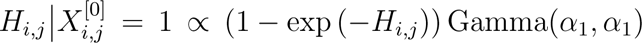” represents the conditionally specified and unnormalized prior:

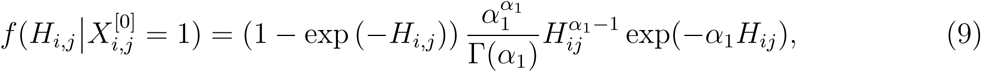

where a gamma probability density function is multiplied by (1 − exp (−*H_i,j_*)) to offset the normalizing constant in the truncated Poisson likelihood. One could similarly replace the truncated Poisson with the truncated negative binomial and modify the prior in (6) to be a weighted logit-beta prior distribution. The negative binomial is popular for modeling counts as it allows modeling over-dispersion. However, the HGT is a class of noisy transformations, where the transformations arise from a posterior distribution of a model that overfits the data, and posterior replicates are used as transformed data in another model (i.e., CARD). Questions of overdispersion are hence more appropriate for the model applied to the transformations (i.e., CARD), and CARD accounts for overdispersion via spatial covariances.

In Equation (3), non-zero count-valued OSCC ST data is modeled as truncated Poisson (⩾ 1) distributed, and in Equation (4) zero-valued OSCC ST data is modeled as a point mass at zero. Zero-inflated models adopt a similar strategy but allow some zeros to be modeled in a Poisson mixture component. We instead use the truncated Poisson and unnormalized prior for *H_i,j_*, (1 − exp (−*H_i,j_*)) Gamma(*α*_1_*, α*_1_) to achieve a closed-form posterior distribution for direct sampling (Equations (10) and (11) below). The prior distributions for the transformation *H_i,j_*, Equations (6) and (7), are conditionally specified on the observed 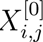. This choice produces a posterior distribution for *H_i,j_* with a closed form and the desired overfitting property (see Section 3.2 for motivation, and Web Figure 42 for a posterior transformation **H** of OSCC Sample 1 split by raw data value *X_i,j_*).

The posterior transformation in closed-form (derived in Web Appendix C) is given by

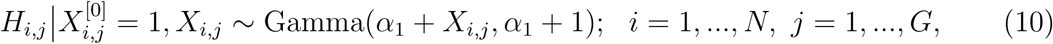

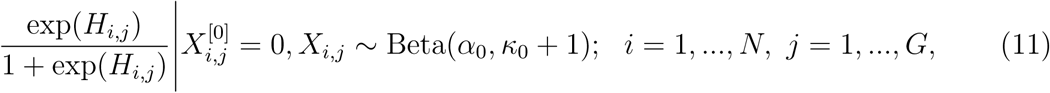

which has mean 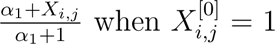 and mean 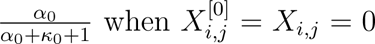, overfitting the data. The ZI-HGT can be sensitive to hyperparameter choices, so discrete ranges for *α*_0_, *κ*_0_, and *α*_1_ are considered. Specific values are chosen to minimize the WAIC (Watanabe, 2010), a generalized version of the AIC for model selection. In simulations, WAIC-chosen hyperparameters result in similar performance to oracle-chosen (RMSE-minimizing) hyperparameters (Web Figure 2). Further details and practical recommendations for choosing hyperparameters are discussed in Web Appendix C.

**Figure 2:**
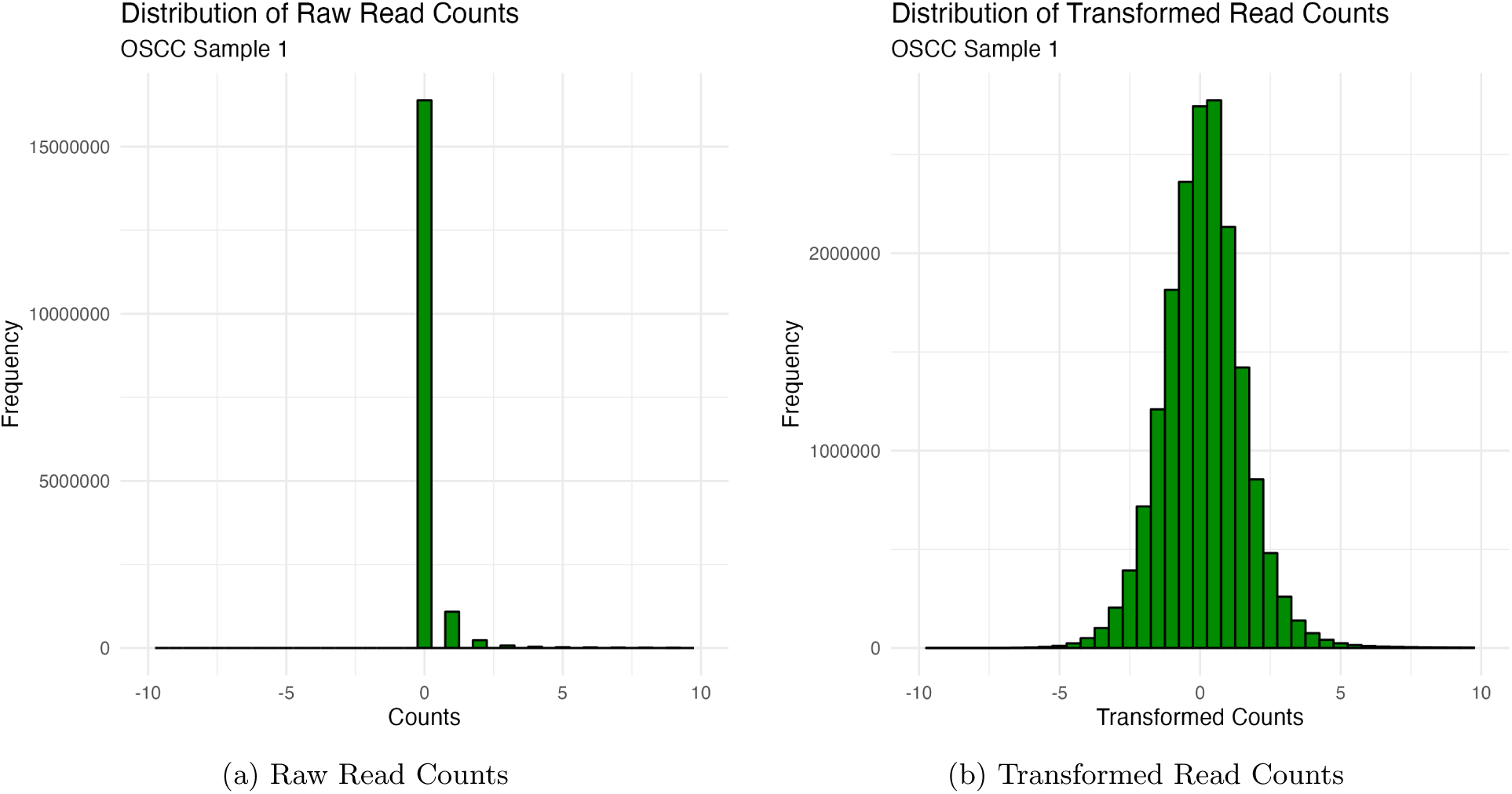
Distribution of the gene expression read counts for OSCC Sample 1. a) Raw read counts. b) A single posterior replicate of the transformed read counts.

Upon marginalizing 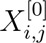 in (4) and (3), we come to the expression

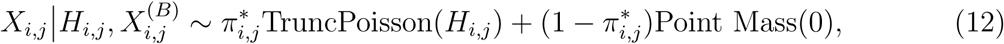

where 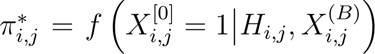 (detailed in Web Appendix E), and the expression in Equation (12) is of the form of a zero-inflated model (Lambert, 1992). While traditionally a Poisson distribution would replace the truncated Poisson, our goal differs from standard zero-inflated models. We aim to produce posterior replicates of *a transformation* for subsequent analysis rather than directly using posterior summaries of *H_i,j_*. Using posterior replicates enables uncertainty quantification of estimated cell-type proportions downstream. Thus, the truncated Poisson distribution is appropriate, yielding Equations (10) and (11) and hence a noisy transformation *H_i,j_* that approximates the original data *X_i,j_* on some scale.

### 4.2 Applying ZI-HGT to cell-type deconvolution

Denote the *c*-th replicate of **H** = {*H_i,j_*} from Equations (10) and (11) with 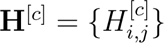 for *c* = 1*,…, C*. Following the HGT strategy, we assume

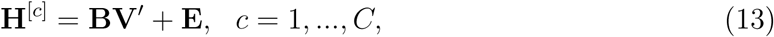

where **E** = {*E_i,j_*} represents independent normal random variables with constant variance, **V** is the non-negative cell-type proportions with columns following a conditional autoregressive (CAR) (Besag, 1974) model, and **B** is the reference basis matrix. Algorithm 1 details the implementation.

The ZI-HGT model is defined by Equations (3-8), (13) and (2) (Bradley, 2022). From Bradley (2022), the BHM expression requires careful consideration of normalizing constants as Equations (3-8) condition on **H**^[^*^c^*^]^, while Equation (13) includes **H** but not as a condition. While the ZI-HGT alone does not model spatial variability, the combined ZI-HGT + CARD does through CARD. Web Appendix C provides the complete BHM for ZI-HGT + CARD. Since no MCMC is required, the full method with CARD can be run efficiently by generating each of the posterior predictive replicates of the transformed data 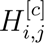 then computing the cell-type proportions estimates 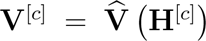 in parallel. The estimator *V̂* is the function that minimizes the least squares 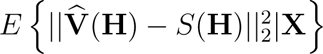, where *S* is a generic real-valued estimator and || · ||_2_ is the *L*_2_ norm. The transformed OSCC data replicates **H** align significantly better with CARD’s normality assumption than the original data **X** (Figure 2), containing no zeros or ties and substantially reducing the large right tail. While normality tests like Kolmogorov-Smirnov would be useful, they require independent data, whereas our data are multivariate and spatially dependent.

Consequently, ZI-HGT-generated estimates are more accurate at no additional computational cost, as demonstrated in our simulation study. We recommend *C* = 100 replicates. UQ is accomplished through replicates of Algorithm 1, the iterated variance formula, and a Taylor series argument to construct pointwise credible intervals for **V̂**. UQ is detailed in Web Appendix F. Though it is not recommended due to the unclear cause of the zero-inflation, Web Appendix G presents the inverse transformation to return predictions back to the original scale of the data for completeness.

**Algorithm 1:**
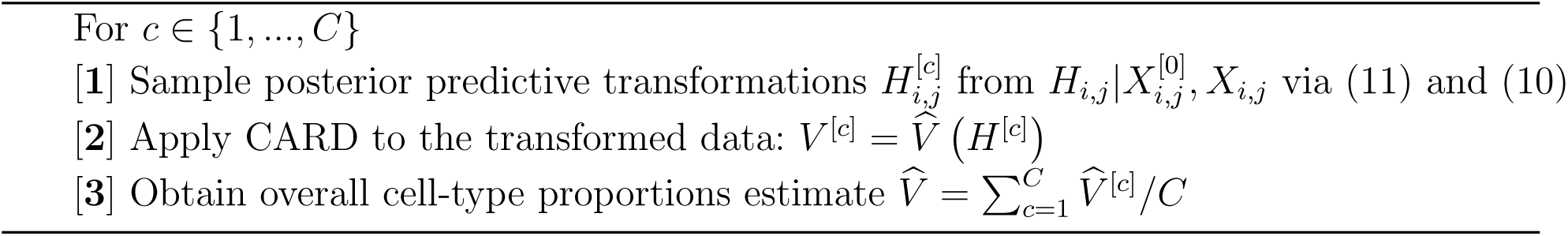
Applying the ZI-HGT to CARD

## 5 Simulations

We conducted a rigorous simulation study using the OSCC data to evaluate the performance of the ZI-HGT + CARD across multiple settings, including varying sparsity levels and numbers of cell types in the ST data. We compared our approach with existing methods and a simple deterministic transformation to ensure comprehensive evaluation. The study focuses on OSCC Sample 2, which we consider representative of most samples, though sample choice has minimal impact (Web Figure 2). Sample 2 contains data from 1, 749 locations and 15, 624 transcripts. Web Appendix B provides additional details regarding the simulations.

### 5.1 Simulation Results

We first examined the impact of ST data sparsity on ZI-HGT + CARD performance, measured by the root mean square error (RMSE) in Figure 3. Higher sparsity in the simulated ST data resulted in stronger performance of the ZI-HGT + CARD relative to CARD alone. Sparsity levels of the data are determined the “library factor” and “*ϕ* factor” simulation hyperparameters; specific choices for both are presented in Figure 3 and detailed in Web Appendix B. At sparsity level 1 (median sparsity 89.8%, representing realistic ST data), the median RMSE reduction of ZI-HGT + CARD beyond CARD was 6.6%. At sparsity level 2 (median sparsity 77.9%), the median reduction was 4.6%. This trend is expected and occurs as higher sparsity creates less normally distributed ST data, further deviating from CARD’s assumptions and reducing its accuracy. The ZI-HGT addresses zero-inflation, so its impact increases as the ST data becomes “less normal.” We discuss the performance of ZI-HGT + CARD primarily in terms of sparsity, but the median proportions of data points tied at 1, 2, 3, 4, and 5 are given for each sparsity level for additional context.

**Figure 3:**
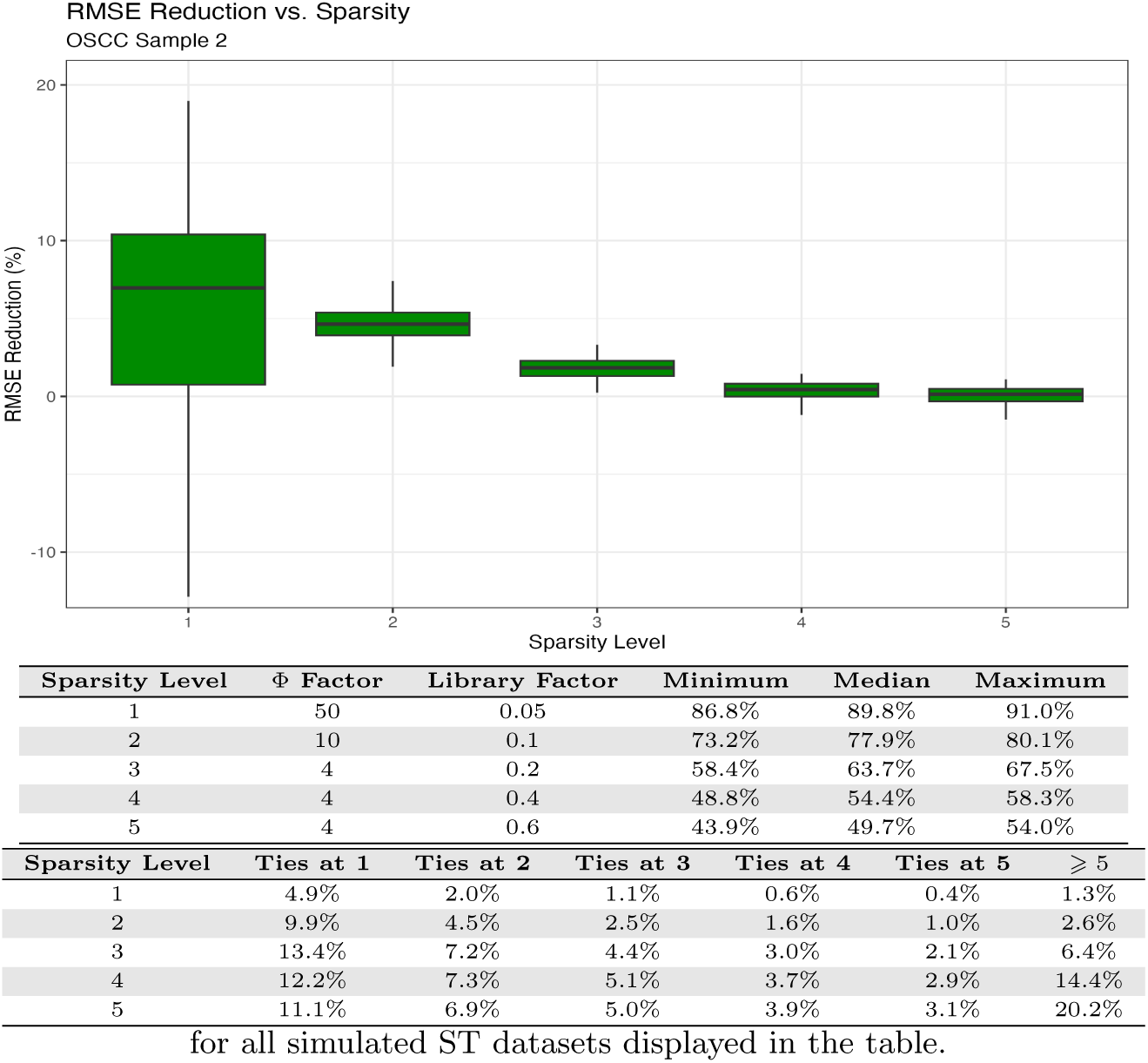
RMSE reduction vs. sparsity level of simulated ST data. RMSE reduction for the ZI-HGT + CARD across 100 simulated ST datasets and various sparsity levels. The sparsity levels in the boxplot have corresponding simulation hyperparameter factors, minimum, median, and maximum sparsity levels, and median proportions of data equal 1, 2, 3, 4, and 5 and ⩾ 5

Next, we investigated how the number of distinct cell types affects performance (Web Figure 1). While median RMSE reduction remains consistent across varying cell type numbers, variance decreases as the number of cell types decreases. This expected outcome occurs because fewer cell types create more compact, simplified simulated ST data. We note that sparsity is unrelated to the number of included cell types, with median sparsity levels ranging from 88.2% to 89.2% across datasets with different cell type counts.

To ensure robustness given our focus on OSCC Sample 2, we examined the impact of sample choice for simulating ST data (Web Figure 2). We generated simulated ST data from SPARSim-generated scRNA-seq data using “realistic” settings (sparsity level 1). Our methods showed consistent RMSE reduction, with a median of 6.6%, though the variance was higher in Samples 8, 9, and 10, which were particularly homogeneous. These results reinforce our confidence in using OSCC Sample 2 as a representative choice for simulations. We also compared ZI-HGT hyperparameters selected using WAIC vs. the oracle hyperparameters that minimize RMSE in simulated datasets based on each OSCC sample (Web Figure 2). Minimizing the WAIC yielded reasonable hyperparameter choices, with median RMSE reduction differences between WAIC-chosen and oracle models consistently small, nearly always within 1%. While the ZI-HGT can be somewhat sensitive to the choices of hyperparameters, hyperparameters that produce significantly worse performance consistently had higher WAIC values (Web Figure 46), indicating that the WAIC selects hyperparameters well. Practical recommendations for choosing hyperparameters are discussed in Web Appendix C.

We compared the ZI-HGT + CARD to several alternatives: a simple deterministic transformation before CARD, HGT + CARD (the ZI-HGT without addressing zero-inflation), imputation with ALRA (Linderman et al., 2022) + CARD, denoising with MIST (Wang et al., 2022), and common cell-type deconvolution methods SPOTlight (Elosua-Bayes et al., 2021), SpatialDecon (Danaher et al., 2022), and STdeconvolve (Miller et al., 2022) (Web Figure 3). The deterministic transformation *log* (1 + *ɛ* + *X_i,j_*) is chosen to mimic the ZI-HGT: the log mitigates right-skewness and the *ɛ* addition addresses zero-inflation. This transformation improved RMSE by 2.1% beyond CARD alone but fell short of the ZI-HGT + CARD’s 6.6% improvement. Clearly, a deterministic transformation cannot address ties in the OSCC ST data, nor does it allow UQ. HGT + CARD improved RMSE by 1.4%, emphasizing the need to address zero-inflation. Neither ALRA imputation nor MIST denoising + CARD significantly impacted performance. Median RMSE reductions were −0.1% and 0.2% respectively, underscoring the need for specific ST pre-processing techniques to transform the data to approximate normality. Other cell-type deconvolution methods performed notably worse than CARD alone; the best (SPOTlight) had 15% higher RMSE. Finally, we examined the impacts of increasing the number of ties at small nonzero values in the data and altering sequencing depth on performance. To generate simulated ST data with additional ties at nonzero values, we randomly selected one cell type in the underlying simulated scRNA-seq data and added 1, 2, or 3 to the gene expression of a randomly selected 50% of genes, inducing ties at nonzero values in simulated ST data. The ZI-HGT improved performance beyond CARD alone by 7.8% when 1 was added, and 8.1% when 2 or 3 were added, demonstrating robustness to additional ties. To explore changes in sequencing depth, we simulated ST data with −50%, −30%, −10%, +10%, +30%, and +50% depth and applied the ZI-HGT + CARD (Web Figure 48). The median improvement of the ZI-HGT + CARD vs. CARD alone is greater than 5% for the data with −10%, +10%, and +30% sequencing depth, and is approximately 4% with +50% sequencing depth, likely due to decreased sparsity (86.1%). Performance decreased when sequencing depth decreased dramatically (−30%, - 50%). Decreased sequencing depth leads to lower signal, which is overwhelmed by the ZI-HGT’s added noise. Thus, we do not recommend applying the ZI-HGT when ST data has few cells per location. Fortunately, ST techniques typically capture approximately 10 cells per location or achieve near single-cell resolution (Moses and Pachter, 2022).

### 5.2 Applying the ZI-HGT + CARD to CARD’s Simulations

To demonstrate our method’s viability in an unbiased manner, we applied the ZI-HGT + CARD to CARD’s exact simulations without modifications. These simulations, constructed similarly to ours, are based on two freely available mouse olfactory bulb (MOB) datasets: a scRNA-seq reference (Zeisel et al., 2018) and an ST dataset (Ståhl et al., 2016). The simulated ST data is much less sparse than real ST datasets, with median sparsity of 68.6% versus 88.5% in the real OSCC ST datasets. Web Appendix B details the simulation procedure.

Five distinct simulation scenarios are considered: 1) The scRNA-seq data is applied as the reference basis matrix for cell-type deconvolution (standard approach). 2) One cell type is removed from the scRNA-seq reference. 3) One cell type (blood cells) is added to the scRNA-seq reference. 4) Two cell-types are merged in the scRNA-seq reference. 5) A mismatched scRNA-seq reference dataset from a different sequencing platform, mouse brain tissue, and mouse age is used (Zeisel et al., 2018). Following CARD, additional robustness comes from considering “noisy” data where the dominant cell type of a given percentage *p_n_* ∈ {0%, 20%, 40%, 60%} of locations is randomly chosen.

Figure 4 displays ZI-HGT + CARD’s performance across 100 simulated ST datasets for all five CARD simulation scenarios with *p_n_* = 0% (Web Figures 5 - 7 show *p_n_* ∈ {20%, 40%, 60%}). The ZI-HGT + CARD is less impactful than in the realistic OSCC simulations (2.1% median RMSE reduction in scenario 1), which is expected given the unrealistically low sparsity of the CARD simulated dataset. For scenario 1 (*p_n_* = 0%), the median sparsity is just 68.6%, significantly lower than the 86%-91% in the OSCC data. As Figure 3 demonstrates, the ZI-HGT is more impactful in sparser datasets where zero-inflation is more pronounced. Overall, The ZI-HGT + CARD outperforms CARD alone for nearly every noisiness percentage *p_n_* and scenario. The exception is scenario 5 with the highly mismatched scRNA-seq reference. As the ZI-HGT adds noise during transformation, the additional noise from the mismatched reference is amplified, slightly reducing accuracy. However, scenario 5 represents an uncommon, suboptimal edge case as using reference datasets from different tissues and platforms are generally avoided in practice. For mismatched references, using CARD alone is advised, and caution when interpreting results is encouraged due to increased risk of inaccuracy. We also compared the ZI-HGT + CARD to CARD alone applied to simulated data with ±10% sequencing depth (number of gene expression read counts per location) (Web Figures 4-7). As a sanity check, our analysis confirms that enhanced sequencing depth improves deconvolution accuracy.

**Figure 4:**
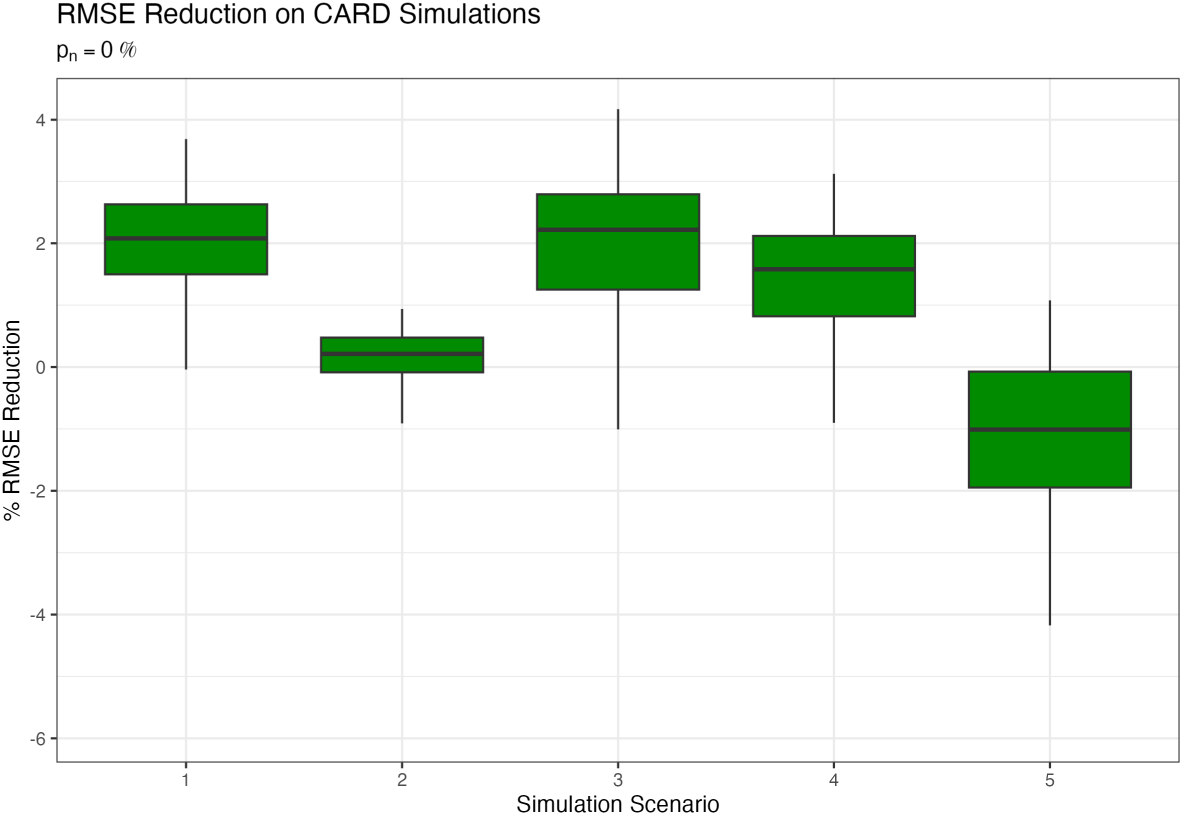
RMSE reduction on CARD simulations. RMSE reduction for the ZI-HGT + CARD across 100 simulated MOB ST datasets and various simulation scenarios following the CARD simulation setup with *p_n_* = 0%.

**Figure 5:**
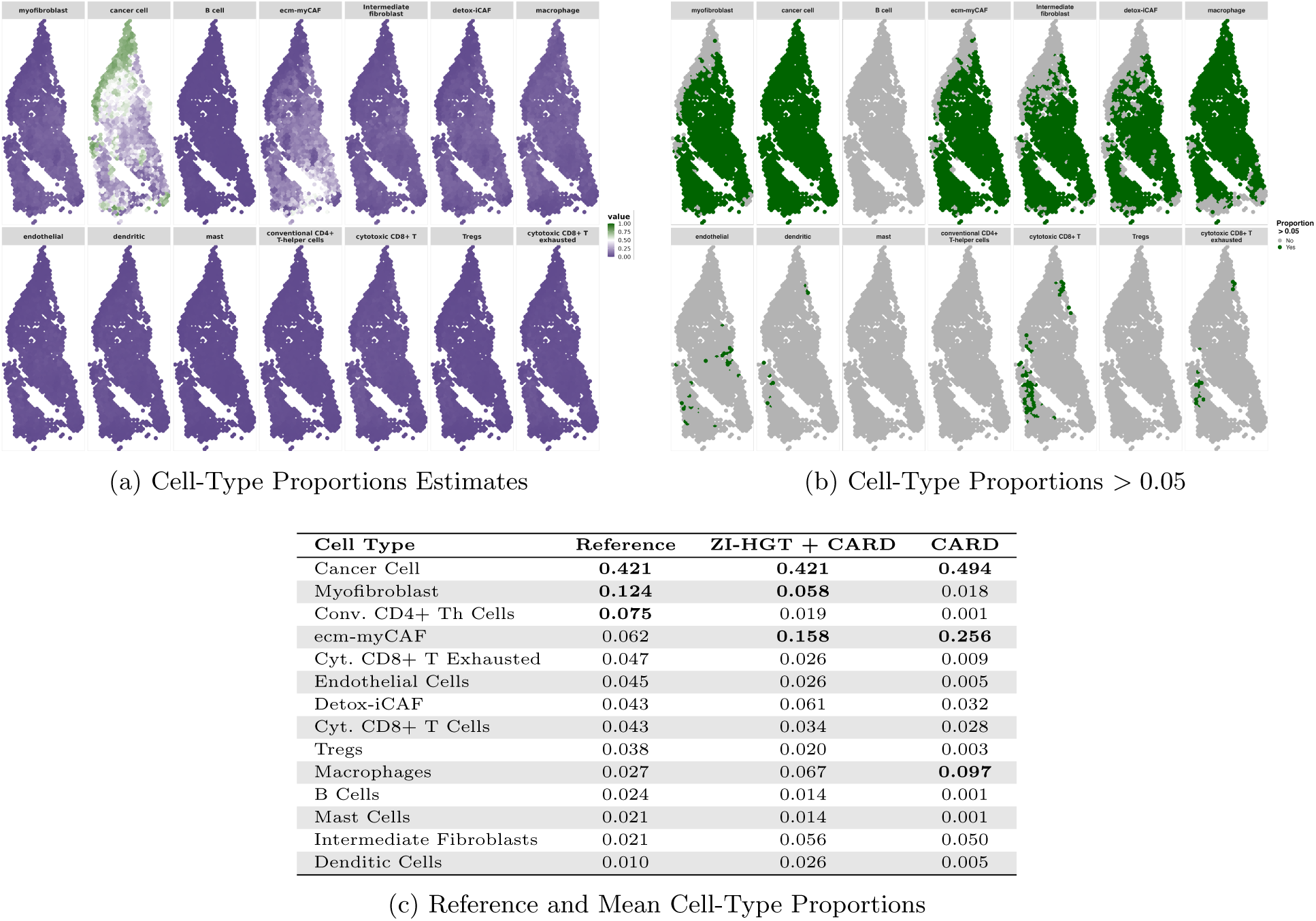
OSCC Sample 1 ZI-HGT + CARD Cell-Type Proportions Estimates. a) Estimated cell-type proportions for all 14 cell types. b) Binary maps showing locations where cell-type proportions are *>* 0.05. c) Reference cell-type proportions from scRNA-seq and average estimated proportions across all locations using ZI-HGT + CARD and CARD. Values in bold indicate the three largest proportions for each method.

## 6 Real Data Results

We applied the ZI-HGT + CARD to the OSCC data (Arora et al., 2023) introduced in Section 2. On all samples, we set the number of ZI-HGT iterations *C* = 100 and used the previously discussed HNSCC scRNA-seq data (GSE103322) as the deconvolution reference. We focus here on Sample 1 and present results for the Samples 2-12 in Web Figures 8-41.

OSCC Sample 1 contains spatially resolved gene expression for 15,844 transcripts across 1,131 locations. We analyze estimated proportions for all 14 cell types at each location in Figure 5 (ZI-HGT + CARD) and Web Figure 8 (CARD), displaying both proportion estimates and binary plots with coloring applied only when estimated proportions exceed 0.05. We selected this threshold as 10x Visium ST data contains approximately 10 cells per location, and therefore proportions above 0.05 reasonably indicate that at least one cell of that type is present. Such presence or absence information is clinically valuable, as the presence of tumor-infiltrating immune cells (T cells, B cells, etc.) can predict OSCC patient survival (Hadler-Olsen and Wirsing, 2019). In practice, each cell type is not present at each location. Determining which cell types are absent is challenging as zero gene expression of marker genes can arise from multiple mechanisms: actual cell type absence, technical artifacts, or transcriptionally quiet cells due to environmental factors. This mechanistic uncertainty is reflected in the uncertainty of the low cell-type proportions estimates.

The ZI-HGT + CARD demonstrates marked improvement in identifying cancer-associated fibroblasts (ecm-myCAFs and detox-iCAFs) and normal fibroblasts (myofibroblasts and intermediate fibroblasts) across the tumor microenvironment (TME) compared to CARD alone. Normal fibroblasts can suppress tumor growth but can transform into cancer-associated fibroblasts during tumorigenesis (Maia et al., 2021). While both fibroblast types distribute throughout the TME, cancer-associated fibroblasts colocalize more closely with tumor cells (Maia et al., 2021)—a pattern clearly illustrated through the application of ZI-HGT + CARD (Figure 5a) but less evident with CARD alone (Web Figure 8a). Since cancer-associated fibroblasts are potential treatment targets (Maia et al., 2021), better determining their locations in the TME could improve targeted therapeutics.

Assessing the precision of deconvoluted cell-type proportions in real datasets is challenging, as the ground truth is unknown. A common metric calculates Pearson’s correlation between mean estimated tissue-wide cell-type proportions and corresponding scRNA-seq proportions (Ma and Zhou, 2022; Li et al., 2023). For OSCC Sample 1, the correlation coefficient was 0.93 for the ZI-HGT + CARD vs. 0.85 for CARD alone (Figure 5). Across all 12 samples, the ZI-HGT + CARD estimates showed stronger correlation with the scRNA-seq reference proportions for 10 samples, with notable differences in complex samples (1, 3, 4, and 5). While 42.1% of the cells in the scRNA-seq dataset are cancer cells (Puram et al., 2017), CARD alone estimates 90.0% cancer cells across all locations. The ZI-HGT + CARD also overestimates the overall proportion of cancer cells, but at a notably better 79.5%. Using the ZI-HGT clearly reduces CARD’s substantial overestimation of dominant cell types.

Beyond CARD’s capabilities, the ZI-HGT naturally quantifies uncertainty in cell-type proportion estimates through pointwise Bayesian credible intervals (Figure 6). For each cell type and location, we approximate the variance using the iterated variance formula and a second order Taylor series approximation, then construct pointwise credible intervals as the estimated proportion ±1.96 standard deviations.

**Figure 6:**
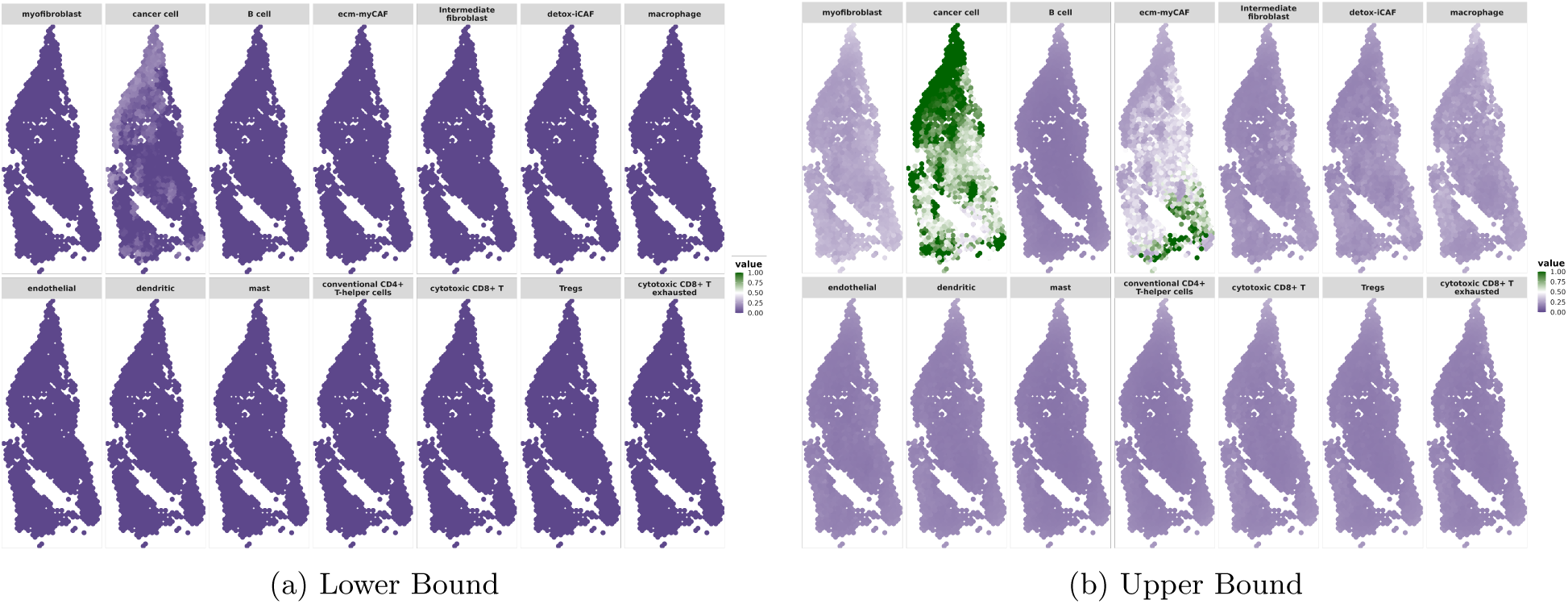
OSCC Sample 1 ZI-HGT + CARD Cell-Type Proportions Pointwise 95% Credible Intervals. a) Lower 95% bound. b) Upper 95% bound.

## 7 Conclusions

By integrating hierarchical generalized transformation models (Bradley et al., 2020; Bradley, 2022; Bradley et al., 2023) with Bayesian zero-inflation methods (Greene, 1994) and CARD cell-type deconvolution (Ma and Zhou, 2022), we address CARD’s normality assumption and incorporate uncertainty in analyzing OSCC spatial transcriptomics data (Arora et al., 2023). The ZI-HGT is broadly applicable and can serve as an auxiliary technique for future ST analysis methods that assume normality.

The combination of the ZI-HGT and CARD yielded compelling insights into the spatial distribution of cell types within the OSCC TME. The ZI-HGT + CARD exhibited superior accuracy compared to CARD alone or with a naive transformation, enabling identification of diverse fibroblast populations—critical members of the TME affecting tumor growth and immunosuppression. A notable strength is the ZI-HGT + CARD’s natural uncertainty quantification through the construction of pointwise Bayesian credible intervals without unrealistic distributional assumptions. UQ is rarely incorporated into ST analysis, and we anticipate numerous applications the ZI-HGT’s UQ framework. Researchers can use these credible intervals to assess the reliability of cell-type proportion estimates and apply pointwise Bayesian hypothesis tests to determine cell type presence at spatial locations.

Numerous methodological research directions remain open. New ST data capture techniques emerge annually, and adaptations of the ZI-HGT + CARD could extend to 3-dimensional single-cell resolution ST data. Approaches downstream of cell-type deconvolution, such as the cell-type-specific differential gene expression method C-SIDE (Cable et al., 2022), could be integrated into deep model structures with the ZI-HGT and CARD. Furthermore, techniques beyond cell-type deconvolution may benefit from the ZI-HGT, such as super-resolution gene expression prediction with iSTAR (Zhang et al., 2024). These promising ideas require substantial development and are reserved for future work. From a biological perspective, spatial patterns of estimated cell-type proportions warrant further exploration through collaboration with domain experts to fully realize these findings’ potential.

We conclude by discussing potential limitations of the ZI-HGT + CARD. First, the ZI-HGT’s transformations add a small amount of unavoidable noise to the ST data that could obstruct biological signal (Web Figure 45). This can be exacerbated when the sequencing depth is abnormally low (Web Figure 48). However, the ZI-HGT simultaneously reduces noise from CARD mis-specifying zero-inflated count-valued data as normal, yielding net improvement in deconvolution accuracy. The transformations may bias the data, as the high zero-inflation in the raw data results in insufficient “ground truth” to compare to. Thus, we used downstream deconvolution accuracy as a proxy to measure the effects of potential bias. Additionally, the limited integration of the ZI-HGT and CARD may be considered a weakness. CARD itself could potentially be altered to better align with the ZI-HGT and the data. CARD’s conditional autoregressive (CAR) spatial structure may be too simple, particularly for non-stationary data where nearest neighbor adjacencies are unsuitable. A more flexible spatial basis function expansion approach may prove superior, though this would require further exploration of the spatial structure of the ST data. Conversely, the independence of the ZI-HGT and CARD can be considered a strength. Alternative methods, such as the aforementioned iSTAR (Zhang et al., 2024), could replace CARD in the ZI-HGT + CARD model structure for tasks beyond cell-type deconvolution. We posit that the ZI-HGT provides a versatile foundation for future model development, enabling researchers to circumvent issues of zero-inflation and ties in ST data and simply assume normality.

## Acknowledgments

We thank the editor, associate editor, and two anonymous referees for their constructive suggestions and valuable feedback on our manuscript.

## Supplementary Material

Web Appendices, Tables, and Figures referenced in Sections 1-7 are available with this paper at the Biometrics website on Oxford Academic. All code used to generate the figures and results shown in this manuscript is available in the Supplementary Material.

## Data Availability

The data used in this paper to illustrate our findings are made available by Arora et al. (2023) at https://doi.org/10.6084/m9.figshare.20304456.v1 and Puram et al. (2017) at https://www.ncbi.nlm.nih.gov/geo/query/acc.cgi?acc=GSE103322.

