## Supplementary for "A Zero-Inflated Hierarchical Generalized Transformation Model to Address Non-Normality in Spatially-Informed Cell-Type Deconvolution"

Biometrics 000, 000–000 DOI: 000
000 0000

### Supplmentary Material for A Zero-Inflated Hierarchical Generalized Transformation Model to Address Non-Normality in Spatially-Informed Cell-Type Deconvolution by Hunter J. Melton, Jonathan R. Bradley, and Chong Wu

This paper has been submitted for consideration for publication in *Biometrics*

#### Web Appendix A: Data Collection and Preparation

*ST Data Preparation*

We analyze oral squamous cell carcinoma spatial transcriptomic (ST) data (Arora et al., 2023), consisting of 12 samples surgically resected from 10 patients' oral cavities. Post-surgery, the tissues were embedded in optimal cutting temperature compound and stored at -80°C. $10\mu m$ sections were obtained using a cryostat, hematoxylin and eosin (H&E) stained, and utilized for spatial transcriptomics. A pathologist annotated the sections into nine tissue layers: squamous cell carcinoma (SCC), lymphocyte-positive stroma, lymphocyte-negative stroma, normal mucosa, glandular stroma, muscle, keratin, and artery/vein, and an artifact layer. After determining optimal permeabilization time, ST was conducted using the 10x Genomics Visium platform, which samples approximately 10 cells per location. Across all samples, reads were obtained for an average of $16,763$ unique transcripts over $2,073$ spatial locations.

##### 0.1 *scRNA-seq Data Preparation*

In addition to the ST data, our analysis incorporates a reference single-cell RNA-sequencing (scRNA-seq) dataset from head and neck squamous cell carcinoma (HNSCC) (Puram et al., 2017) (GSE103322), encompassing 5,902 cells from 18 patients after quality control (QC). The scRNA-seq data were processed following Arora et al. (2023), involving normalization using 'SCTransform', dimension reduction, and Louvain clustering. Cells were initially clustered based on global expression patterns (again following Arora et al. (2023)) into eight clusters: cancer cells, T cells, B cells, macrophages, dendritic cells, mast cells, endothelial cells, and fibroblasts. T cells were further subcategorized into conventional CD4+ T-helper cells, regulatory T cells (Tregs), cytotoxic CD8+ T cells, and exhausted cytotoxic CD8+ T cells. Fibroblasts were further subcategorized into myofibroblasts, intermediate fibroblasts, extracellular matrix myofibroblastic cancer-associated fibroblasts (ecm-myCAFs), and detoxification pathway inflammatory cancer-associated fibroblasts (detox-iCAFs). Each of these cell types is discussed in more detail below.

First, the cancer cells are malignant, overproduced squamous cells that make up the tumor. While tumors may exhibit distinct oncogenic signaling pathways, the majority of OSCC cancer cells express epidermal growth factor receptors (*EGFR*), which induce OSCC cell proliferation, metastasis, invasion, and resistance to apoptosis (Tan et al., 2023). Many TMEs, including those surrounding OSCC, are marked by a diverse set of colocalizing fibroblasts. Myofibroblasts, marked by the expression of alpha smooth muscle actin (*ACTA2*) and myosin light chain proteins (*MYLK*, *MYL9*), have long been established in TMEs and play a major role in wound contracture and healing (Puram et al., 2017; Rockey et al., 2013). Intermediate, or resting, fibroblasts were depleted for markers indicating myofibroblasts or cancer-associated fibroblasts. The cancer-associated fibroblasts (CAFs), which spur tumor growth, angiogenesis, inflammation, and metastasis (Madar et al., 2013), were subdivided into two groups: extracellular matrix-associated myofibroblastic cancer-associated fibroblasts (ecm-myCAFs) and detoxification pathway inflammatory cancer-associated fibroblasts (detox-iCAFs). The subgroups were identified by marker genes *LRRC15* and *GBJ2* for ecm-MYCAFs and *ADH1B* and *GPX3* for detox-iCAFs (Arora et al., 2023).

T cells were divided into four subgroups: regulatory T cells ($T_{regs}$), conventional CD4$^+$ T-helper cells, cytotoxic CD8$^+$ T cells, and exhausted cytotoxic CD8$^+$ T cells. $T_{regs}$, typically identified by *FOXP3*, primarily act to suppress immune response (Kondělková¡ et al., 2010). Elevated levels of circulating $T_{regs}$ have been previously noted in lung, breast, pancreas, and prostate cancer, and they are postulated to facilitate tumorigenesis through immunosuppression (Wang et al., 2020). Conventional CD4$^+$ T-helper cells, marked by *CCR7* and *CCF7*, modulate the adaptive immune system by activating other immune cells such as cytotoxic CD8$^+$ T cells, B cells, and macrophages (Corthay, 2009). Cytotoxic CD8$^+$ T cells play a major role in immune defense against tumors and pathogens. Marked by *CD8* and *GZMA* and a lack of co-inhibitory receptors *PD1* and *CTLA4* (Puram et al., 2017), a high volume of cytotoxic $CD8^+$ T cells in squamous cell carcinoma is typically associated with positive clinical outcomes (Shimizu et al., 2019). Lastly, exhausted cytotoxic $CD8^+$ T cells are a persistent antigen stimulation-induced (as often occurs in tumors) differentiated state of cytotoxic $CD8^+$ T cells that is characterized by a loss of effector function and marked by the expression of *PD1* and *CTLA4* (Jiang et al., 2020).

B cells play a complex role in tumorigenesis. While B-cell depletion has been associated with the development of secondary malignancies, some studies indicate a pro-tumorigenic effect of B cells in squamous carcinomas (Gavrielatou et al., 2021). Mast cells, found in epithelial and mucosal tissues, function as part of both the adaptive and innate immune systems (Krystel-Whittemore et al., 2015). An overabundance of mast cells has been observed in OSCC (Kalra et al., 2012), as mast cells may drive tumor progression through angiogenesis and extracellular matrix degradation (Gudiseva et al., 2017). Macrophages are a form of white blood cell in the innate immune system that often become cancer-associated in the TME and promote tumor progression and metastasis (Xue et al., 2022). Tumor-associated macrophages (TAMs) gather at the leading edge of the tumor and may induce Epithelial-Mesenchymal Transition (EMT) in cancer cells, thereby worsening malignancy (Hu et al., 2016). Dendritic cells present antigens that drive pathogen response. As such, it is believed that improperly signaling dendritic cells lead to tumor progression (Dumitru et al., 2022). Endothelial cells line blood vessels, and tumor-derived vasculature, including endothelial cells, carries blood to the tumor (Yao and Zeng, 2023). Tumor-derived endothelial cells are typically marked by the lack of von Willebrand factor (vWF) (Naschberger et al., 2011).

**Web Appendix B: Simulation Study Design**

The simulation study primarily focuses on OSCC Sample 2, as we believe it to be fairly representative of the majority of the samples, though we ensure that the choice of sample has little impact (Web Figure 2). Sample 2 includes data from $1,749$ locations and $15,624$ transcripts.

Following convention, we begin by identifying the tissue layers and corresponding dominant cell types for each of these locations according to pathologist annotations (Arora et al., 2023) (see Web Table 14 for a list of pathologist-annotated layers and corresponding dominant cell types). At this point, it is typical to use the matching scRNA-seq data (Puram et al., 2017) to build the simulated ST data; however, after doing so, we found that the median sparsity level of the simulated ST was approximately 52%, drastically different than the 90% sparsity for sample 2 and 87-91% across all 12 samples. To reconcile the simulated ST data with the real OSCC data, we chose to generate sparser scRNA-seq data with SPARSim (Baruzzo et al., 2020) to subsequently build sparser simulated ST data.

**Generating scRNA-seq data:** SPARSim employs a two-step Gamma-Multivariate Hypergeometric mixed model for generating simulated scRNA-seq data. In the first step, the true scRNA-seq data $\mathbf{W}$, an $N \times M$ matrix with rows corresponding to transcripts, or genes, and columns corresponding to individual cells, is simulated from a Gamma distribution. Consider a specific cell type $ct$, and let there be $M_{ct}$ such cells in the dataset. Then, let $W_{i,j}$ be a random variable representing the expression level of gene $i$ in cell $j$ ($i = 1, ..., N;\ j = 1, ..., M_{ct}$). For any gene $i$,

$$W_{i,j} \sim Gamma\left(\text{shape} = \frac{1}{\Phi_{i,ct}},\ \text{scale} = Z_{i,ct} \cdot \Phi_{i,ct}\right), \tag{1}$$

where $Z_{i,ct}$ is the "average" intensity, or expression level of gene i in cell type $ct$, and $\Phi_{i,ct}$ describes the biological variability in the expression level of gene $i$ in cell type $ct$. This procedure iterates through all genes $i$ and then all cell types $ct$. Then, in the second step, the observed scRNA-seq data $\mathbf{Y}$, also an $N \times M$ matrix with rows corresponding to transcripts and columns corresponding to individual cells, is simulated from a Multivariate Hypergeometric distribution. This step attempts to model the technical variability of observed scRNA-seq data, as no data capture process can perfectly record the true scRNA-seq data. For any cell $j$,

$$\mathbf{Y}_j \sim \text{Multivariate Hypergeometric}(n = L_j, m = \mathbf{W}_j), \tag{2}$$

where $L_j$ is the library size, and $\mathbf{W}_j$ is the true scRNA-seq data to sample from. Note that the library size is the total number of gene expression reads included in the "observed" data $\mathbf{Y}_j$ for cell $j$.

The three SPARSim hyperparameters, $Z_{i,ct}$, $\Phi_{i,ct}$, and $L_j$, are originally estimated from the real scRNA-seq data (Puram et al., 2017). The intensity $Z_{i,ct}$ is the average value for the scran-normalized (Lun et al., 2016) expression read counts for gene $i$ and cell type $ct$. The biological variability parameter $\Phi_{i,ct}$ is estimated using the "*estimateGLMTagwiseDisp*" function in the *edgeR* package (Robinson et al., 2010). The library size $L_j$ is computed by summing the raw counts for each cell. To alter the sparsity of the simulated scRNA-seq data, and therefore the sparsity of the downstream simulated ST data, we estimated the three hyperparameters with the real scRNA-seq data (Puram et al., 2017) and then re-scaled the biological variability $\Phi_{i,ct}$ and library size $L_j$. For each set of scaled hyperparameters, we generated 10 simulated scRNA-seq datasets. Each simulated scRNA-seq dataset was split into two equal parts; the first was used to simulated the ST data as described below, and the second was used as the scRNA-seq reference dataset for the cell-type deconvolution.

**Generating ST data:** We construct the simulated ST data following the same scaffolding as CARD (Ma and Zhou, 2022), with a crucial modification: our simulations are based on both the real OSCC ST data (Arora et al., 2023) and the simulated scRNA-seq data. This modification was specifically designed to ensure closer alignment with the characteristics of the real-world datasets, such as the sparsity. Although methods to directly simulate ST data exist, such as SRTsim (Zhu et al., 2023), we chose to build the simulated ST data with simulated scRNA-seq data so that a "ground truth" set of cell-type proportions would exist. We divided the OSCC samples into layers according to pathologist annotations and assigned each layer a dominant cell type: endothelial cells for veins and arteries, intermediate fibroblasts for glandular stroma, ecm-myCAFs for lymphocyte-negative stroma, cytotoxic CD8+ T cells for lymphocyte-positive stroma, myofibroblasts for muscles, macrophages for non-cancerous mucosa, and cancer cells for squamous cell carcinoma. The simulations proceeded as follows: for a desired OSCC sample (Sample 2 was used for most analyses), a simulated scRNA-seq dataset was selected, enabling control over the sparsity of the simulated ST data. The simulated scRNA-seq data was divided into two parts: one for constructing the simulated ST data and the other for use in the cell-type deconvolution process. For each tissue layer in the sample, the number of colocalizing cell types with the dominant cell type was sampled from a discrete uniform distribution with a minimum of 1 and a maximum of 13. Next, the cell type proportions for each location within that tissue layer were determined by simulating from a Dirichlet distribution with concentration set to one. The largest proportion was assigned to the dominant cell type, and the proportions of other colocalizing cell types were randomly assigned. Ten cells were allocated to each location, with the counts for each cell type being ten times the sampled cell type proportions, rounded to the nearest whole number. Cells were sampled without replacement from the first split of the simulated scRNA-seq data, and gene expression levels were aggregated across the ten cells to determine the gene expression at each location. Finally, ten simulated scRNA-seq datasets were generated for each set of SPARSim (Baruzzo et al., 2020) hyperparameters. From each of these datasets, ten sets of ST data were simulated, resulting in a total of 100 sets of simulated ST data for each set of hyperparameters.

**Simulation settings and evaluation criteria:** In our simulation study, we explored various settings, encompassing different sparsity levels in the simulated spatial transcriptomics (ST) data (Figure 3), simulations based on diverse OSCC samples (Web Figure 2), and varying numbers of cell types in the ST data (Web Figure 1). Additionally, we compared the ZI-HGT + CARD to other cell-type deconvolution methods, including SPOTlight (Elosua-Bayes et al., 2021), SpatialDecon (Danaher et al., 2022), STdeconvolve (Miller et al., 2022), as well as CARD with a simple deterministic transformation and the HGT + CARD (without considering zero-inflation). We note that STdeconvolve (Miller et al., 2022) is a scRNA-seq reference-free method, so cell types are annotated post-deconvolution. To best compare with the other methods, we constructed 14 clusters (to match the true number of cell types) with STdeconvolve and deconvoluted the simulated ST data. Then, to map clusters back to specific cell types, we examined the correlation between the gene expression patterns of the clusters and the gene expression patterns of the known cell types in the scRNA-seq reference dataset. Beginning with the largest cluster, cell types are annotated by matching the cluster to the cell-type with the highest correlation in gene expression.

Our primary focus is on comparing the ZI-HGT + CARD to CARD alone, as CARD is the current state-of-the-art method and performed best among all evaluated methods. We primarily focused on OSCC Sample 2 because it is representative of the majority of the samples. Sparsity levels in the simulated ST data were controlled by multiplying the SPARSim hyperparameters by specific factors. Specifically, we generated the three SPARSim hyperparameters using the real scRNA-seq data (Puram et al., 2017). Through iterative adjustments, we found that multiplying the library size $L_j$ by 0.05 and the biological variability $\Phi_{i,ct}$ by 50 yielded the most realistic data in terms of sparsity and structure; these values were used in all simulations unless otherwise specified. We refer to these factors as the "library factor" and "$\Phi$ factor", respectively. Across 100 simulated ST datasets using these settings, sparsity levels ranged between 86% and 92%, closely aligning with the observed sparsity levels in the real OSCC data (Web Table 1).

We benchmarked the performance of the ZI-HGT + CARD vs. CARD alone in a variety of scenarios. First, we examined the impact of the level of sparsity in the data (Figure 3). We next ensured that the ZI-HGT's improvement was consistent in simulations with differing numbers of cell types (Web Figure 1) and using the 12 distinct OSCC samples as the basis for the simulated datasets (Web Figure 2). We explored the effect of increasing the number of nonzero ties in the simulated ST data (Web Figure 47). Increasing the number of nonzero ties was achieved by artificially raising the expression of a randomly selected subset of 20% of all genes in the simulated scRNA-seq data by 1, 2, or 3 for all simulated single-cells, which then propagates into the simulated ST data. Finally, we determined the impact of altering the sequencing depth of the simulated data (Web Figure 48), which was achieved by changing the number of simulated single cells allocated to each location in the simulated ST data.

In all simulations, we set $C$, the number of ZI-HGT iterations, to 100. To assess accuracy, we measured the root mean square error (RMSE) for the estimated cell-type proportions, given by

$$\text{RMSE} = \sqrt{\frac{1}{NK}\sum_{i=1}^{N}\sum_{k=1}^{K}\left(V_{i,k} - \hat{V}_{i,k}\right)^2}, \tag{3}$$

where $N$ is the number of spatial locations and $K$ is the number of cell types. We then compared the ZI-HGT + CARD to the original CARD method by determining the percent RMSE improvement over CARD, calculated by $100\% \times \left(1 - \text{RMSE}_{\text{ZI-HGT+CARD}}/\text{RMSE}_{\text{CARD}}\right)$.

*CARD Simulation Study Design*

We applied the ZI-HGT + CARD to an additionally simulation study mimicking the setup of CARD's simulations exactly. The design of this simulation study is quite similar to ours, though it is based on mouse olfactory bulb ST (Ståhl et al., 2016) and scRNA-seq datasets (Zeisel et al., 2018). The simulated ST data is much less sparse than the real ST data for this study: the median sparsity level was 68.6% in these simulated datasets vs. 88.5% in the real OSCC ST datasets.

The scRNA-seq reference comprises gene expression read counts for 20,515 cells across over six cell types: neurons, astrocytes, oligodendrocytes, vascular cells, immune cells, and ependymal cells. The scRNA-seq data is split into two pieces: one to construct the simulated ST data and the other to conduct cell-type deconvolution. The ST dataset contains 260 locations and three anatomic regions: the granule cell layer, the mitral cell layer, and the nerve layer. Each anatomic region was assigned a dominant cell type: oligodendrocytes for the granule cell layer, neurons for the mitral cell layer, and astrocytes for the nerve cell layer (Svensson et al., 2018). For additional robustness, Ma and Zhou (2022) considered simulations with added noise, which we follow. Let $p_n$ be the percentage of the 260 spatial locations that are "noisy". Data at noisy locations are generated identically to non-noisy locations, with the exception that all cell-type proportions are randomly assigned, so the dominant cell type no longer automatically receives the largest sampled proportion of cells. Noisiness percentages $p_n \in \{0\%, 20\%, 40\%, 60\%\}$ are considered.

Five simulation scenarios are considered: 1) The correct scRNA-seq data functions as the cell-type deconvolution reference dataset. 2) One cell-type is removed from the scRNA-seq reference dataset. 3) One cell-type (blood cells) is added to the scRNA-seq reference dataset. 4) Two cell-types are merged in the scRNA-seq reference dataset. 5) A mismatched scRNA-seq reference dataset from a different sequencing platform, mouse brain tissue, and mouse age is used. The mismatched reference comes from a different platform, a custom microwell-seq + Drop-seq approach (Mizrak et al., 2019) than the standard 10X Chromium platform that was used to generate the matched scRNA-seq reference (Zeisel et al., 2018). This is critical, as the simulated ST data were generated from an independent split of the matched scRNA-seq reference (Ma and Zhou, 2022), so platform-specific biases may lead to differences in gene expression patterns. Even more importantly, the ST data and scRNA-seq reference were generated from the olfactory bulb from adult mice. The mismatched reference was generated from the ventricular-subventricular zone (V-SVZ) of young adult mice (Mizrak et al., 2019). The V-SVZ is a major germinal zone of the mouse brain. While it produces olfactory bulb interneurons, among other cells, gene expression patterns are expected to be notably different than in the olfactory bulb (Lim and Alvarez-Buylla, 2016). The degree to which the ST data and scRNA-seq reference are mismatched in this scenario is uncommon–different scRNA-seq platforms, different studies, different ages of the mice, and different tissues (though both are in the brain). Since the ZI-HGT transforms the ST data by adding noise, the noise from the highly mismatched scRNA-seq reference is amplified, unsurprisingly leading to diminished accuracy.

Beyond CARD's simulation construction, we also alter the sequencing depth of the simulated ST data. The sequencing depth of a platform is defined as the total number of read counts across the entire tissue. As comparators for applying the ZI-HGT to CARD, we also apply CARD to simulated datasets with total sequencing depth $-10\%$ and $+10\%$ as compared to the original simulated datasets. We achieve increased and decreased sequencing depth by decreasing and increasing the number of cells per location in the simulation to nine and eleven, respectively.

#### Web Appendix C: Bayesian Hierarchical Model and Transformation Posterior Derivation

Below, we present the Bayesian hierarchical model (BHM) for the ZI-HGT + CARD. We believe it is simplest to individually display the ZI-HGT and CARD hierarchical models then their multiplication that results in the full model for the combined method. Let $f$ represent a generic probability density or mass function and $I(\cdot)$ represent the indicator function. Let $f(X|\theta) = \text{dist}(\theta)$ mean that $X|\theta$ follows the noted distribution with parameter $\theta$.

ZI-HGT BHM:

$$\prod_{i,j}\left[\left\{f\left(X_{i,j}\middle|H_{i,j},X_{i,j}^{[0]}=0,X_{i,j}^{(B)}\right)\ f\left(H_{i,j}\middle|X_{i,j}^{[0]}=0,X_{i,j}^{(B)}\right)\ f\left(X_{i,j}^{[0]}=0\middle|X_{i,j}^{(B)}\right)\ f\left(g\left(X_{i,j}^{(B)}\right)\right)\right\}^{I\left(X_{i,j}^{[0]}=0\right)}\right.$$
$$\left.\times\left\{f\left(X_{i,j}\middle|H_{i,j},X_{i,j}^{[0]}=1\right)\ f\left(H_{i,j}\middle|X_{i,j}^{[0]}=1\right)\right\}^{I\left(X_{i,j}^{[0]}=1\right)}\right];\quad i=1,...,N,\ j=1,...,G, \quad (4)$$

where:

- $f\left(X_{i,j}\middle|H_{i,j},X_{i,j}^{[0]}=0,X_{i,j}^{(B)}\right) \;=\; \text{Point Mass}(0)$
- $f\left(H_{i,j}\middle|X_{i,j}^{[0]}=0,X_{i,j}^{(B)}\right) \;=\; \text{Point Mass}\left(X_{i,j}^{(B)}\right)$
- $f\left(X_{i,j}^{[0]}\middle|X_{i,j}^{(B)}\right) \;=\; \text{Bernoulli}\left(g\left(X_{i,j}^{(B)}\right)\right)$
- $f\left(g\left(X_{i,j}^{(B)}\right)\right) \;=\; \text{Beta}(\alpha_0,\kappa_0)$
- $g\left(X_{i,j}^{(B)}\right) = \frac{\exp\left(X_{i,j}^{(B)}\right)}{1+\exp\left(X_{i,j}^{(B)}\right)}$
- $f\left(X_{i,j}\middle|H_{i,j},X_{i,j}^{[0]}=1\right) \;=\; \text{TruncPoisson}\left(H_{i,j}\right)$
- $f\left(H_{i,j}\middle|X_{i,j}^{[0]}=1\right) \;\propto\; (1-\exp(-H_{i,j}))\,\text{Gamma}(\alpha_1,\alpha_1)$

We note that, in practice, we let $\alpha_0 \in \{1.2975, 1.5, 1.7475, 2.0625\}$, $\kappa_0 \in \{0.4325, 0.5, 0.5825, 0.6875\}$ ($\kappa_0$ is set equal to $\frac{\alpha_0}{3}$ to reduce the computational burden), and $\alpha_1 \in \{0.1, 0.2, 0.3, 0.4, 0.5\}$, then chose a single set of hyperparameters by minimizing the WAIC (Watanabe, 2010). The ZI-HGT can be sensitive to the choice of hyperparameters, as the transformed data must overfit the real data. We decided the range for $\alpha_0$ and $\kappa_0$ to ensure that the transformed zero-data clusters near to zero, as $\alpha_0$ near 1.5 and $\kappa_0$ near 0.5 accomplishes this. We selected the range for $\alpha_1$ to ensure that the transformed non-zero data does not shrink too close to zero, which would occur with larger values of $\alpha_1$. To reduce the computational burden in simulations, we restrict the hyperparameter range slightly to $\alpha_0 \in \{1.5, 1.7475, 2.0625\}$, $\kappa_0 \in \{0.5, 0.5825, 0.6875\}$, and $\alpha_1 \in \{0.1, 0.3, 0.5\}$. We note that choosing the hyperparameters via the WAIC resulted in similar performance to the oracle-chosen (to minimize RMSE) hyperparameters in simulations (Web Figure 2).

Then the inference on the cell-type proportions estimate $\mathbf{V}$ in CARD can be expressed as an empirical Bayesian hierarchical model (EBHM)

CARD EBHM:

$$f\Big(\mathbf{H}\Big|\mathbf{V}, \sigma_e^2\Big)\ f\Big(\sigma_e^2\Big) \prod_k \Big\{ f\Big(\mathbf{V}_k | b_k, \lambda_k, \hat{\phi}(\mathbf{H})\Big)\ f\big(b_k\big)\ f\big(\lambda_k\big) \Big\};$$

$$k = 1, ..., K, \quad (5)$$

where:

- $f\Big(\mathbf{H}\Big|\mathbf{V}, \sigma_e^2\Big) \;=\; N\Big(\mathbf{B}\mathbf{V}', \sigma_e^2 \mathrm{I}_{N\times G}\Big)$
- $f\Big(\sigma_e^2\Big) \;=\; 1$
- $f\Big(\mathbf{V}_k | b_k, \lambda_k, \hat{\phi}(\mathbf{H})\Big) \;=\; N\Big(b_k \times \mathbf{1}_N, \mathbf{\Sigma}_k\Big)$
- $f\Big(b_k\Big) \;=\; 1$
- $\mathbf{\Sigma}_k = \lambda_k \big(\mathbf{D} - \phi \mathbf{W}\big)^{-1}$
- $\mathbf{D} = \mathrm{diag}\big(w_{1+}, w_{2+}, ..., w_{N+}\big)$
- $w_{i+} = \sum_{j=1}^{G} w_{i,j}$
- $\mathbf{W} = \{w_{i,j}\}$; where $w_{i,j}$ comes from the Gaussian kernel function commonly used in a CAR model (Besag, 1974).
- A gridded search over $\hat{\phi}\big(\mathbf{H}\big) \in \{0.01, 0.1, 0.3, 0.5, 0.7, 0.9, 0.99\}$ is conducted in the CARD algorithm and $\hat{\phi}\big(\mathbf{H}\big)$ is chosen to maximize the log-likelihood.
- $f\Big(\lambda_k\Big) \;=\; \text{Inverse Gamma}\Big(1, \frac{N\times G}{2}\Big)$

As mentioned above, CARD is an empirical Bayesian method. The log posterior predictive distribution $\log P\left(\mathbf{V}, b_k, \lambda_k, \sigma_e^2 | \mathbf{B}, \mathbf{X}, \phi\right)$ is presented in Equation (9) in the supplementary material of Ma and Zhou (2022) (for the ZI-HGT + CARD, $\mathbf{X}$ is replaced by $\mathbf{H}$ and $\phi$ by $\phi(\mathbf{H})$ as shown above). Ma and Zhou (2022) then developed an algorithm that iterates through each of element of $\mathbf{V}$ and the hyperparameters $b_k, \lambda_k, \sigma_e^2$ to minimize the negative log likelihood. This procedure results in the point estimate of the cell type proportions $\widehat{\mathbf{V}}$.

Then the BHM for the combination of the ZI-HGT + CARD is given below:

ZI-HGT + CARD BHM:

$$\prod_{i,j}\Bigg[\Big\{f\Big(X_{i,j}\Big|H_{i,j}, X_{i,j}^{[0]}=0, X_{i,j}^{(B)}\Big)\ f\Big(H_{i,j}\Big|X_{i,j}^{[0]}=0, X_{i,j}^{(B)}\Big)\ f\Big(X_{i,j}^{[0]}=0\Big|X_{i,j}^{(B)}\Big)\ f\Big(g\Big(X_{i,j}^{(B)}\Big)\Big)\Big\}^{I\left(X_{i,j}^{[0]}=0\right)}$$
$$\times\Big\{f\Big(X_{i,j}\Big|H_{i,j}, X_{i,j}^{[0]}=1\Big)\ f\Big(H_{i,j}\Big|X_{i,j}^{[0]}=1\Big)\Big\}^{I\left(X_{i,j}^{[0]}=1\right)}\Bigg]$$
$$\times f\Big(\mathbf{H}\Big|\mathbf{V},\sigma_e^2\Big)\ f\Big(\sigma_e^2\Big)\prod_k\Big\{f\Big(\mathbf{V}_k|b_k,\lambda_k,\hat{\phi}(\mathbf{H})\Big)\ f\big(b_k\big)\ f\big(\lambda_k\big)\Big\}$$
$$\Bigg/\int\int\dots\int f\Big(\mathbf{H}\Big|\mathbf{V},\sigma_e^2\Big)\ f\Big(\sigma_e^2\Big)\prod_k\Big\{f\Big(\mathbf{V}_k|b_k,\lambda_k,\hat{\phi}(\mathbf{H})\Big)\ f\big(b_k\big)\ f\big(\lambda_k\big)\Big\}\, d\sigma_e^2 db_1\dots db_K d\lambda_1\dots d\lambda_K;$$
$$i=1,\dots,N,\ \ j=1,\dots,G,\ \ k=1,\dots,K. \quad (6)$$

where we have multiplied the ZI-HGT model by the predictive distribution

$$f(\mathbf{V},\boldsymbol{\theta}|\mathbf{H}) = \frac{f\Big(\mathbf{H}\Big|\mathbf{V},\sigma_e^2\Big)\ f\Big(\sigma_e^2\Big)\prod_k\Big\{f\Big(\mathbf{V}_k|b_k,\lambda_k,\hat{\phi}(\mathbf{H})\Big)\ f\big(b_k\big)\ f\big(\lambda_k\big)\Big\}}{\int\int\dots\int f\Big(\mathbf{H}\Big|\mathbf{V},\sigma_e^2\Big)\ f\Big(\sigma_e^2\Big)\prod_k\Big\{f\Big(\mathbf{V}_k|b_k,\lambda_k,\hat{\phi}(\mathbf{H})\Big)\ f\big(b_k\big)\ f\big(\lambda_k\big)\Big\}\, d\sigma_e^2 db_1\dots db_K d\lambda_1\dots d\lambda_K}, \quad (7)$$

where $\theta = \{\sigma_\epsilon^2, b_1, \dots, b_K, \lambda_1, \dots, \lambda_K\}$.

Below, we present the derivation of the transformation posterior for $H_{i,j}$ shown in Equations (10) and (11). First, consider the case where the original data $X_{i,j} > 0$. Per Bayes' rule,

$$f\left(H_{i,j}|X_{i,j}^{[0]}=1, X_{i,j}\right)$$
$$\propto f\left(X_{i,j}|X_{i,j}^{[0]}=1, H_{i,j}\right) f\left(H_{i,j}|X_{i,j}^{[0]}=1\right)$$
$$\propto \frac{H_{i,j}^{X_{i,j}}\exp\left(-H_{i,j}\right)}{X_{i,j}!\left(1-\exp\left(-H_{i,j}\right)\right)}\times\left(1-\exp\left(-H_{i,j}\right)\right)\frac{\alpha_1^{\alpha_1}H_{i,j}^{\alpha_1-1}\exp\left(-\alpha_1 H_{i,j}\right)}{\Gamma(\alpha_1)}$$
$$\propto H_{i,j}^{\alpha_1+X_{i,j}-1}\exp\left(-H_{i,j}(\alpha_1+1)\right); \quad i=1,\dots,N,\ \ j=1,\dots,G. \quad (8)$$

Clearly, this is the kernel of a Gamma distribution with shape parameter $\alpha_1 + X_{i,j}$ and rate parameter $\alpha_1 + 1$, so $H_{i,j} \big| X_{i,j}^{[0]} = 1, X_{i,j} \sim \text{Gamma}(\alpha_1 + X_{i,j}, \alpha_1 + 1)$ as shown in Equation (10). Now examine the case where the original data $X_{i,j} = 0$. Per Bayes' rule,

$$\begin{aligned} & f\left(H_{i,j} \big| X_{i,j}^{[0]} = 0, X_{i,j}\right) \\ & \qquad \propto f\left(X_{i,j} \big| X_{i,j}^{[0]} = 0, H_{i,j}, X_{i,j}^{(B)}\right) f\left(H_{i,j} \big| X_{i,j}^{[0]} = 0, X_{i,j}^{(B)}\right) f\left(X_{i,j}^{[0]} \big| X_{i,j}^{(B)}\right) f\left(X_{i,j}^{(B)}\right) \\ & \propto I\left(X_{i,j} = 0\right) I\left(H_{i,j} = X_{i,j}^{(B)}\right) g\left(X_{i,j}^{(B)}\right)^{0} \left(1 - g\left(X_{i,j}^{(B)}\right)\right)^{1-0} g\left(X_{i,j}^{(B)}\right)^{\alpha_0 - 1} \left(1 - g\left(X_{i,j}^{(B)}\right)\right)^{\kappa_0 - 1} \\ & \qquad \propto g\left(H_{i,j}\right)^{\alpha_0 - 1} \left(1 - g\left(H_{i,j}\right)\right)^{(\kappa_0 + 1) - 1}; \quad i = 1, ..., N, \; j = 1, ..., G, \end{aligned} \tag{9}$$

where the last line holds true as $H_{i,j}$ is necessarily equal to $X_{i,j}^{(B)}$. Clearly, this is the kernel of a beta distribution with parameters $\alpha_0$ and $\kappa_0+1$, so $g\left(H_{i,j}\right) \big| X_{i,j}^{[0]} = 0, X_{i,j} \sim \text{Beta}(\alpha_0, \kappa_0+1)$ as shown in Equation (11), and $g\left(H_{i,j}\right) = \frac{\exp(H_{i,j})}{1+\exp(H_{i,j})}$.

*Practical Considerations for Choosing Hyperparameters*

We recognize that, in practice, the selection of candidate values for the ZI-HGT hyperparameters $\alpha_0$ and $\alpha_1$ is important. This section provides practical guidance to users for making sensible choices given our experiences.

When using the ZI-HGT, we recommend beginning with a small set of hyperparameters, applying the ZI-HGT, and examining the WAIC for each candidate pair, as we did in the simulation study and real data analysis. Then, if there is a clear "best" choice that minimizes the WAIC, select that pair of hyperparameters and proceed. If there is no clear "best" choice using the WAIC, we recommend using a finer grid when searching for hyperparameters. For initial choices, we recommend letting $\alpha_0 \in \{1.2975, 1.5, 1.7475, 2.0625\}$, $\kappa_0 = \frac{\alpha_0}{3}$, and $\alpha_1 \in \{0.1, 0.2, 0.3, 0.4, 0.5\}$. The range for $\alpha_0$ and $\kappa_0$ ensures that the transformed zero-data clusters near to zero, and the range for $\alpha_1$ ensures that the transformed non-zero data does not shrink too close to zero (as would occur with larger values of $\alpha_1$). If there is no best choice, we suggest appending the candidate hyperparameter space with intermediate values and re-analyzing the data, choosing the best hyperparameter pair from this analysis.

#### Web Appendix D: Literature Review on Theoretical Support for HGT

In general, the HGT infuses noise into the observations, which breaks ties and leads to conjugacy and tractable computations. Of course, one can induce noise in several different ways (e.g., adding simple Gaussian noise with a small variance). However, there has been theoretical support for this particular way to inject noise (or transform data) presented in the literature, which we now review. In particular, this strategy to introduce noise can be seen as a parametric model for signal-to-noise cross covariances.

For many datasets it is expected that signal-to-noise cross-covariances may be present. For example, the variability associated with genes with low expression may be disproportionately influenced by measurement errors as there are potential drop-out effects and the signal-to-noise ratio is naturally smaller. Consequently, the value of the noise from measurement error depends on the value of the signal, suggesting that this covariance may be present in the dataset. In general, accounting for the presence of a covariance offers an opportunity to obtain more precise predictions in spatial and spatio-temporal contexts (Banerjee et al., 2014). Furthermore, Bradley et al. (2020) show that one obtains more precise predictions in mean squared error when accounting for signal-to-noise covariances. However, the presence of a signal-to-noise cross covariance introduces theoretical concerns of identifiability that need to be addressed to be able to formally account for signal-to-noise covariance in Bayesian and other likelihood based settings. In particular, not all parametric models for the cross-signal-to-noise covariance produce a likelihood that is identifiable. For discussion, consider Gaussian distributed data where

$$\mathbf{X} = \mathbf{B}\mathbf{V}' + \delta,$$

where $\boldsymbol{\theta}$ is a generic finite-dimensional real parameter vector and denote $\boldsymbol{\Sigma}_X = \text{Cov}(\mathbf{X}|\boldsymbol{\theta}), \boldsymbol{\Sigma}_{BV} = \text{Cov}(\mathbf{BV}'|\boldsymbol{\theta}), \boldsymbol{\Sigma}_\delta = \text{Cov}(\delta|\boldsymbol{\theta})$, and $\boldsymbol{\Sigma}_{BV,\delta} = \text{Cov}(\mathbf{BV}', \delta|\boldsymbol{\theta})$. To avoid identifiability issues in this setup one typically assumes $\boldsymbol{\Sigma}_\delta = \mathbf{D}_\delta$ is a diagonal matrix with positive diagonal entries and $\boldsymbol{\Sigma}_{BV,\delta}$ is a zero matrix. Bradley et al. (2020) call this specification the "standard assumption," which produces an identifiable likelihood, since

$$\boldsymbol{\Sigma}_X = \boldsymbol{\Sigma}_{BV} + \mathbf{D}_\delta, \tag{10}$$

contains cross diagonal elements in $\boldsymbol{\Sigma}_{BV}$ that do not compete with the cross-diagonal elements in $\mathbf{D}_\delta$ that are zero. Bradley et al. (2020) investigated whether or not a different specification of $\boldsymbol{\Sigma}_\delta$ and $\boldsymbol{\Sigma}_{BV,\delta}$ could produce the same identifiable form in (10). They show that such a parameterization exists and involves introducing two matrix valued parameters: $\boldsymbol{\Sigma}_{BV,H}$ and $\boldsymbol{\Sigma}_H$. In particular, Bradley et al. (2020) consider the following specification of $\boldsymbol{\Sigma}_\delta$ and $\boldsymbol{\Sigma}_{BV,\delta}$

$$\begin{aligned} \boldsymbol{\Sigma}_\delta &= \boldsymbol{\Sigma}_{BV} + \boldsymbol{\Sigma}_H - \boldsymbol{\Sigma}_{BV,H} - \boldsymbol{\Sigma}'_{BV,H} + \mathbf{D}_\delta, \\ \boldsymbol{\Sigma}_{BV,\delta} &= \boldsymbol{\Sigma}_{BV,H} - \boldsymbol{\Sigma}_{BV}, \end{aligned} \tag{11}$$

which is referred to as the "general assumption". It is immediate that $\boldsymbol{\Sigma}_{BV,\delta} = \boldsymbol{\Sigma}_{BV,H} - \boldsymbol{\Sigma}_{BV}$ is not necessarily a zero matrix. Additionally, when substituting $\boldsymbol{\Sigma}_\delta$ and $\boldsymbol{\Sigma}_{BV,\delta}$ as defined in the general assumption into the expression for $\boldsymbol{\Sigma}_X$ we obtain

$$\boldsymbol{\Sigma}_X = \boldsymbol{\Sigma}_H + \mathbf{D}_\delta,$$

which is the same identifiable form as in (10) where a positive definite matrix $\boldsymbol{\Sigma}_H$ has off-diagonal elements that do not compete in the likelihood with the off-diagonal elements of $\mathbf{D}_\delta$. The telescoping nature of (11) allows for potentially non-zero $\boldsymbol{\Sigma}_{BV,\delta}$ while preserving an identifiable form for $\text{Cov}(\mathbf{BV} + \delta|\boldsymbol{\theta})$. In Bradley et al. (2020), several special cases are considered, including $\boldsymbol{\Sigma}_{BV,H} = \boldsymbol{\Sigma}_H = \boldsymbol{\Sigma}_{BV}$, which reproduces the standard assumption. "Special Case 1" in Bradley et al. (2020) occurs when $\boldsymbol{\Sigma}_H \neq \boldsymbol{\Sigma}_{BV,H} = \boldsymbol{\Sigma}_{BV}$, where there is no cross-signal-to-noise covariance since $\boldsymbol{\Sigma}_{BV,\delta} = \boldsymbol{\Sigma}_{BV,H} - \boldsymbol{\Sigma}_{BV} = \boldsymbol{\Sigma}_{BV} - \boldsymbol{\Sigma}_{BV} = \mathbf{0}$ where

$\mathbf{0}$ is a zero matrix. Their "Special Case 2" occurs when when $\boldsymbol{\Sigma}_H = \boldsymbol{\Sigma}_{BV,H} \neq \boldsymbol{\Sigma}_{BV}$, which produces a non-zero cross-signal-to-noise covariance matrix $\boldsymbol{\Sigma}_{BV,\delta}$.

A common strategy to avoid working directly with covariance matrices is to instead condition on random effects (Banerjee et al., 2014). The same can be done in the context of the general assumption. In particular, the general assumption arises when assuming

$$\mathbf{X} = \mathbf{B}\mathbf{V}' + \delta$$
$$\delta = \mathbf{H}\text{–}\mathbf{B}\mathbf{V}' + \epsilon,$$

which upon substituting $\delta$ into $\mathbf{X} = \mathbf{B}\mathbf{V}' + \delta$ produces the identifiable form

$$\mathbf{X} = \mathbf{H} + \delta,$$

where $\mathbf{H}$ is mutually independent of $\delta$. However, signal-to-noise covariance can be present, since $\text{Cov}(\mathbf{B}\mathbf{V}', \mathbf{H} - \mathbf{B}\mathbf{V}' + \epsilon|\boldsymbol{\theta}) = \text{Cov}(\mathbf{B}\mathbf{V}', \mathbf{H}|\boldsymbol{\theta}) - \text{Cov}(\mathbf{B}\mathbf{V}'|\boldsymbol{\theta})$ may not be zero. Bradley et al. (2020) show that Special Case 1 occurs when defining the hierarchical model $f(\mathbf{H}|\mathbf{B}\mathbf{V}', \boldsymbol{\theta})$ where $\mathbf{B}\mathbf{V}'$ is treated as the conditional mean for the model for $\mathbf{H}$ (recall this is the case where no cross-signal-to-noise is present). Special Case 2 occurs when defining the density $f(\mathbf{B}\mathbf{V}'|\mathbf{H}, \boldsymbol{\theta})$ (or $f(\mathbf{V}, \boldsymbol{\theta}|\mathbf{H})$) which produces cross-signal-to-noise covariances. We use such a specification in our ZI-HGT + CARD model via Equation (7). The inclusion of a discrepancy term in non-Gaussian settings is fairly straightforward, and was developed in Bradley et al. (2023). To do this, one specifies a link function $g$ as is typical in generalized mixed effects models (in our case the logit for Benroulli data and identity for Poisson data), defining

$$g(E[\mathbf{X}|\mathbf{V}]) = \mathbf{B}\mathbf{V}' + \delta$$
$$\delta = \mathbf{H} - \mathbf{B}\mathbf{V}'$$

and then defining a model $f(\mathbf{V}, \boldsymbol{\theta}|\mathbf{H})$, which we define according to CARD in (7).

#### Web Appendix E: The ZI-HGT written in the form of a zero-inflated Poisson model

In Section 4.1, we note that the ZI-HGT can be written in the form of a zero-inflated Poisson model (Lambert, 1992) by marginalizing across $X_{i,j}^{[0]}$:

$$X_{i,j}\big|H_{i,j}, X_{i,j}^{(B)} \sim \pi^*_{i,j}\text{TruncPoisson}(H_{i,j}) + (1-\pi^*_{i,j})\text{Point Mass}(0). \tag{12}$$

We write the argument to demonstrate this below. Let $f$ represent a generic probability density or mass function and $I(\cdot)$ represent the indicator function, and let $f(X|\theta) \sim \text{dist}(\theta)$ mean that $X|\theta$ follows the noted distribution with parameter $\theta$. Then,

$$f\left(X_{i,j}, X_{i,j}^{[0]}\middle|H_{i,j}, X_{i,j}^{(B)}\right) = f\left(X_{i,j}\middle|X_{i,j}^{[0]}, H_{i,j}, X_{i,j}^{(B)}\right) f\left(X_{i,j}^{[0]}\middle|H_{i,j}, X_{i,j}^{(B)}\right), \tag{13}$$

where the equality comes from a simple conditioning argument. Marginalize across the two values of $X_{i,j}^{[0]}$, namely 0 and 1, to construct the following expression:

$$\begin{aligned} f\left(X_{i,j}\middle|X_{i,j}^{[0]}=1, H_{i,j}, X_{i,j}^{(B)}\right) f\left(X_{i,j}^{[0]}=1\middle|H_{i,j}, X_{i,j}^{(B)}\right) \\ + f\left(X_{i,j}\middle|X_{i,j}^{[0]}=0, H_{i,j}, X_{i,j}^{(B)}\right) f\left(X_{i,j}^{[0]}=0\middle|H_{i,j}, X_{i,j}^{(B)}\right). \end{aligned} \tag{14}$$

Recall that $f\left(X_{i,j}\middle|X_{i,j}^{[0]}=1, H_{i,j}, X_{i,j}^{(B)}\right) = \text{TruncPoisson}\left(H_{i,j}\right)$ and $f\left(X_{i,j}|X_{i,j}^{[0]}=0, H_{i,j}, X_{i,j}^{(B)}\right) = \text{Point Mass}\left(0\right)$. So, letting $\pi^*_{i,j} = f\left(X_{i,j}^{[0]}=1|H_{i,j}, X_{i,j}^{(B)}\right)$, we arrive back at the previously noted expression, which is of the form of a zero-inflated Poisson model as desired.

By a conditioning argument,

$$\pi^*_{i,j} = f\left(X_{i,j}^{[0]}=1|H_{i,j}, X_{i,j}^{(B)}\right) = \frac{f\left(X_{i,j}^{[0]}=1, H_{i,j}, X_{i,j}^{(B)}\right)}{f\left(H_{i,j}, X_{i,j}^{(B)}\right)}, \tag{15}$$

so we consider the distributions on the right-hand side of this equation in more detail. First,

$$f\left(X_{i,j}^{[0]}=1,H_{i,j},X_{i,j}^{(B)}\right)=f\left(H_{i,j}\middle|X_{i,j}^{[0]}=1,X_{i,j}^{(B)}\right)f\left(X_{i,j}^{[0]}=1\middle|X_{i,j}^{(B)}\right)f\left(X_{i,j}^{(B)}\right)$$
$$=(1-\exp(-H_{i,j}))\text{Gamma}(\alpha_1,\alpha_1)\frac{\exp\left(X_{i,j}^{(B)}\right)}{1+\exp\left(X_{i,j}^{(B)}\right)}\text{Beta}(\alpha_0,\kappa_0), \tag{16}$$

which follows from the conditional independence of $H_{i,j}$ and $X_{i,j}^{(B)}$ given $X_{i,j}^{[0]}=1$. Then, with a similar argument,

$$f\left(X_{i,j}^{[0]}=0,H_{i,j},X_{i,j}^{(B)}\right)=f\left(H_{i,j}\middle|X_{i,j}^{[0]}=0,X_{i,j}^{(B)}\right)f\left(X_{i,j}^{[0]}=0\middle|X_{i,j}^{(B)}\right)f\left(X_{i,j}^{(B)}\right)$$
$$=\text{Point Mass}(X_{i,j}^{(B)})\frac{1}{1+\exp\left(X_{i,j}^{(B)}\right)}\text{Beta}(\alpha_0,\kappa_0). \tag{17}$$

Finally, recalling that $\pi_{i,j}=\frac{\exp\left(X_{i,j}^{(B)}\right)}{1+\exp\left(X_{i,j}^{(B)}\right)}$,

$$f\left(H_{i,j},X_{i,j}^{(B)}\right)=\sum_{k=0}^{1}f\left(X_{i,j}^{[0]}=k,H_{i,j},X_{i,j}^{(B)}\right)$$
$$=\pi_{i,j}(1-\exp(H_{i,j}))\text{Gamma}(\alpha_1,\alpha_1)\text{Beta}(\alpha_0,\kappa_0)$$
$$+(1-\pi_{i,j})\text{Point Mass}(X_{i,j}^{(B)})\text{Beta}(\alpha_0,\kappa_0). \tag{18}$$

Thus, all of the expressions in the specification of $\pi_{i,j}^*$ are defined.

#### Web Appendix F: Uncertainty Quantification

Through the ZI-HGT, we construct pointwise credible intervals for all estimated cell-type proportions $\widehat{\mathbf{V}}$. Plainly, we estimate $\text{Var}(\widehat{\mathbf{V}}|\mathbf{X})$. A fully Bayesian approach using a Gibbs sampler could in principle estimate this quantity exactly, but would be computationally intensive and would largely undermine the efficiency of our conjugate posterior sampling strategy in the ZI-HGT + CARD framework. Instead, we adopt an approximation based on the iterated variance formula:

$$\text{Var}(\mathbf{V}|\mathbf{X}) \;=\; \text{Var}\big(E(\mathbf{V}|\mathbf{H},\mathbf{X})|\mathbf{X}\big) \;+\; E\big(\text{Var}(\mathbf{V}|\mathbf{H},\mathbf{X})|\mathbf{X}\big). \tag{19}$$

The first term on the right-hand side of Equation 19, $\text{Var}\big(E(\mathbf{V}|\mathbf{H},\mathbf{X})|\mathbf{X}\big)$, captures the uncertainty arising from the variability in $\mathbf{H}$. We approximate this via the empirical variance of the point estimates of the cell-type proportions, i.e., $\text{Var}\big(E(\mathbf{V}|\mathbf{H},\mathbf{X})|\mathbf{X}\big) \approx \text{Var}\big(\widehat{\mathbf{V}}(\mathbf{H})\big)$. The second term on the right-hand side of Equation 19, $E\big(\text{Var}(\mathbf{V}|\mathbf{H},\mathbf{X})|\mathbf{X}\big)$, represents the uncertainty of the cell-type proportions estimate from CARD. We use a second order Taylor series approximation of $\text{Var}(\mathbf{V}|\mathbf{H},\mathbf{X})$. In particular, this approximation of $\text{Var}(\mathbf{V}|\mathbf{H},\mathbf{X})$ is simply the Fisher information matrix associated with the CARD model, which is consistent with the existing theory (e.g., see Bernstein von Mises theorem (van der Vaart, 1998)). Therefore, $E\big(\text{Var}(\mathbf{V}|\mathbf{H},\mathbf{X})|\mathbf{X}\big) \approx \mathbf{I}^{-1}(\widehat{\mathbf{V}})$, where $\mathbf{I}(\cdot)$ denotes the Fisher information. Taken together, we have

$$\text{Var}(\mathbf{V}|\mathbf{X}) \;\approx\; \text{Var}\big(\widehat{\mathbf{V}}(\mathbf{H})\big) \;+\; \frac{1}{C}\sum_{c=1}^{C}\mathbf{I}^{-1}\big(\widehat{\mathbf{V}}(\mathbf{H}^{[c]})\big), \tag{20}$$

where $C$ is the number of generated posterior predictions of $\widehat{\mathbf{V}}$. We now derive an expression for $\mathbf{I}\big(\widehat{\mathbf{V}}(\mathbf{H}^{[c]})\big)$. From Equation (12) in the supplementary material of CARD (Ma and Zhou, 2022), the partial derivative of the negative log-likelihood $\mathbf{Q} = -\log\big(\mathbf{V},\lambda_k,\sigma_e^2,b_k|\mathbf{B},\mathbf{X},\phi,\alpha,\beta\big)$ with respect to the column vector $\mathbf{V}_k$ (the cell-type proportions for cell type $k$):

$$\frac{\partial\mathbf{Q}}{\partial\mathbf{V}_k} \;=\; \frac{1}{2\sigma_e^2}\left(-2\mathbf{X}'\mathbf{B}_k + 2\mathbf{V}_k{\mathbf{B}_k}'\mathbf{B}_k + 2\sum_{j\neq k}\mathbf{V}_j\mathbf{B}_j'\mathbf{B}_k\right) + \frac{1}{2\lambda_k}2\mathbf{L}\mathbf{V}_k - \frac{1}{\lambda_k}b_k\mathbf{L}\mathbf{1}_N, \tag{21}$$

where $\mathbf{L} = \mathbf{D} - \phi\mathbf{W}$ and all other variables are as described in Web Appendix C. Differentiating again with respect to $\mathbf{V}_k$, we obtain the observed Fisher information

$$\mathbf{I}\big(\widehat{\mathbf{V}}_k(\mathbf{H})\big) \;=\; \frac{\partial^2\mathbf{Q}}{\partial\mathbf{V}_k^2} \;=\; \frac{1}{\sigma_e^2}{\mathbf{B}_k}'\mathbf{B}_k + \frac{1}{\lambda_k}\mathbf{L}. \tag{22}$$

For a given cell type $k$ and spatial location $i$, the observed Fisher information is

$$\mathbf{I}\big(\widehat{V}_{i,k}(H)\big) = \frac{\partial^2 \mathbf{Q}}{\partial V_{i,k}^2} = \mathbf{e}_i{}' \left(\frac{1}{\sigma_e^2}\mathbf{B}_k{}'\mathbf{B}_k + \frac{1}{\lambda_k}\mathbf{L}\right)\mathbf{e}_i = \frac{\mathbf{B}_k{}'\mathbf{B}_k}{\sigma_e^2} + \frac{L_{i,i}}{\lambda_k}, \tag{23}$$

where $\mathbf{e}_i$ is the $i$-th unit vector. Thus,

$$E\big(\mathrm{Var}(V_{i,k}|\mathbf{H},\mathbf{X})|\mathbf{X}\big) \approx \mathbf{I}^{-1}(\widehat{V}_{i,k}) = \left[\frac{1}{\sigma_e^2}\mathbf{B}_k{}'\mathbf{B}_k + \frac{1}{\lambda_k}\mathbf{L}\right]^{-1}_{(i,i)}. \tag{24}$$

We note that, in practice, CARD estimates $\lambda_k$, $\sigma_e^2$, and $\phi$ with $\widehat{\lambda}_k$, $\widehat{\sigma}_e^2$, and $\widehat{\phi}$, so the above expression becomes

$$E\big(\mathrm{Var}(V_{i,k}|\mathbf{H},\mathbf{X})|\mathbf{X}\big) \approx \widehat{\mathbf{I}}^{-1}(\widehat{V}_{i,k}) = \left[\frac{1}{\widehat{\sigma}_e^2}\mathbf{B}_k{}'\mathbf{B}_k + \frac{1}{\widehat{\lambda}_k}\widehat{\mathbf{L}}\right]^{-1}_{(i,i)}. \tag{25}$$

So, the total variance for $\widehat{V}_{i,k}$ is approximated by

$$\widehat{\mathrm{Var}}(V_{i,k}|\mathbf{X}) = \mathrm{Var}\big(\widehat{V}_{i,k}(\mathbf{H})\big) + \frac{1}{C}\sum_{c=1}^{C}\left[\widehat{\mathbf{I}}^{-1}\big(\widehat{\mathbf{V}}_k(\mathbf{H}^{[c]})\big)\right]_{(i,i)} = {\xi_{i,k}}^2. \tag{26}$$

Finally, pointwise credible intervals for each estimated cell-type proportion $\widehat{V}_{i,k}(\mathbf{H})$ are constructed by

$$\widehat{V}_{i,k}(\mathbf{H}) \pm 1.96\sqrt{{\xi_{i,k}}^2}. \tag{27}$$

#### Web Appendix G: Inverse transformations with the ZI-HGT + CARD

Let $\mathbf{BV}^{[c]'} = \mathbf{BV}(\mathbf{H}^{[c]})'$. To predict on the original scale of the data, one can produce $C$ posterior predictive replicates (Meng, 1994)

$$X_{i,j}^{[0][c],\mathrm{new}}\big|\mathbf{BV}^{[c]'} \sim \mathrm{Bernoulli}\left(\frac{\exp(\mathbf{BV}^{[c]'})}{1+\exp(\mathbf{BV}^{[c]'})}\right). \tag{28}$$

Let $X_{a,b}^{[c],\text{new}} = X_{a,b}^{[0][c],\text{new}}\ \forall(a,b)$ such that $X_{a,b}^{[0][c],\text{new}} = 0$. Then, $\forall(y,z)$ such that $X_{y,z}^{[0][c],\text{new}} = 1$, we generate

$$X_{y,z}^{[c],\text{new}} \Big| X_{y,z}^{[0][c],\text{new}} = 1, \mathbf{BV}^{[c]'} \sim \text{Poisson}\left(\{\mathbf{BV}^{[c]'}\}_{y,z}\right). \tag{29}$$

Then, averages and variances across $\mathbf{X}^{[c],\text{new}}$ can be used for inference on the original scale of the data. However, posterior predictive data on the original scale may not be desirable due to the zero-inflation. A major part of the original difficulty with interpreting $X_{i,j}$ was that the zero data was already on a different scale than the non-zero data in some sense, as it is unknown whether a zero datum represents a true lack of gene expression or is an artifact of limited sequencing depth. Consequently, we recommend using averages across $c$ of $\mathbf{BV}^{[c']}$ for prediction, not data on the original scale.

#### Web Appendix H: Further discussion of CARD

##### *Review of* $\mathbf{B}$

In CARD's cell-type deconvolution method, $\mathbf{B}$ is the $G \times K$ reference basis matrix. $\mathbf{B}$ is constructed from the reference scRNA-seq dataset, and it contains cell-type specific average expression level of all informative genes. If a gene's average expression in a given cell type is at least a 1.25 log-fold higher than the average expression level of every other included cell type, it is dubbed cell-type informative and included in $\mathbf{B}$. $\mathbf{B}$ is built outside of CARD's inferential procedure on the cell-type proportions $\mathbf{V}$, and as such is considered a fixed input to the CARD model. We describe the construction of $\mathbf{B}$, following Ma and Zhou (2022), below.

For ease in understanding, we begin with the following notation.

- Let $i$ index locations.
- Let $g$ index cell-type informative genes.

- Let $k$ index cell types.
- Let $c$ index individual cells.
- Let $C_{i,k}$ be the set of all cells at location $i$ of type $k$.
- Let $n_{i,k}$ be the number of cells at location $i$ of type $k$.
- Let $x_{i,g}$ be the gene expression (or read counts) at location $i$ for gene $g$.
- Let $x_{i,g,c}$ be the gene expression (or read counts) at location $i$ for gene $g$ from cell $c$.
- Let $N$ be the total number of locations in the ST data.

Then $\mathbf{B}$ is constructed by

$$B_{i,g,k} = \frac{\sum_{c \in C_{i,k}} x_{i,g,c}}{n_{i,k}} = \frac{\sum_{c \in C_{i,k}} x_{i,g,c}}{\sum_{c \in C_{i,k}} \sum_{g'=1}^{G} x_{i,g',c}} \frac{\sum_{c \in C_{i,k}} \sum_{g'=1}^{G} x_{i,g',c}}{n_{i,k}} = \theta_{i,g,k} S_{i,k}, \tag{30}$$

where $\theta_{i,g,k}$ is the relative abundance at location $i$ of gene $g$ from cell type $k$ and $S_{i,k}$ is the mean number of total read counts (i.e., total gene expression) for cells at location $i$ of type $k$. It is then assumed that for any cell type $k$ and any location $i$ the mean relative abundance of gene $g$ $\theta_{i,g,k}$ is the same as the mean in the scRNA-seq data, which is denoted $\theta_{g,k}$. It is also assumed that the average number of total read counts for any cell type $k$ and any location $i$ has the same mean as the scRNA-seq data.

Under the two assumptions, the scRNA-seq reference data is used to estimate $\mathbf{B} = \{B_{g,k}\}$, where $B_{g,k} = \theta^{sc}_{g,k} S^{sc}_{k}$. $\theta^{sc}_{g,k}$ is the mean relative abundance of gene $g$ for cell type $k$, and it is found from the scRNA-seq data. $S^{sc}_{k}$ is the average total gene expression for cells of cell type $k$, and it is also determined from the scRNA-seq data. Essentially, the two assumptions previously mentioned can be summarized as $\frac{\sum_{i=1}^{N} \theta_{i,g,k}}{N} \approx \theta^{sc}_{g,k}$ and $\frac{\sum_{i=1}^{N} S_{i,k}}{N} \approx S^{sc}_{k}$, where the left side of each equation represents the ST data and the right side the scRNA-seq data. If these hold, which is a reasonable assumption so long as the ST and scRNA-seq data come from similar samples and technologies, then $\frac{\sum_{i=1}^{N} B_{i,g,k}}{N} \approx B_{g,k}$. Therefore, the gene expression at location $i$ for gene $g$,

$$x_{i,g} = \sum_k B_{g,k} V_{i,k} + \epsilon_{g,i}, \tag{31}$$

where $V_{i,k}$ is the proportion of cells at location $i$ belonging to cell type $k$ and $\epsilon_{g,i}$ is a normally distributed error term. Put into matrix form, this becomes CARD's non-negative matrix factorization model

$$\mathbf{X} = \mathbf{B}\mathbf{V}' + \mathbf{E}. \tag{32}$$

*Review of CARD's Normal Assumption*

As discussed in the preceding section, CARD (Ma and Zhou, 2022) models the gene expression at location $i$ for gene $g$ by

$$x_{i,g} = \sum_k B_{g,k} V_{i,k} + \epsilon_{g,i}, \tag{33}$$

where $V_{i,k}$ is the proportion of cells at location $i$ belonging to cell type $k$ and $\epsilon_{g,i}$ is a Gaussian error term. Therefore, $x_{i,g} \sim N\left(\sum_k B_{g,k} V_{i,k}, \sigma_e^2\right)$. However, ST data are often highly zero-inflated and naturally count-valued and therefore violate this assumption. The model-data mismatch leads to major concerns. CARD interprets the zeros in the data as informative observations drawn from the lower tail of a normal distribution. In reality, many of the zeros are better understood as missing signal due to technical dropout.

In CARD's empirical hierarchical Bayesian model (EHBM), the negative log-likelihood includes the quadratic term $(\mathbf{X} - \mathbf{B}\mathbf{V}')'(\mathbf{X} - \mathbf{B}\mathbf{V}')$. When the data $\mathbf{X}$ is highly zero-inflated, minimizing this term causes the model to shrink $\mathbf{V}$ toward values that reproduce the zeros, even when those zeros are due to missing data. Moreover, the spatial smoothing prior on $\mathbf{V}$ propagates this bias to neighboring locations, further distorting the estimated spatial distribution of cell types.

The ZI-HGT framework addresses this issue by transforming $\mathbf{X}$ into $\mathbf{H}$, a transformation that better fits with the assumptions of CARD. The ZI-HGT enables CARD to perform more robust and accurate spatial deconvolution.

**Web Tables**

| Sample | Sparsity | Sample | Sparsity |
|---|---|---|---|
| 1 | 91% | 7 | 90% |
| 2 | 90% | 8 | 88% |
| 3 | 87% | 9 | 90% |
| 4 | 86% | 10 | 87% |
| 5 | 88% | 11 | 89% |
| 6 | 91% | 12 | 87% |

Table 1: Sparsity levels for all OSCC ST samples (Arora et al., 2023).

| Number of Transcript Reads | Frequency | Number of Transcript Reads | Frequency |
|---|---|---|---|
| 0 | 16384736 | 11 | 4481 |
| 1 | 1085656 | 12 | 3635 |
| 2 | 233420 | 13 | 2841 |
| 3 | 78919 | 14 | 2395 |
| 4 | 36662 | 15 | 1852 |
| 5 | 23043 | 16 | 1508 |
| 6 | 16769 | 17 | 1281 |
| 7 | 12144 | 18 | 1050 |
| 8 | 9228 | 19 | 888 |
| 9 | 7016 | 20 | 692 |
| 10 | 5743 | $> 20$ | 5605 |

Table 2: Frequencies of number of transcript reads across 15,844 transcripts and 1,131 spatial locations in the OSCC ST data Sample 1 (Arora et al., 2023).

| Number of Transcript Reads | Frequency | Number of Transcript Reads | Frequency |
|---|---|---|---|
| 0 | 24548148 | 11 | 9502 |
| 1 | 1926440 | 12 | 7904 |
| 2 | 422008 | 13 | 6710 |
| 3 | 151485 | 14 | 5461 |
| 4 | 69296 | 15 | 4658 |
| 5 | 43445 | 16 | 3926 |
| 6 | 30105 | 17 | 3352 |
| 7 | 22575 | 18 | 2884 |
| 8 | 17438 | 19 | 2418 |
| 9 | 14212 | 20 | 2120 |
| 10 | 11471 | $> 20$ | 20818 |

Table 3: Frequencies of number of transcript reads across 15,624 transcripts and 1,749 spatial locations in the OSCC ST data Sample 2.

**Web Figures**

| **Number of Transcript Reads** | **Frequency** | **Number of Transcript Reads** | **Frequency** |
|---|---|---|---|
| 0 | 13356156 | 11 | 6488 |
| 1 | 1300680 | 12 | 5545 |
| 2 | 319755 | 13 | 4628 |
| 3 | 103602 | 14 | 3981 |
| 4 | 48044 | 15 | 3252 |
| 5 | 28181 | 16 | 2950 |
| 6 | 19297 | 17 | 2474 |
| 7 | 14499 | 18 | 2142 |
| 8 | 11702 | 19 | 1886 |
| 9 | 9113 | 20 | 1678 |
| 10 | 7940 | $> 20$ | 15926 |

Table 4: Frequencies of number of transcript reads across 16,023 transcripts and 953 spatial locations in the OSCC ST data Sample 3.

| **Number of Transcript Reads** | **Frequency** | **Number of Transcript Reads** | **Frequency** |
|---|---|---|---|
| 0 | 30165798 | 11 | 16943 |
| 1 | 3211102 | 12 | 14099 |
| 2 | 933258 | 13 | 11804 |
| 3 | 313887 | 14 | 9923 |
| 4 | 149420 | 15 | 8607 |
| 5 | 87524 | 16 | 7262 |
| 6 | 58206 | 17 | 6319 |
| 7 | 42259 | 18 | 5526 |
| 8 | 32739 | 19 | 4971 |
| 9 | 25965 | 20 | 4398 |
| 10 | 21159 | $> 20$ | 54255 |

Table 5: Frequencies of number of transcript reads across 18,364 transcripts and 1,916 spatial locations in the OSCC ST data Sample 4.

| Number of Transcript Reads | Frequency | Number of Transcript Reads | Frequency |
|---|---|---|---|
| 0 | 24273032 | 11 | 11844 |
| 1 | 2343248 | 12 | 9916 |
| 2 | 581876 | 13 | 8018 |
| 3 | 167094 | 14 | 6932 |
| 4 | 77153 | 15 | 5767 |
| 5 | 47082 | 16 | 5101 |
| 6 | 33969 | 17 | 4248 |
| 7 | 26728 | 18 | 3622 |
| 8 | 21714 | 19 | 3026 |
| 9 | 17677 | 20 | 2782 |
| 10 | 14523 | $> 20$ | 31598 |

Table 6: Frequencies of number of transcript reads across 16,585 transcripts and 1,670 spatial locations in the OSCC ST data Sample 5.

| **Number of Transcript Reads** | **Frequency** | **Number of Transcript Reads** | **Frequency** |
|---|---|---|---|
| 0 | 48538068 | 11 | 13713 |
| 1 | 3205204 | 12 | 10725 |
| 2 | 722530 | 13 | 8297 |
| 3 | 252663 | 14 | 6665 |
| 4 | 118643 | 15 | 5515 |
| 5 | 75209 | 16 | 4426 |
| 6 | 54977 | 17 | 3790 |
| 7 | 41212 | 18 | 3297 |
| 8 | 31049 | 19 | 2912 |
| 9 | 23114 | 20 | 2619 |
| 10 | 17748 | $> 20$ | 34721 |

Table 7: Frequencies of number of transcript reads across 16,791 transcripts and 3,167 spatial locations in the OSCC ST data Sample 6.

| **Number of Transcript Reads** | **Frequency** | **Number of Transcript Reads** | **Frequency** |
|---|---|---|---|
| 0 | 38223364 | 11 | 6577 |
| 1 | 2983694 | 12 | 5380 |
| 2 | 694834 | 13 | 4382 |
| 3 | 244663 | 14 | 3790 |
| 4 | 109904 | 15 | 3131 |
| 5 | 58955 | 16 | 2792 |
| 6 | 36609 | 17 | 2500 |
| 7 | 24037 | 18 | 2165 |
| 8 | 16211 | 19 | 1938 |
| 9 | 11685 | 20 | 1790 |
| 10 | 8697 | $> 20$ | 14036 |

Table 8: Frequencies of number of transcript reads across 17,324 transcripts and 2,451 spatial locations in the OSCC ST data Sample 7.

| Number of Transcript Reads | Frequency | Number of Transcript Reads | Frequency |
|---|---|---|---|
| 0 | 35303254 | 11 | 13553 |
| 1 | 3463278 | 12 | 10717 |
| 2 | 826085 | 13 | 8859 |
| 3 | 277967 | 14 | 7050 |
| 4 | 136501 | 15 | 5733 |
| 5 | 78597 | 16 | 4800 |
| 6 | 52746 | 17 | 3992 |
| 7 | 38062 | 18 | 3469 |
| 8 | 28494 | 19 | 3015 |
| 9 | 21695 | 20 | 2745 |
| 10 | 16911 | $> 20$ | 34647 |

Table 9: Frequencies of number of transcript reads across 17,058 transcripts and 2,365 spatial locations in the OSCC ST data Sample 8.

| Number of Transcript Reads | Frequency | Number of Transcript Reads | Frequency |
|---|---|---|---|
| 0 | 51622846 | 11 | 14265 |
| 1 | 3943341 | 12 | 11611 |
| 2 | 881299 | 13 | 9475 |
| 3 | 317880 | 14 | 7929 |
| 4 | 148975 | 15 | 6735 |
| 5 | 86174 | 16 | 5790 |
| 6 | 59166 | 17 | 5134 |
| 7 | 43023 | 18 | 4509 |
| 8 | 31641 | 19 | 4017 |
| 9 | 24036 | 20 | 3465 |
| 10 | 18481 | $> 20$ | 46188 |

Table 10: Frequencies of number of transcript reads across 17,206 transcripts and 3,330 spatial locations in the OSCC ST data Sample 9.

| Number of Transcript Reads | Frequency | Number of Transcript Reads | Frequency |
|---|---|---|---|
| 0 | 40339285 | 11 | 21537 |
| 1 | 3833913 | 12 | 18340 |
| 2 | 992599 | 13 | 16010 |
| 3 | 338928 | 14 | 14190 |
| 4 | 161908 | 15 | 12836 |
| 5 | 94874 | 16 | 11164 |
| 6 | 64162 | 17 | 10204 |
| 7 | 47749 | 18 | 9112 |
| 8 | 37260 | 19 | 8425 |
| 9 | 30870 | 20 | 7322 |
| 10 | 25334 | $> 20$ | 92393 |

Table 11: Frequencies of number of transcript reads across 17,155 transcripts and 2,693 spatial locations in the OSCC ST data Sample 10.

| Number of Transcript Reads | Frequency | Number of Transcript Reads | Frequency |
|---|---|---|---|
| 0 | 29005205 | 11 | 14916 |
| 1 | 2467025 | 12 | 13174 |
| 2 | 574498 | 13 | 11265 |
| 3 | 180888 | 14 | 9601 |
| 4 | 86038 | 15 | 8234 |
| 5 | 50754 | 16 | 7078 |
| 6 | 36686 | 17 | 6165 |
| 7 | 28928 | 18 | 5265 |
| 8 | 23414 | 19 | 4423 |
| 9 | 19736 | 20 | 3930 |
| 10 | 16849 | $> 20$ | 42292 |

Table 12: Frequencies of number of transcript reads across 16,658 transcripts and 1,958 spatial locations in the OSCC ST data Sample 11.

| Number of Transcript Reads | Frequency | Number of Transcript Reads | Frequency |
|---|---|---|---|
| 0 | 21435134 | 11 | 10886 |
| 1 | 2162934 | 12 | 8888 |
| 2 | 565847 | 13 | 7397 |
| 3 | 186720 | 14 | 6273 |
| 4 | 82304 | 15 | 5241 |
| 5 | 48714 | 16 | 4527 |
| 6 | 33463 | 17 | 3703 |
| 7 | 25028 | 18 | 3181 |
| 8 | 19852 | 19 | 2643 |
| 9 | 16029 | 20 | 2268 |
| 10 | 12846 | $> 20$ | 23468 |

Table 13: Frequencies of number of transcript reads across 16,522 transcripts and 1,493 spatial locations in the OSCC ST data Sample 12.

| Pathologist Tissue Layer | Dominant Cell Type |
|---|---|
| Artery/Vein | Endothelial Cells |
| Glandular Stroma | Intermediate Fibroblasts |
| Lymphocyte Negative Stroma | ecm-myCAFs |
| Lymphocyte Positive Stroma | Cytotoxic CD8+ T Cells |
| Muscle | Myofibroblasts |
| Lymphocyte Positive Muscles | Myofibroblasts |
| Non-Cancerous Mucosa | Macrophages |
| Squamous Cell Carcinoma | Cancer Cells |
| Artifact/Cautery/Edge Effect/Fold | NA |

Table 14: Dominant cell types for OSCC TME pathologist annotated tissue layers.

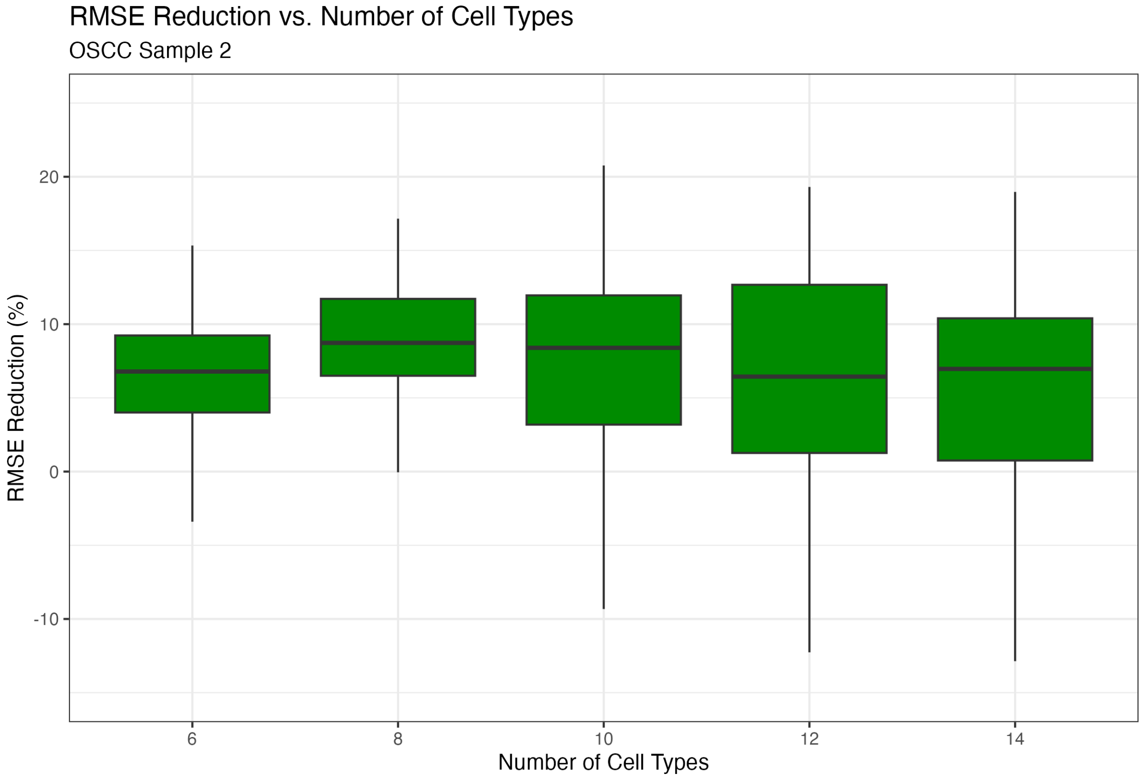


**Figure 1**: RMSE reduction vs. number of cell types. RMSE reduction for the ZI-HGT + CARD across 100 simulated ST datasets for OSCC Sample 2 with 14, 12, 10, 8, and 6 cell types. The SPARSim scRNA-seq hyperparameters were set to 0.05 and 50 for the library factor and $\Phi$ factor respectively, which lead to the most "realistic" simulated ST data. The median RMSE reduction is fairly consistent across the distinct numbers of cell types, but the variance of the RMSE reduction tends to decrease as the number of cell types does.

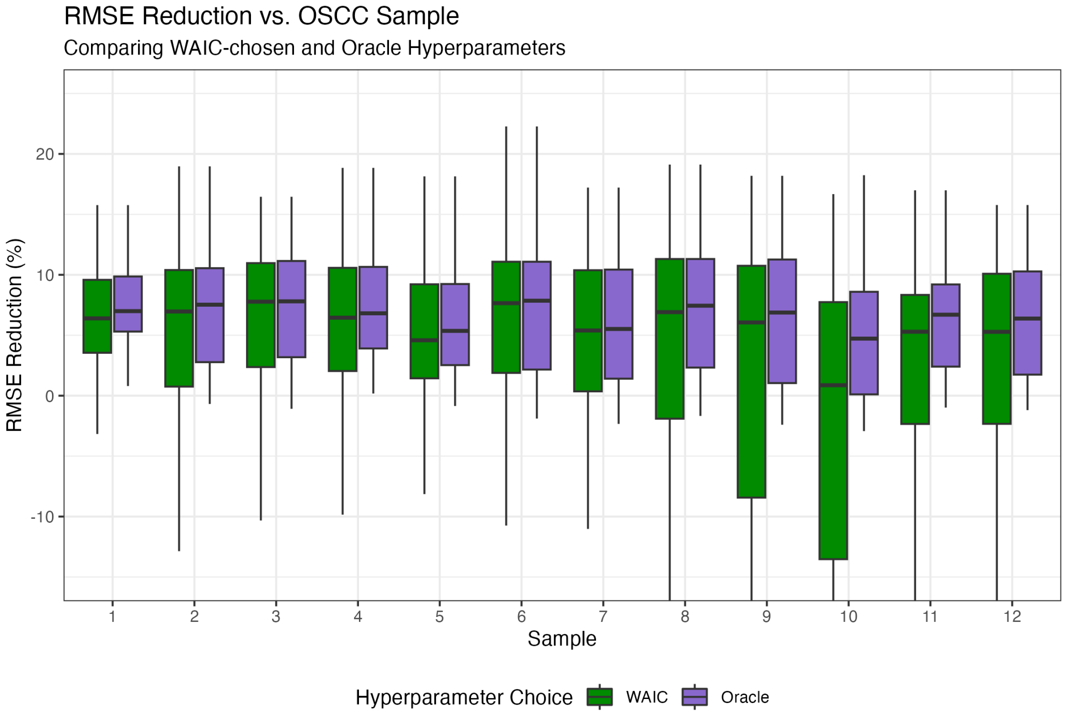


**Figure 2**: RMSE reduction vs. OSCC sample comparing WAIC-chosen vs. oracle hyperparameters. RMSE reduction for the ZI-HGT + CARD across 100 simulated ST datasets over all samples. The SPARSim scRNA-seq hyperparameters were set to 0.05 and 50 for the library factor and $\Phi$ factor respectively. The efficacy of using the WAIC to determine the model hyperparameters was assessed by comparing the RMSE reduction to the RMSE reduction using the oracle hyperparameters (chosen to minimize the RMSE in each simulated dataset). The median RMSE reduction when using the WAIC to choose hyperparameters was within 1% of the median RMSE reduction for the oracle hyperparameters on all samples but Sample 10. The variance in the ZI-HGT + CARD results is notably higher in Samples 9, 10, and to a lesser extent, 8. These samples are uniquely homogenous; at least 75% of the spots were annotated to be of the most prominent tissue layer. No other samples were above 70%, and all but three were below 60%. This leads to simulated ST datasets in which cell-type composition for the overwhelming majority of the spots in the sample is highly homogeneous. Naturally, CARD performs well in scenarios with high spatial autocorrelation and homogeneity (Ma and Zhou, 2022), leading to lesser improvements by applying the ZI-HGT to CARD.

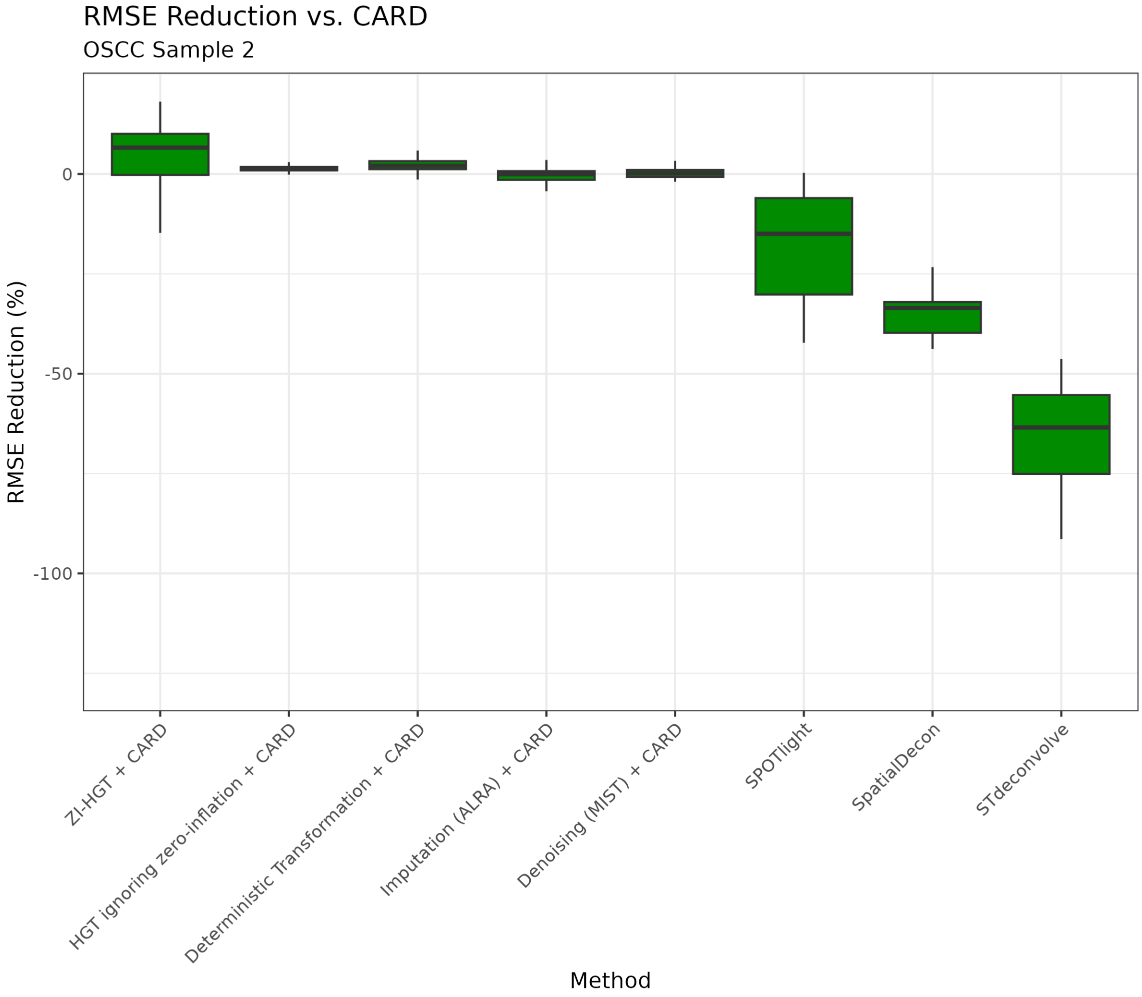


**Figure 3**: RMSE reduction for other methods. RMSE reduction or promotion vs. CARD for the ZI-HGT + CARD, a simple deterministic transformation + CARD (in practice, a $log\,(1+\epsilon+X_{i,j})$ where $\epsilon = 0.05$), ALRA (Linderman et al., 2022), a data imputation technique, + CARD, MIST (Wang et al., 2022), a denoising technique, + CARD, SPOTlight (Elosua-Bayes et al., 2021), SpatialDecon (Danaher et al., 2022), and STdeconvolve (Miller et al., 2022) on 100 simulated ST datasets for OSCC Sample 2. The SPARSim scRNA-seq hyperparameters were set to 0.05 and 50 for the library factor and $\Phi$ factor, respectively, which lead to the most "realistic" simulated ST data. All comparison methods were applied following the authors' instructions.

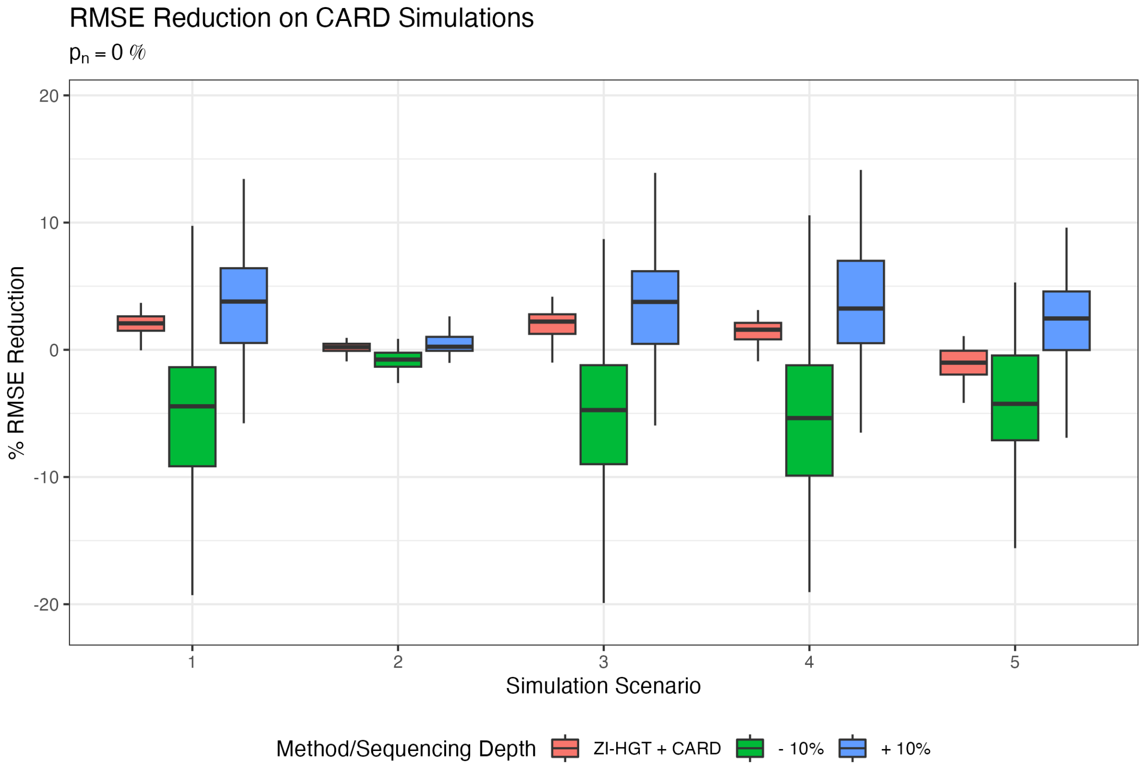


**Figure 4**: RMSE reduction on CARD simulations. RMSE reduction for the ZI-HGT + CARD across 100 simulated MOB ST datasets and various simulation scenarios following the CARD simulation setup with $p_n = 0\%$. The -10% and +10% sequencing depths refer to applying CARD to simulated ST data with accordingly decreased or increased sequencing depth. The ZI-HGT + CARD achieved median RMSE reductions of 2.1%, 0.2%, 2.2%, and 1.6% in scenarios 1-4. For comparison, CARD with +10% (i.e., 110% ) sequencing depth resulted in median RMSE reductions of 3.8%, 0.2%, 3.8%, and 3.2%. Given the low sparsity of the simulated datasets biasing the results against the ZI-HGT, the ZI-HGT + CARD achieving RMSE reduction within 1.7% of CARD with 110% sequencing depth demonstrates our method's viability.

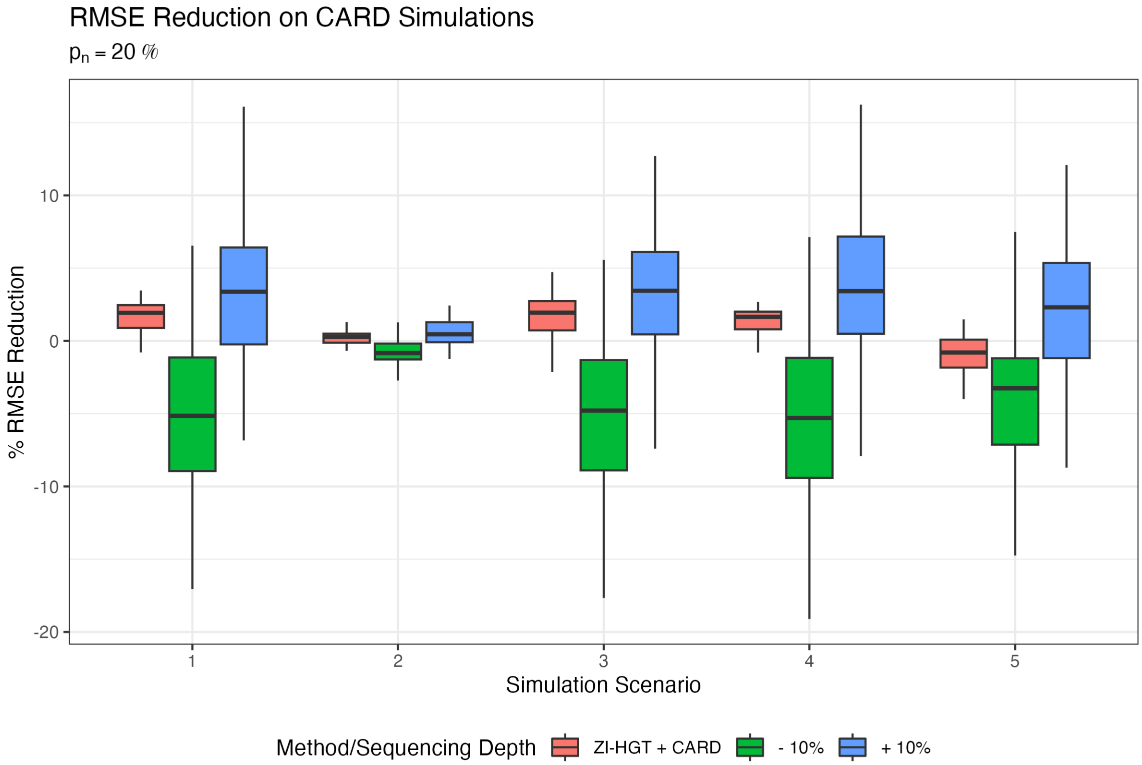


**Figure 5**: RMSE reduction on CARD simulations with $p_n = 20\%$. RMSE reduction for the ZI-HGT + CARD across 100 simulated MOB ST datasets and various simulation scenarios following the CARD simulation setup with 20% "noisy" locations. The -10% and +10% sequencing depths refer to applying CARD to simulated ST data with accordingly decreased or increased sequencing depth.

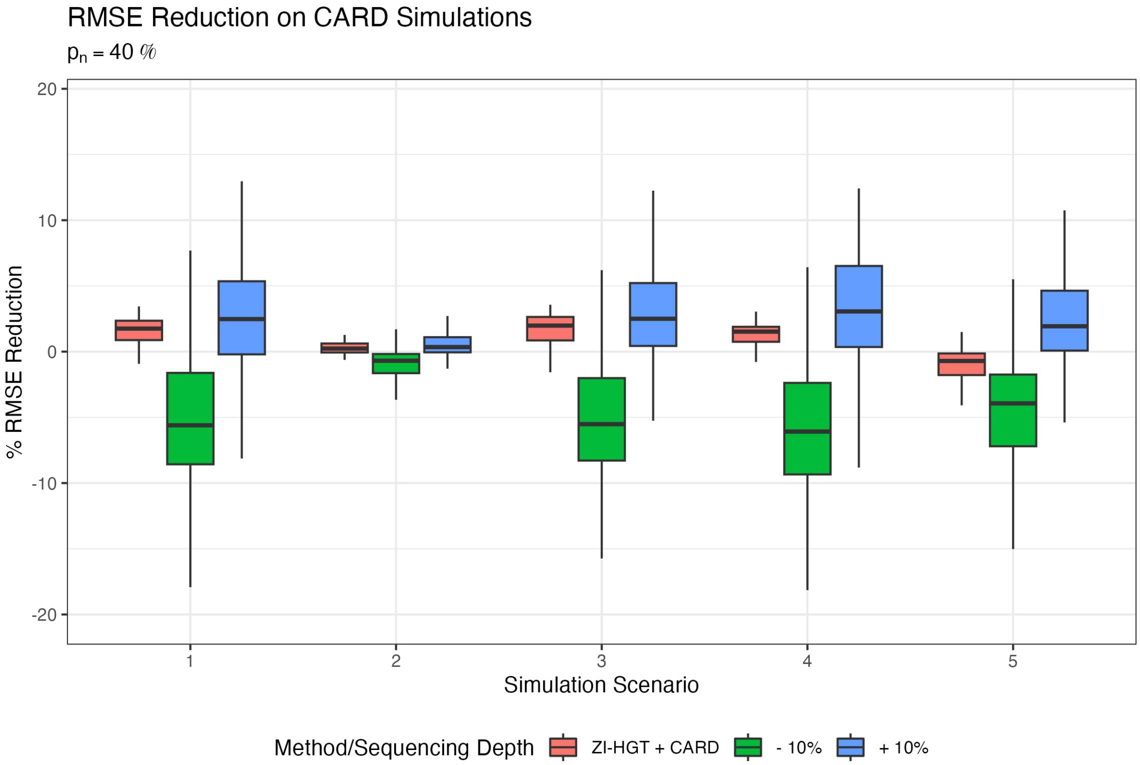


**Figure 6**: RMSE reduction on CARD simulations with $p_n = 40\%$. RMSE reduction for the ZI-HGT + CARD across 100 simulated MOB ST datasets and various simulation scenarios following the CARD simulation setup with 40% "noisy" locations. The -10% and +10% sequencing depths refer to applying CARD to simulated ST data with accordingly decreased or increased sequencing depth.

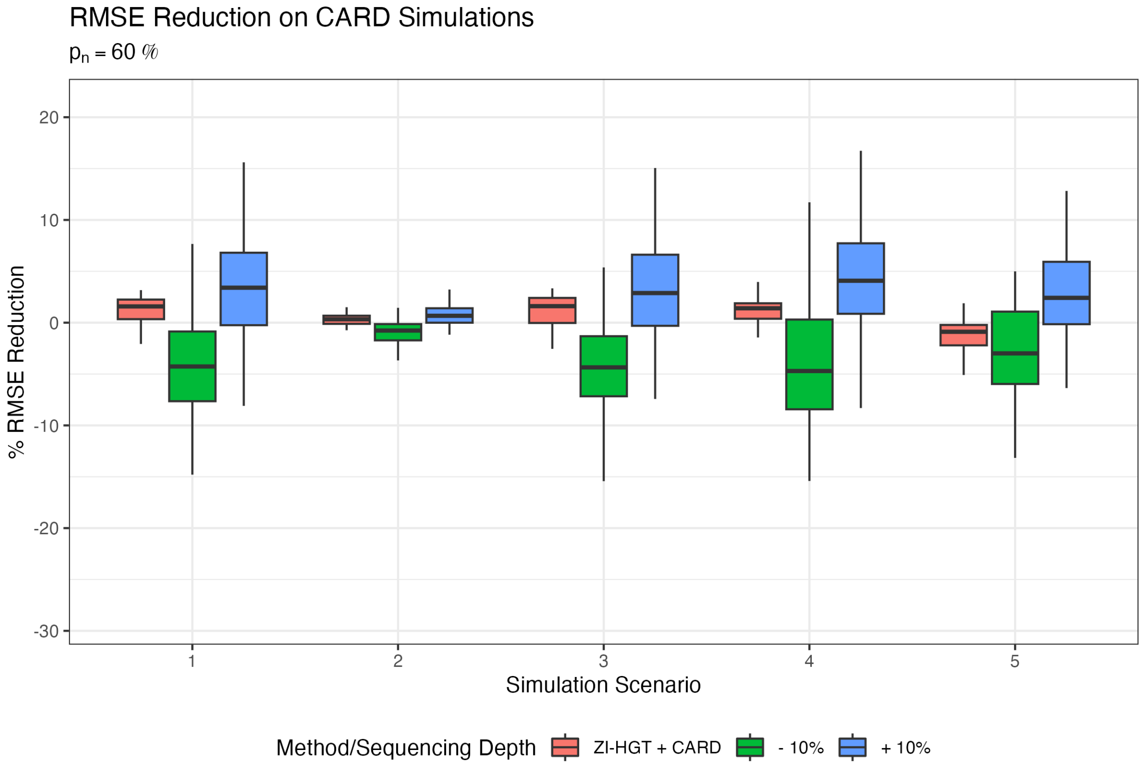


**Figure 7**: RMSE reduction on CARD simulations with $p_n = 60\%$. RMSE reduction for the ZI-HGT + CARD across 100 simulated MOB ST datasets and various simulation scenarios following the CARD simulation setup with 60% "noisy" locations. The -10% and +10% sequencing depths refer to applying CARD to simulated ST data with accordingly decreased or increased sequencing depth.

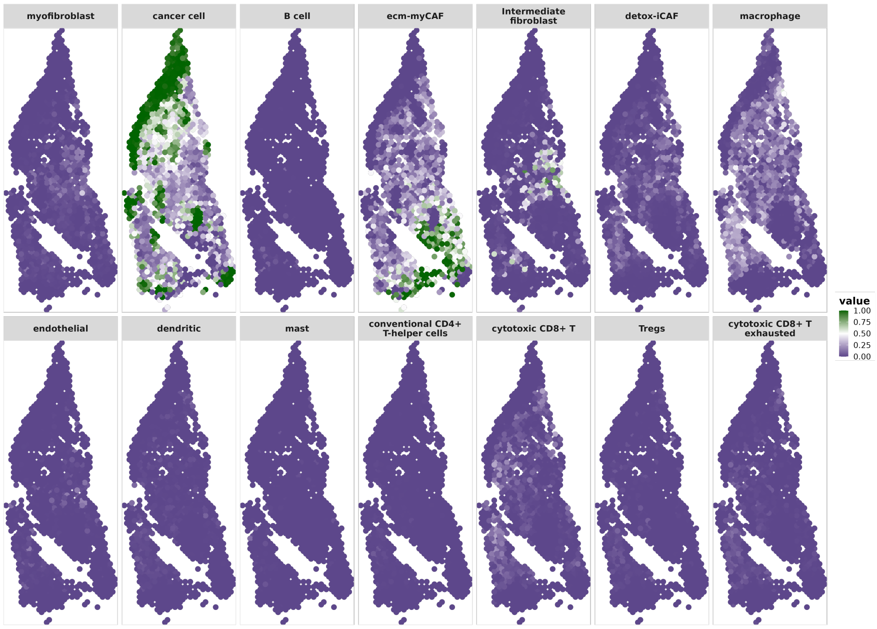


(a) Cell-Type Proportions Estimates

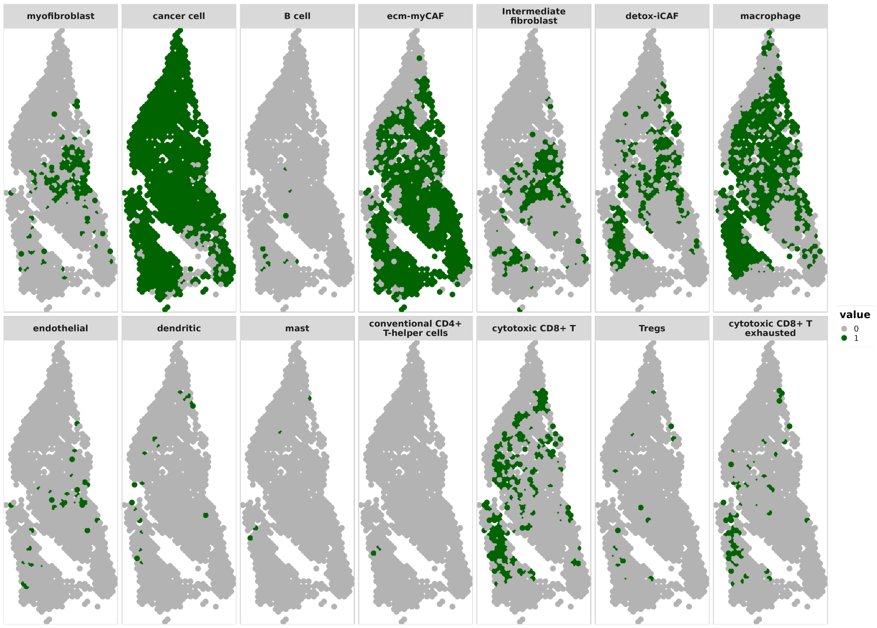


(b) Cell-Type Proportions $> 0.05$

**Figure 8**: OSCC Sample 1 CARD Cell-Type Proportions Estimates. a) Plots of cell-type proportions for all 14 cell types. b) Binary plots of cell-type proportions $> 0.05$ for all 14 cell types.

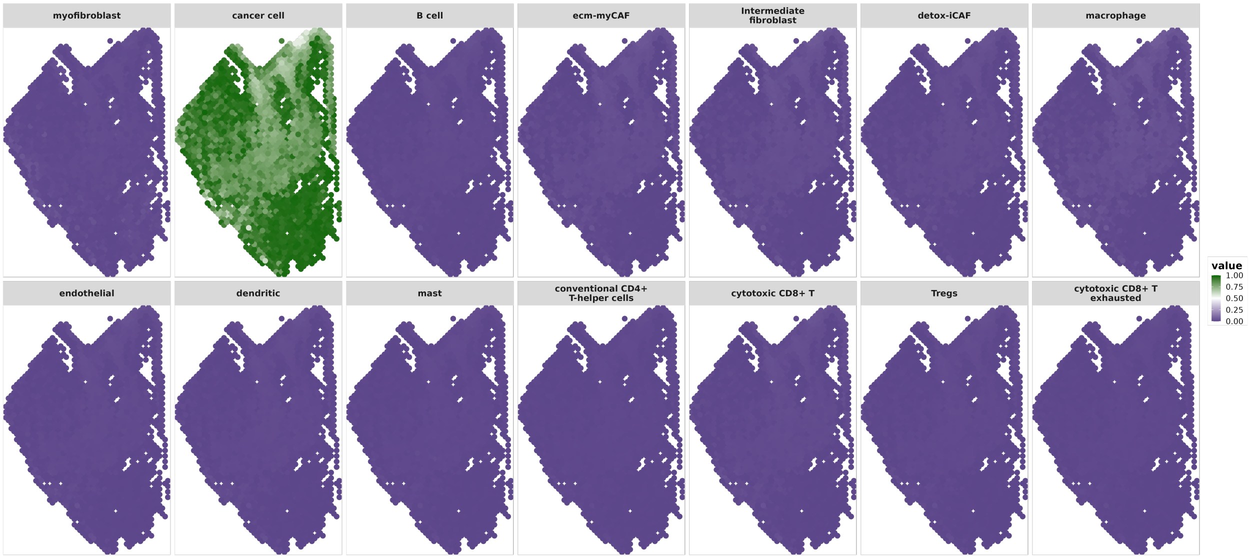


(a) Cell-Type Proportions Estimates

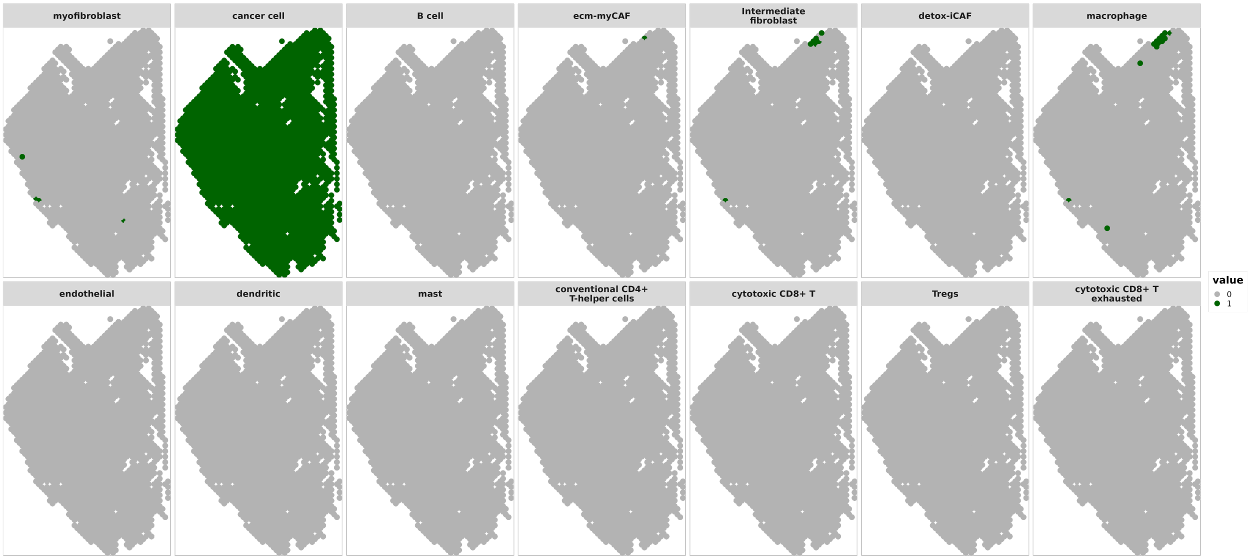


(b) Cell-Type Proportions $> 0.05$

**Figure 9**: OSCC Sample 2 ZI-HGT + CARD Cell-Type Proportions Estimates. a) Plots of cell-type proportions for all 14 cell types. b) Binary plots of cell-type proportions $> 0.05$ for all 14 cell types.

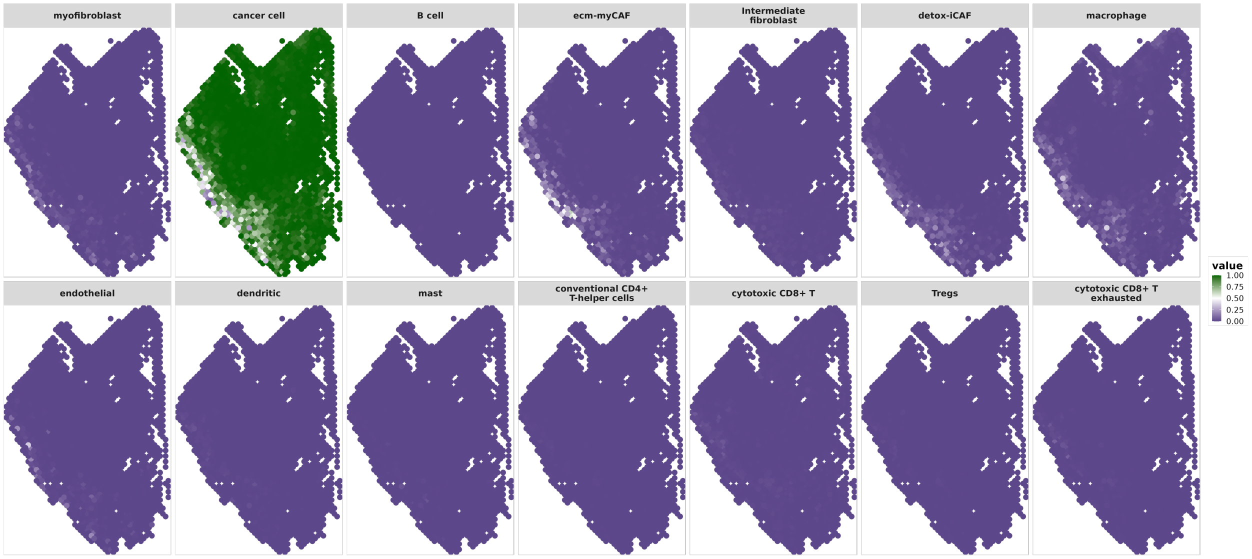


(a) Cell-Type Proportions Estimates

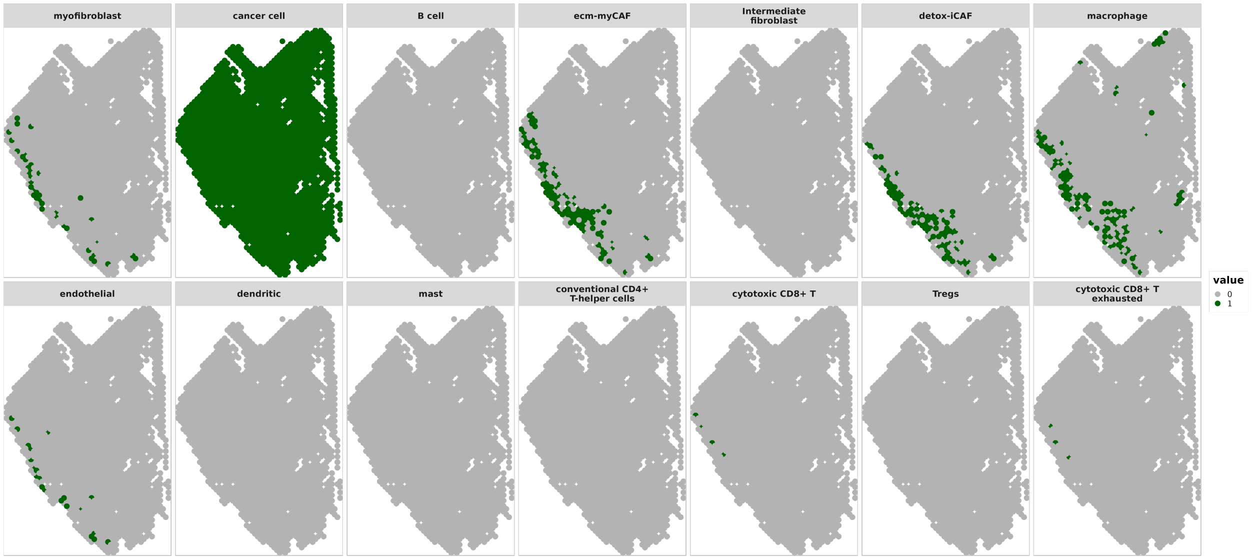


(b) Cell-Type Proportions $> 0.05$

**Figure 10**: OSCC Sample 2 CARD Cell-Type Proportions Estimates. a) Plots of cell-type proportions for all 14 cell types. b) Binary plots of cell-type proportions $> 0.05$ for all 14 cell types.

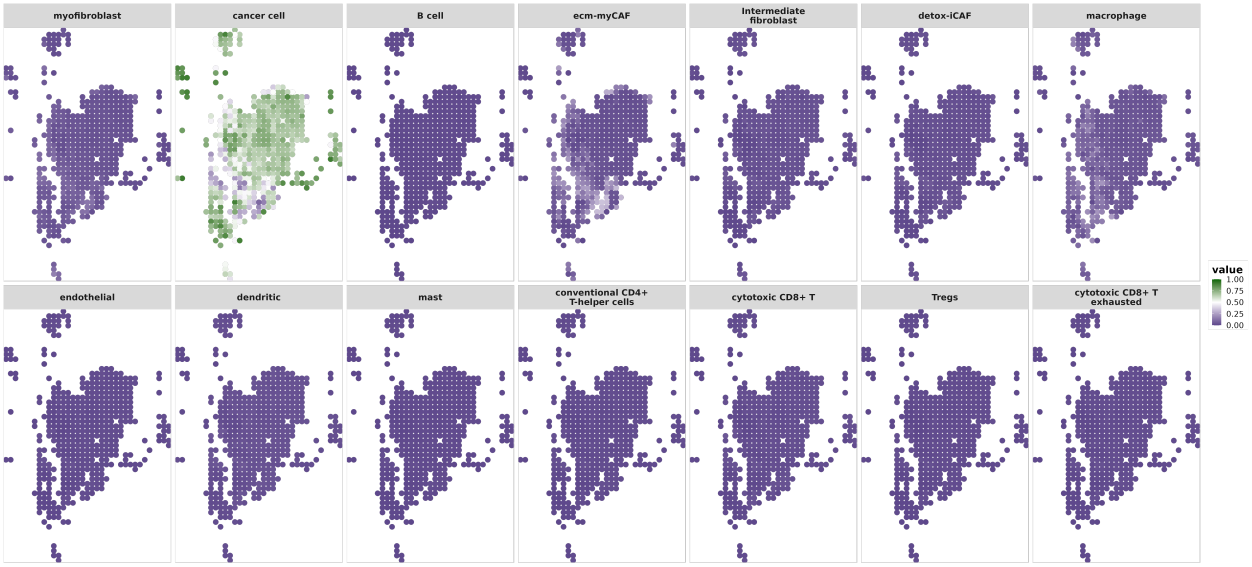


(a) Cell-Type Proportions Estimates

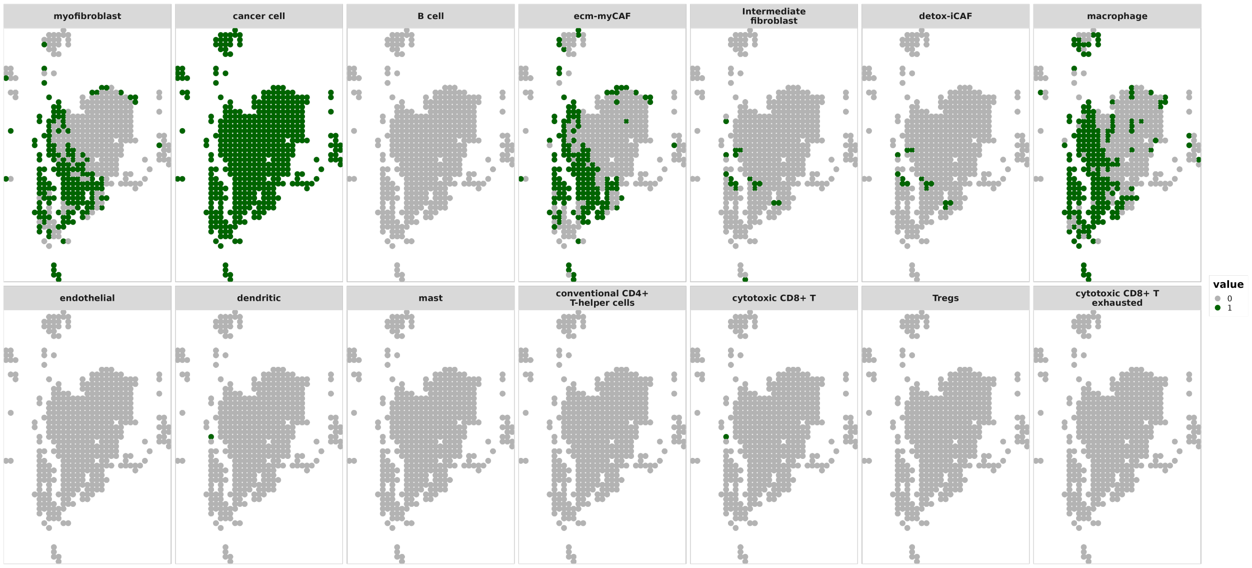


(b) Cell-Type Proportions > 0.05

**Figure 11**: OSCC Sample 3 ZI-HGT + CARD Cell-Type Proportions Estimates. a) Plots of cell-type proportions for all 14 cell types. b) Binary plots of cell-type proportions > 0.05 for all 14 cell types.

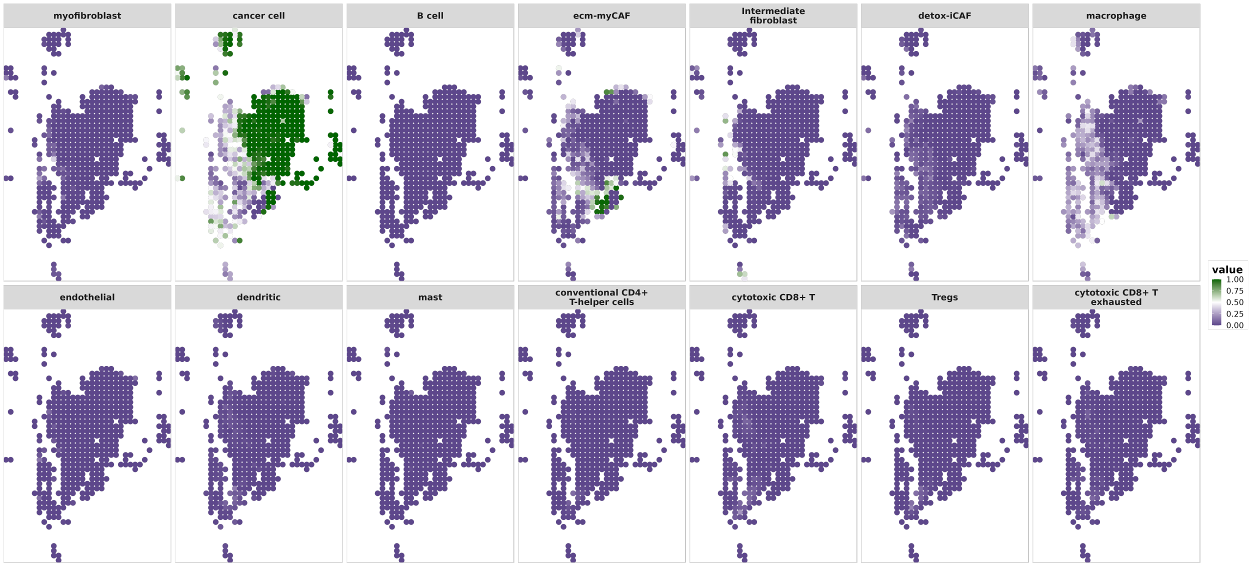


(a) Cell-Type Proportions Estimates

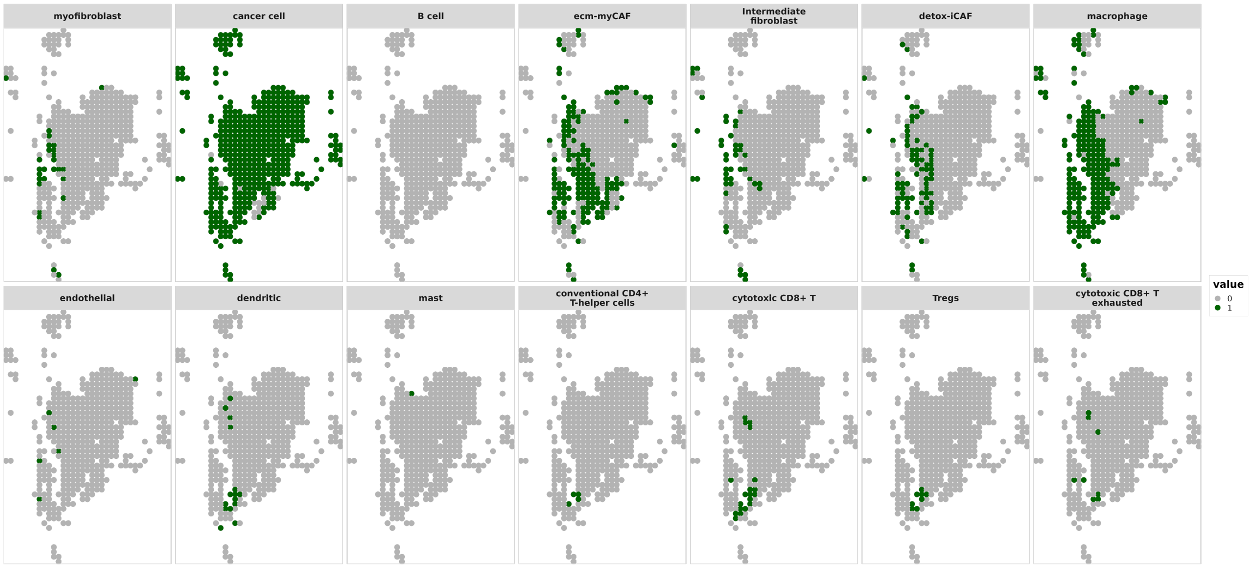


(b) Cell-Type Proportions $> 0.05$

**Figure 12**: OSCC Sample 3 CARD Cell-Type Proportions Estimates. a) Plots of cell-type proportions for all 14 cell types. b) Binary plots of cell-type proportions $> 0.05$ for all 14 cell types.

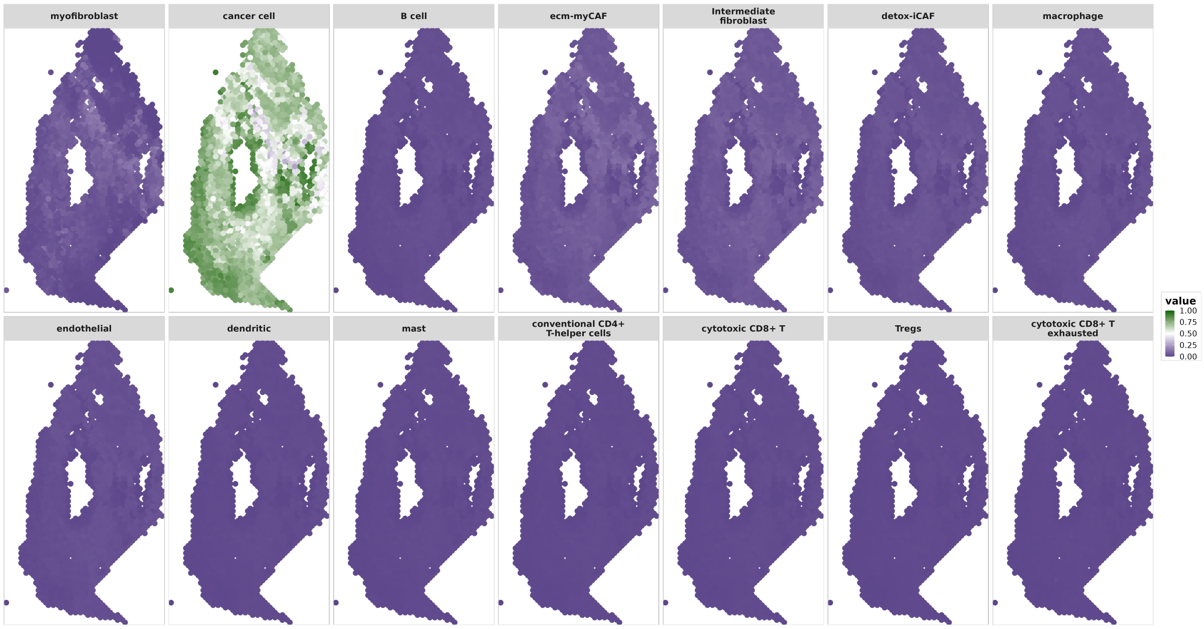


(a) Cell-Type Proportions Estimates

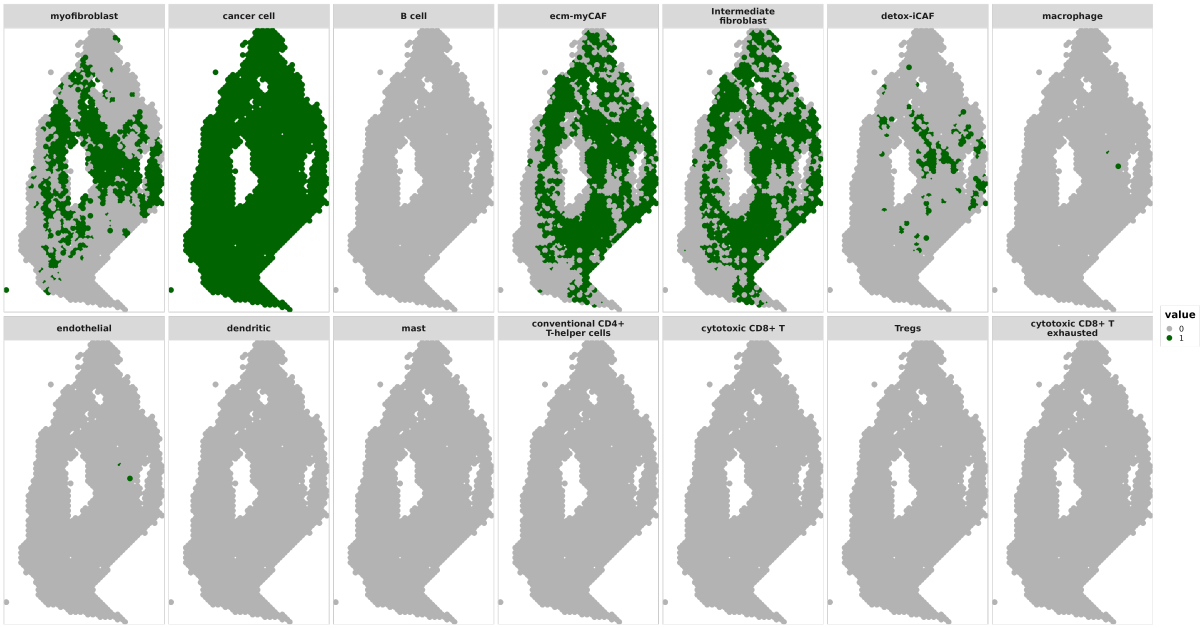


(b) Cell-Type Proportions $> 0.05$

**Figure 13**: OSCC Sample 4 ZI-HGT + CARD Cell-Type Proportions Estimates. a) Plots of cell-type proportions for all 14 cell types. b) Binary plots of cell-type proportions $> 0.05$ for all 14 cell types.

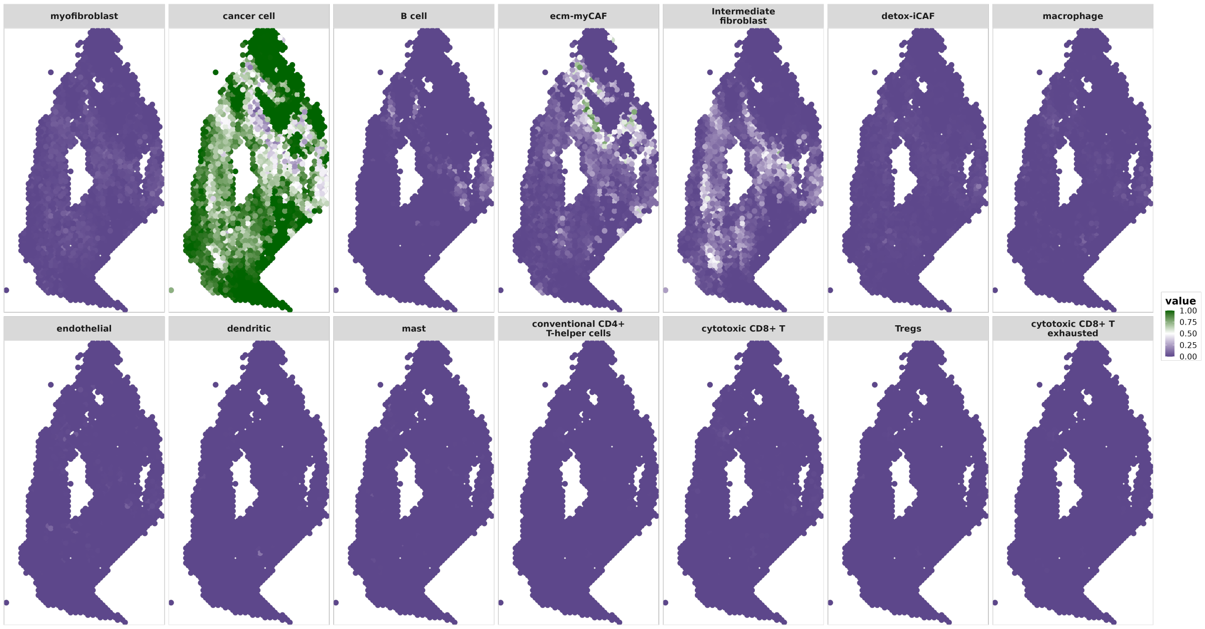


(a) Cell-Type Proportions Estimates

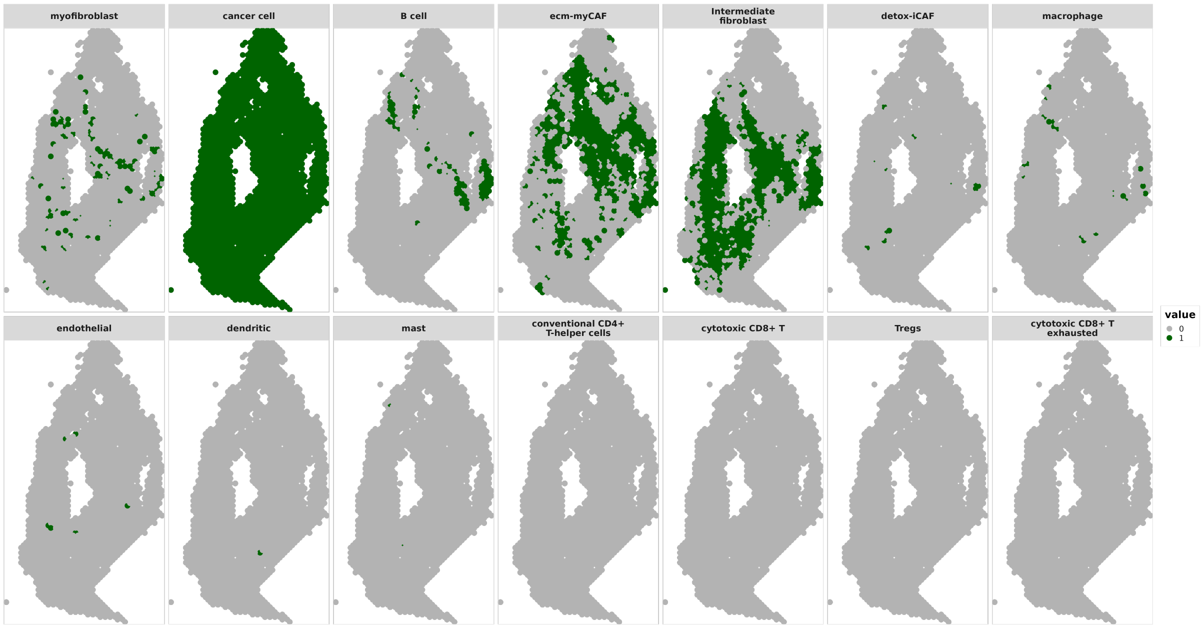


(b) Cell-Type Proportions > 0.05

**Figure 14**: OSCC Sample 4 CARD Cell-Type Proportions Estimates. a) Plots of cell-type proportions for all 14 cell types. b) Binary plots of cell-type proportions > 0.05 for all 14 cell types.

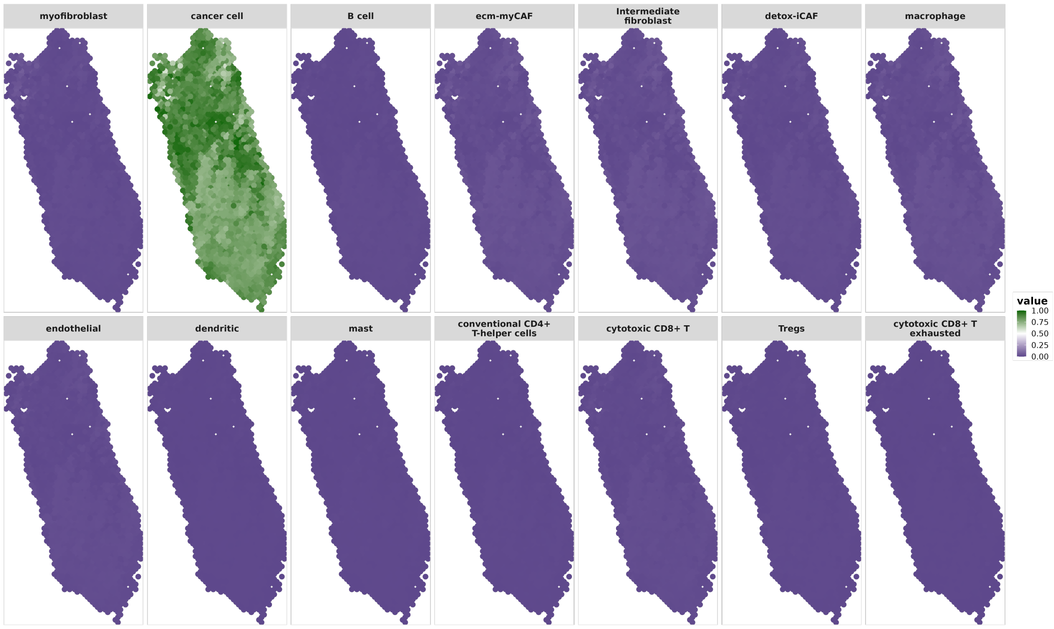


(a) Cell-Type Proportions Estimates

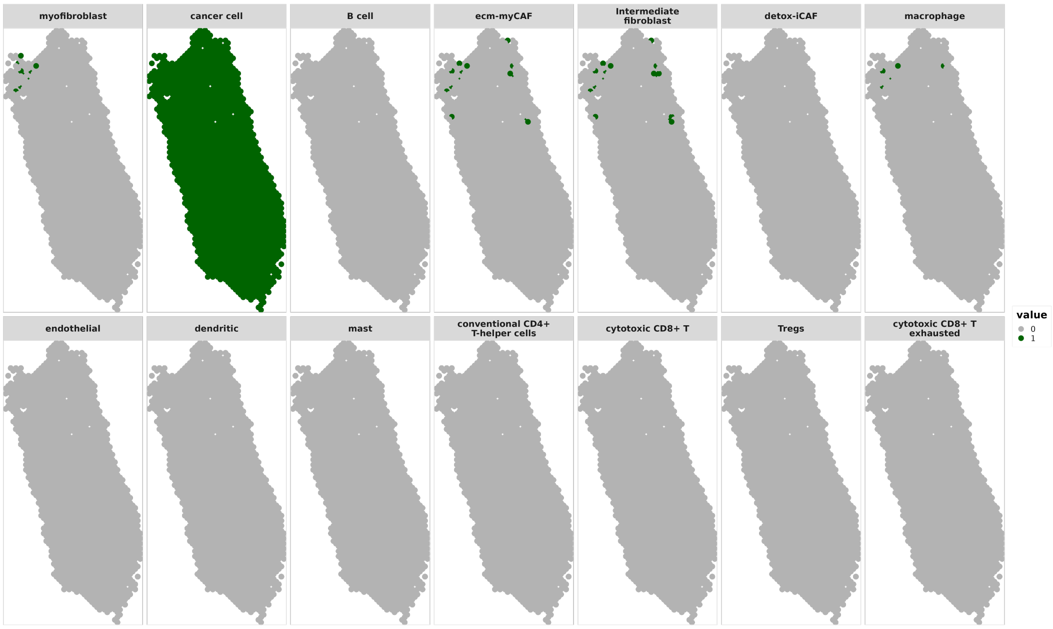


(b) Cell-Type Proportions > 0.05

**Figure 15**: OSCC Sample 5 ZI-HGT + CARD Cell-Type Proportions Estimates. a) Plots of cell-type proportions for all 14 cell types. b) Binary plots of cell-type proportions > 0.05 for all 14 cell types.

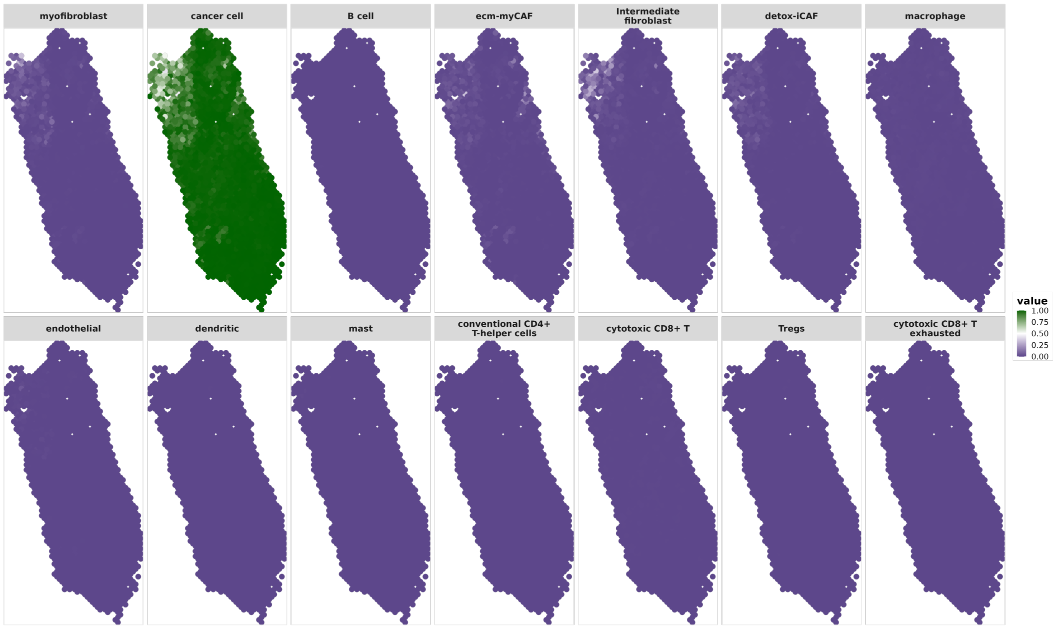


(a) Cell-Type Proportions Estimates

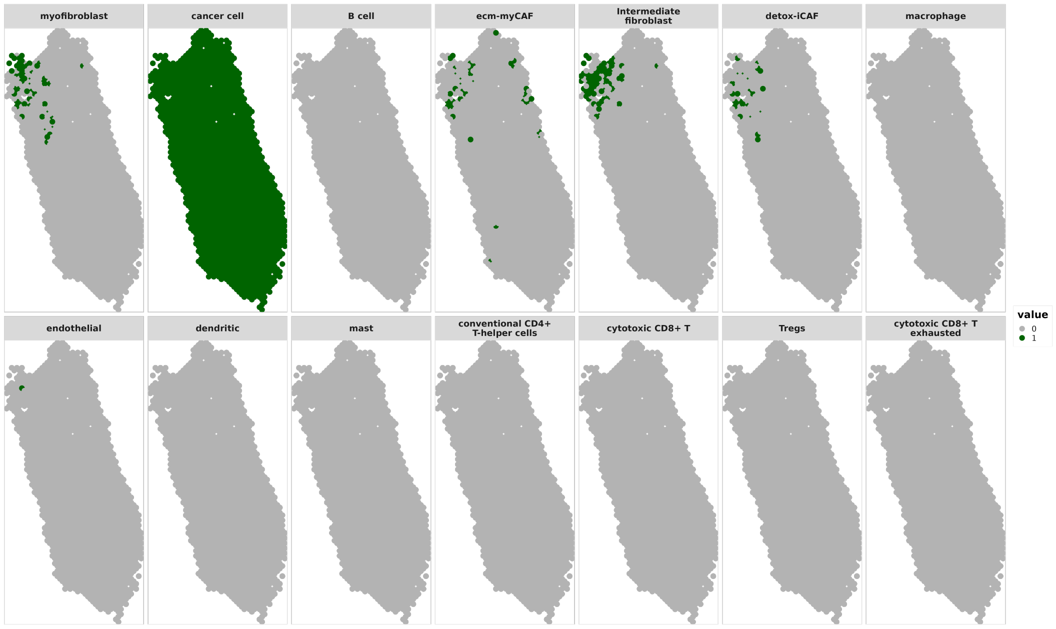


(b) Cell-Type Proportions > 0.05

**Figure 16**: OSCC Sample 5 CARD Cell-Type Proportions Estimates. a) Plots of cell-type proportions for all 14 cell types. b) Binary plots of cell-type proportions > 0.05 for all 14 cell types.

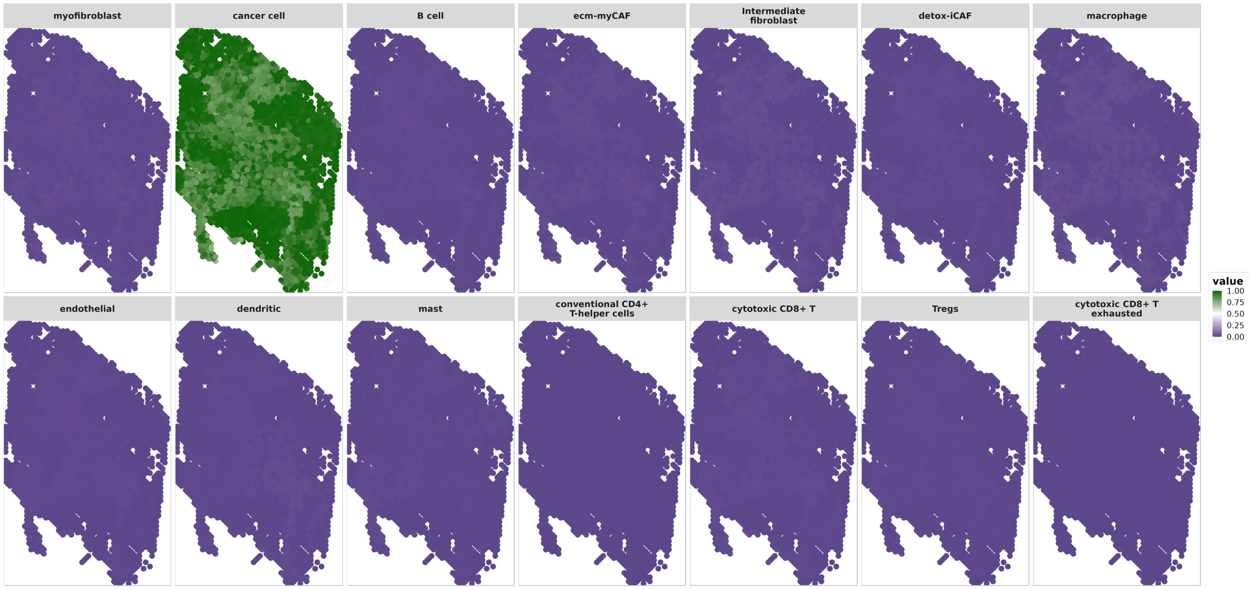


(a) Cell-Type Proportions Estimates

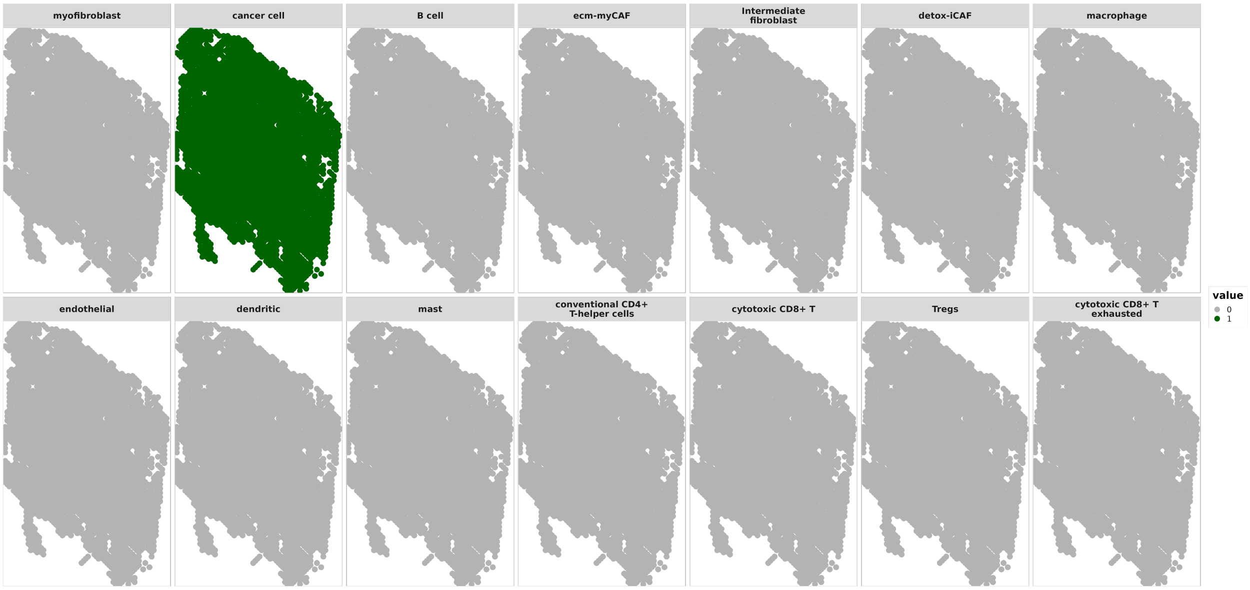


(b) Cell-Type Proportions $> 0.05$

**Figure 17**: OSCC Sample 6 ZI-HGT + CARD Cell-Type Proportions Estimates. a) Plots of cell-type proportions for all 14 cell types. b) Binary plots of cell-type proportions $> 0.05$ for all 14 cell types.

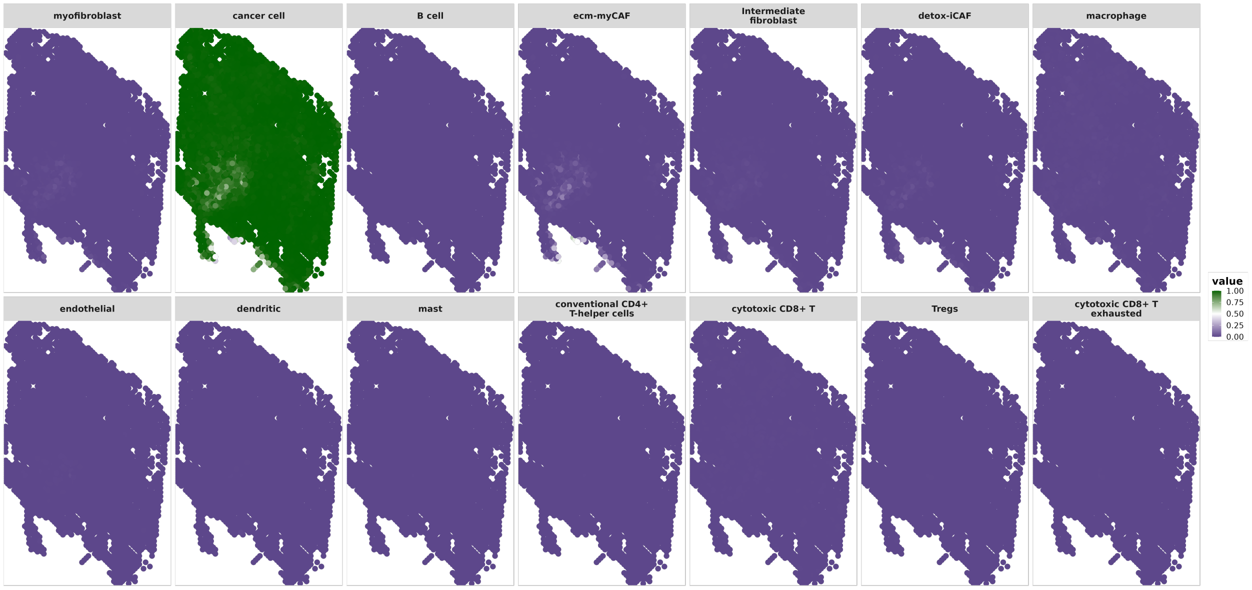


(a) Cell-Type Proportions Estimates

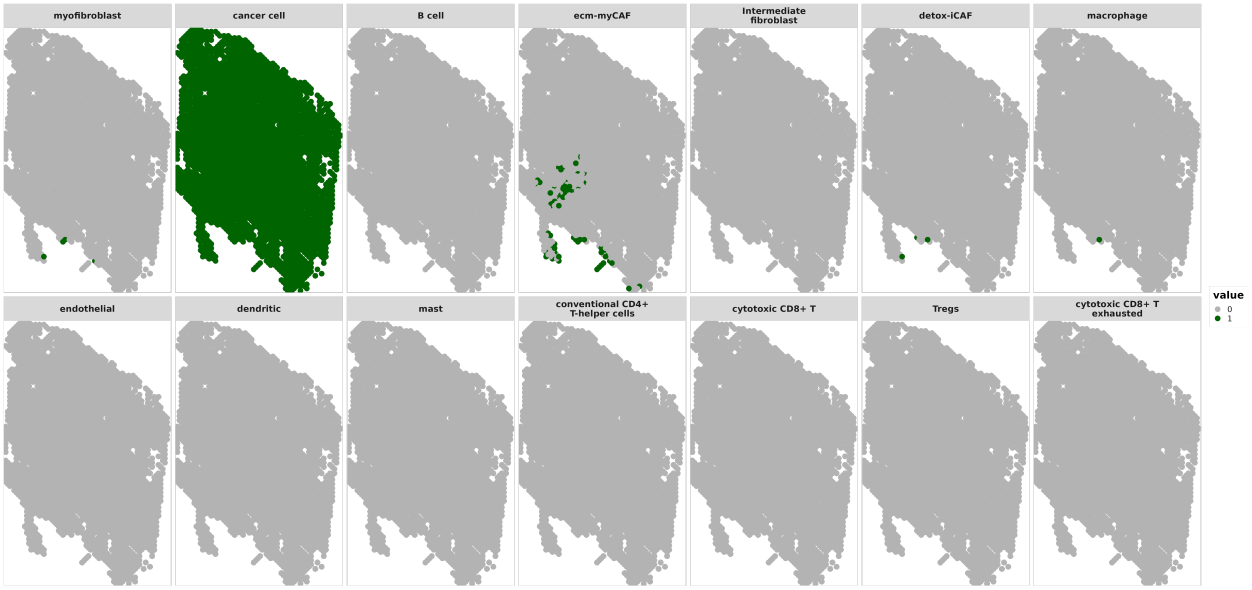


(b) Cell-Type Proportions $> 0.05$

**Figure 18**: OSCC Sample 6 CARD Cell-Type Proportions Estimates. a) Plots of cell-type proportions for all 14 cell types. b) Binary plots of cell-type proportions $> 0.05$ for all 14 cell types.

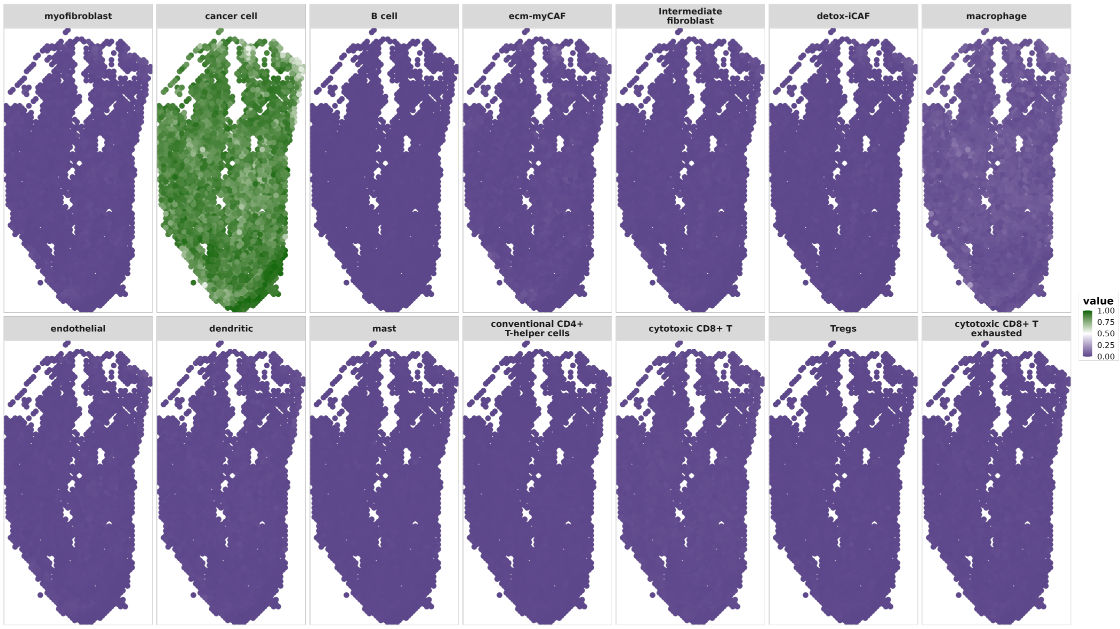


(a) Cell-Type Proportions Estimates

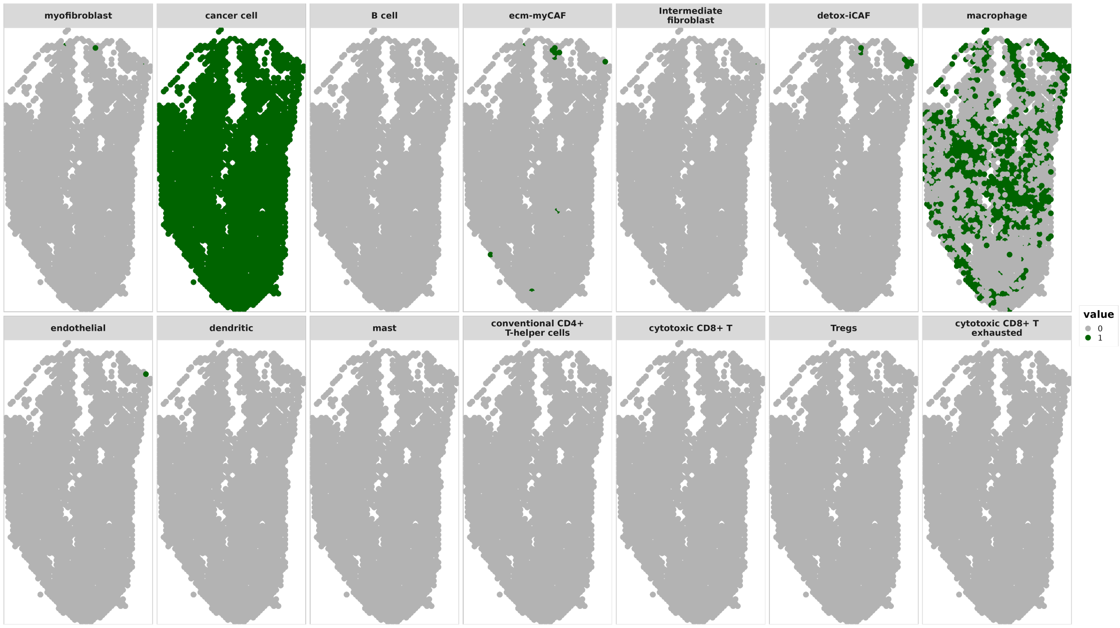


(b) Cell-Type Proportions $> 0.05$

**Figure 19**: OSCC Sample 7 ZI-HGT + CARD Cell-Type Proportions Estimates. a) Plots of cell-type proportions for all 14 cell types. b) Binary plots of cell-type proportions $> 0.05$ for all 14 cell types.

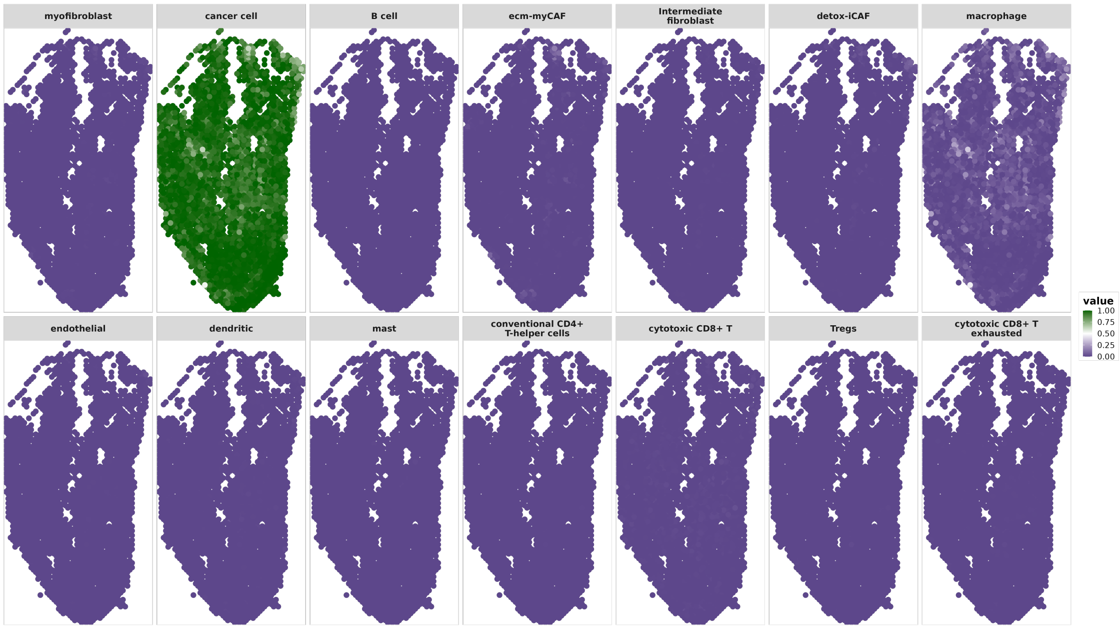


(a) Cell-Type Proportions Estimates

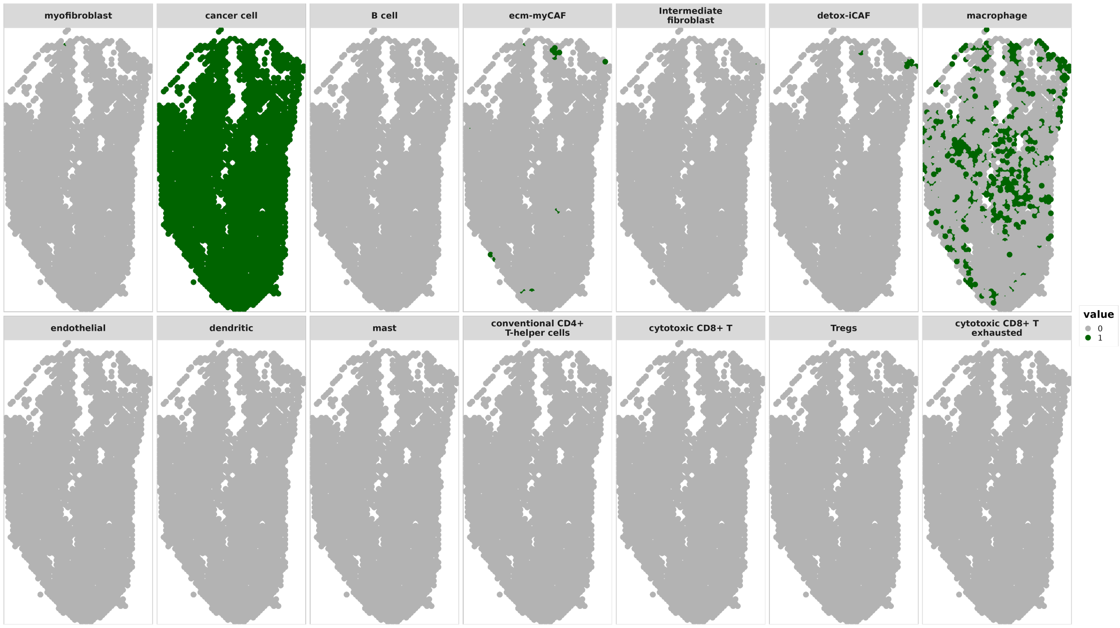


(b) Cell-Type Proportions > 0.05

**Figure 20**: OSCC Sample 7 CARD Cell-Type Proportions Estimates. a) Plots of cell-type proportions for all 14 cell types. b) Binary plots of cell-type proportions > 0.05 for all 14 cell types.

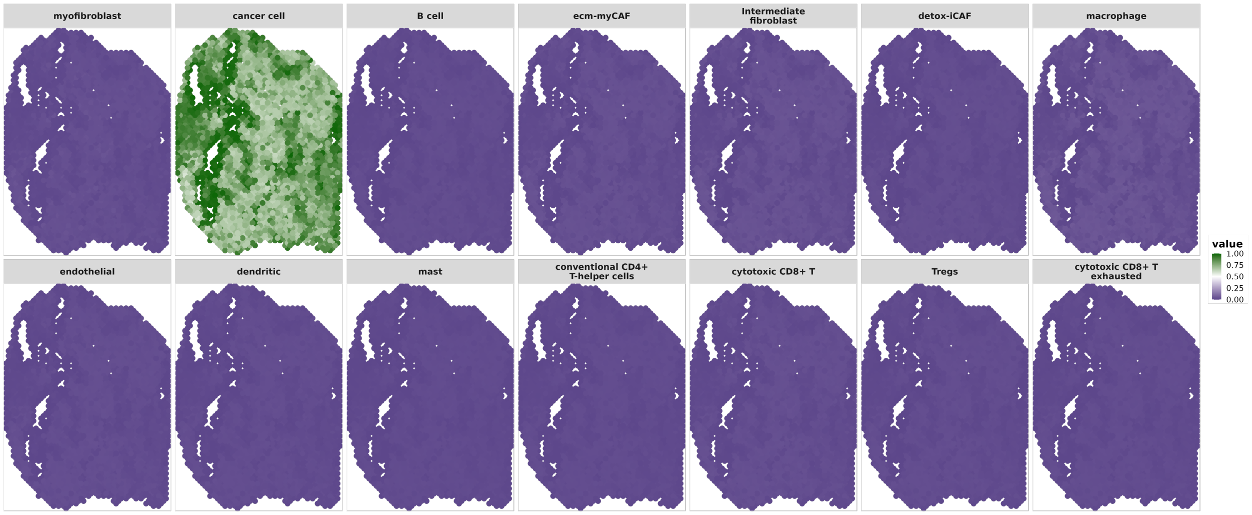


(a) Cell-Type Proportions Estimates

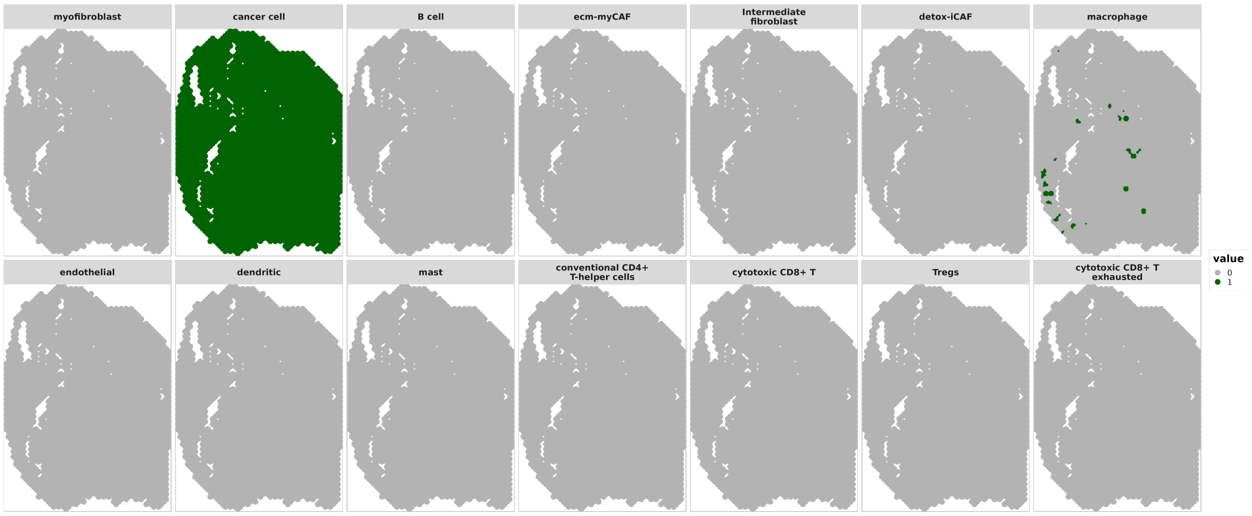


(b) Cell-Type Proportions > 0.05

**Figure 21**: OSCC Sample 8 ZI-HGT + CARD Cell-Type Proportions Estimates. a) Plots of cell-type proportions for all 14 cell types. b) Binary plots of cell-type proportions > 0.05 for all 14 cell types.

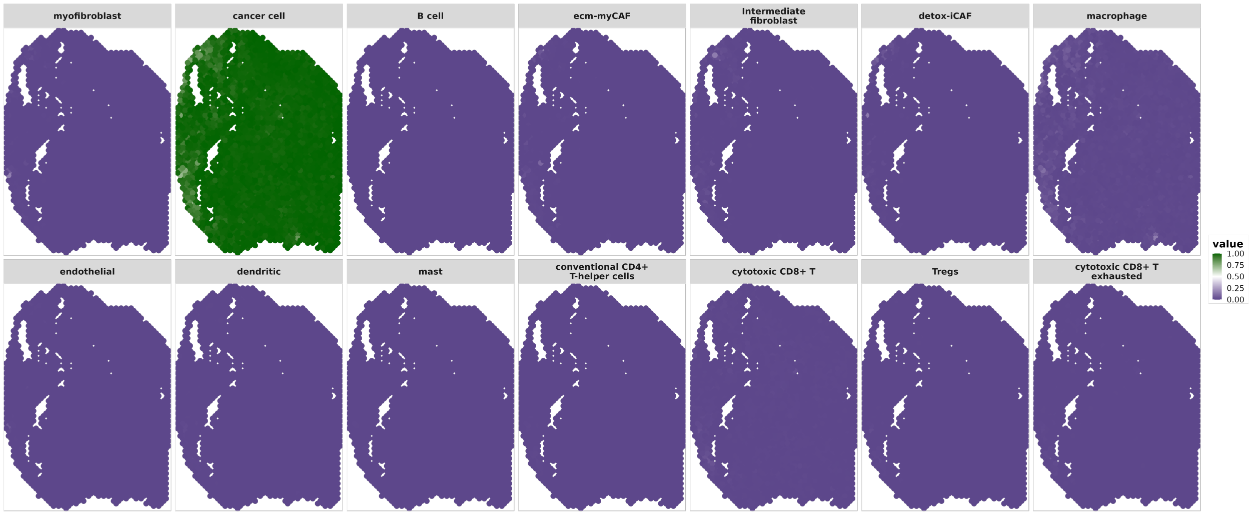


(a) Cell-Type Proportions Estimates

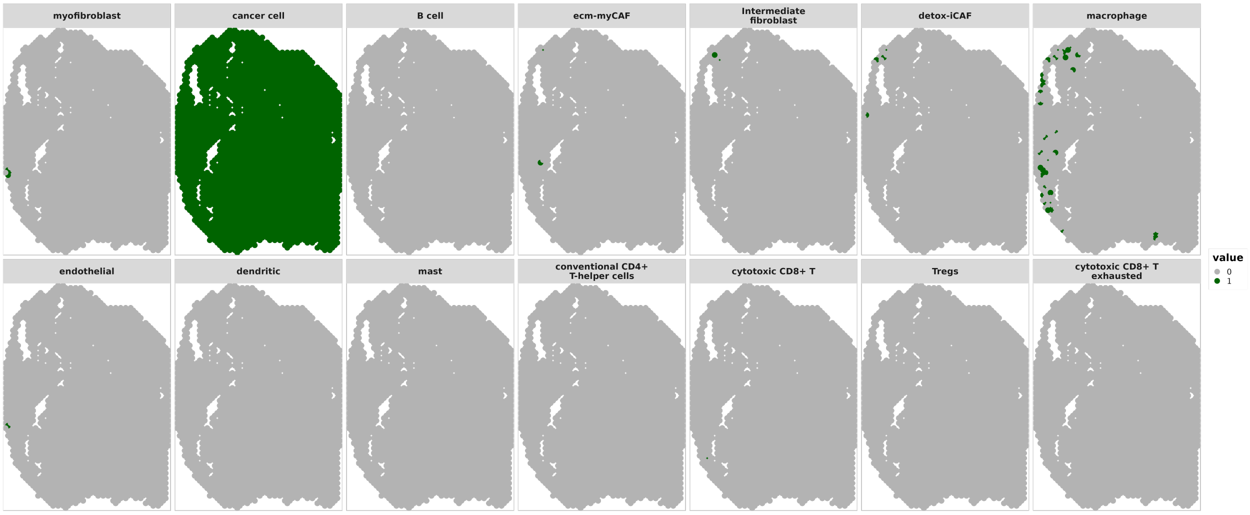


(b) Cell-Type Proportions $> 0.05$

**Figure 22**: OSCC Sample 8 CARD Cell-Type Proportions Estimates. a) Plots of cell-type proportions for all 14 cell types. b) Binary plots of cell-type proportions $> 0.05$ for all 14 cell types.

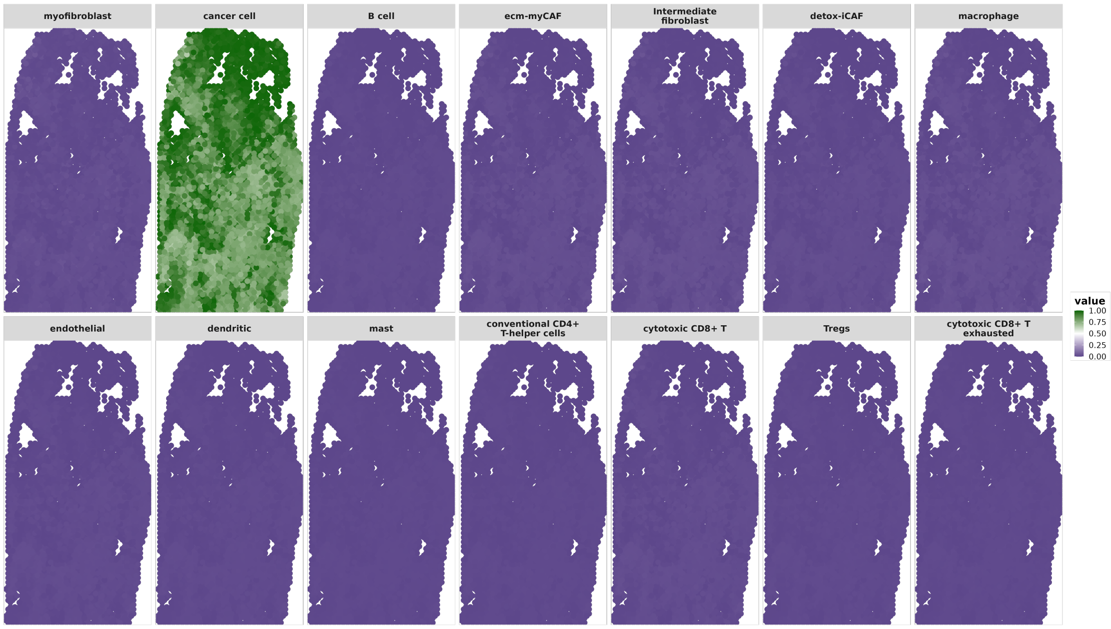


(a) Cell-Type Proportions Estimates

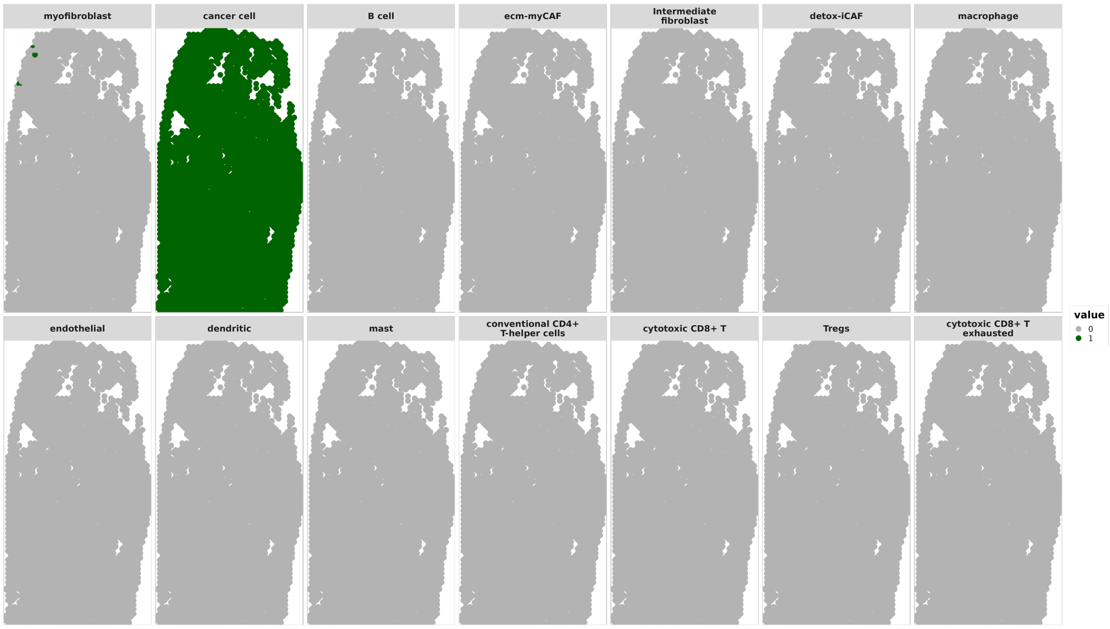


(b) Cell-Type Proportions $> 0.05$

**Figure 23**: OSCC Sample 9 ZI-HGT + CARD Cell-Type Proportions Estimates. a) Plots of cell-type proportions for all 14 cell types. b) Binary plots of cell-type proportions $> 0.05$ for all 14 cell types.

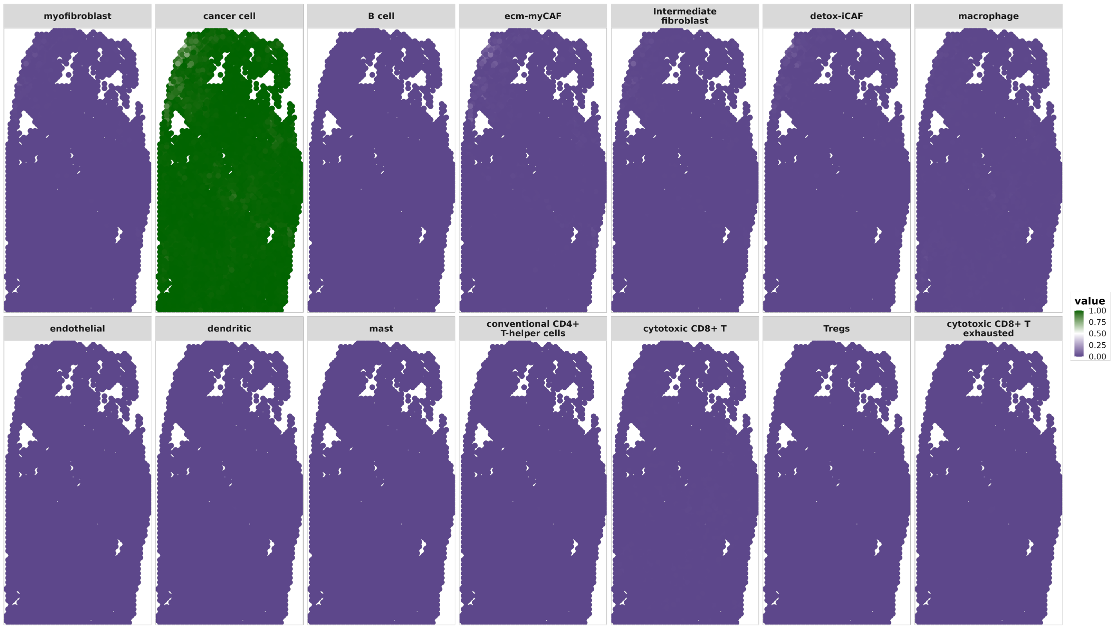


(a) Cell-Type Proportions Estimates

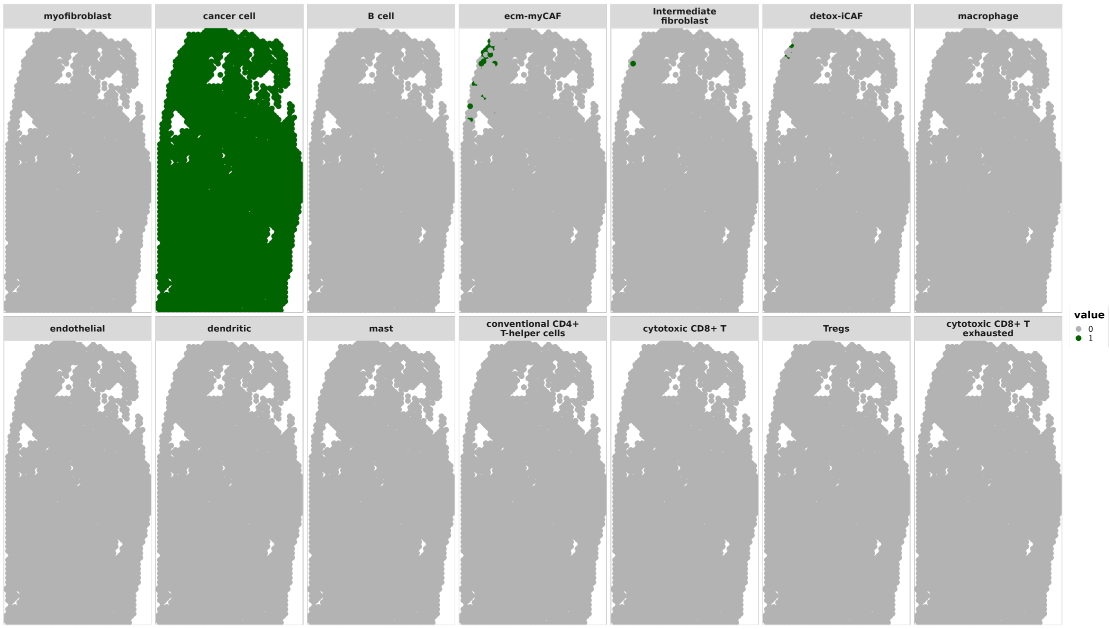


(b) Cell-Type Proportions $> 0.05$

**Figure 24**: OSCC Sample 9 CARD Cell-Type Proportions Estimates. a) Plots of cell-type proportions for all 14 cell types. b) Binary plots of cell-type proportions $> 0.05$ for all 14 cell types.

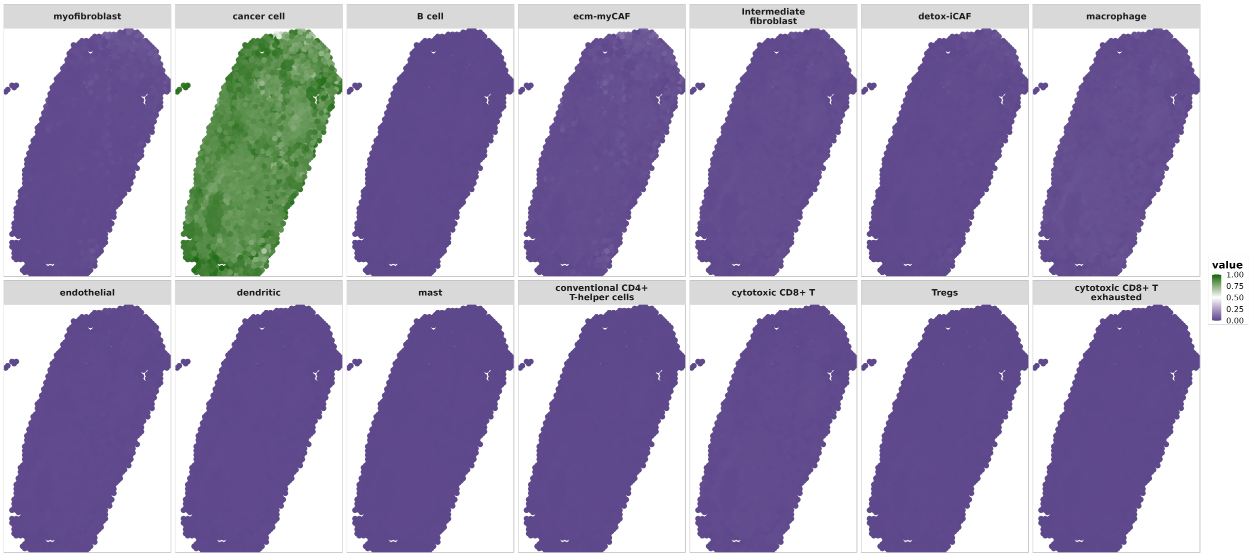


(a) Cell-Type Proportions Estimates

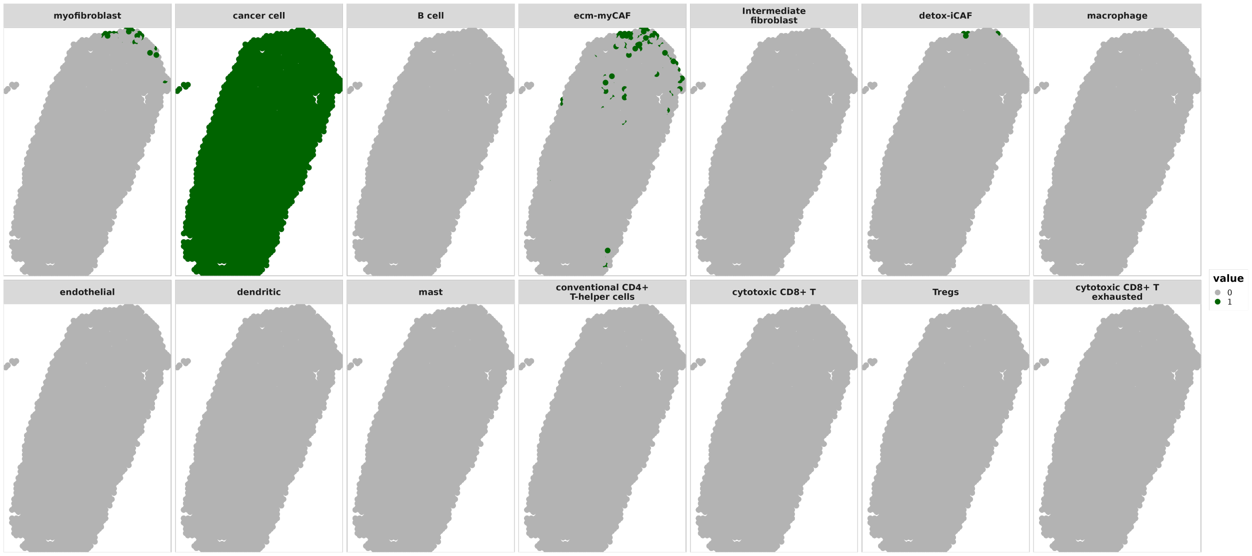


(b) Cell-Type Proportions $> 0.05$

**Figure 25**: OSCC Sample 10 ZI-HGT + CARD Cell-Type Proportions Estimates. a) Plots of cell-type proportions for all 14 cell types. b) Binary plots of cell-type proportions $> 0.05$ for all 14 cell types.

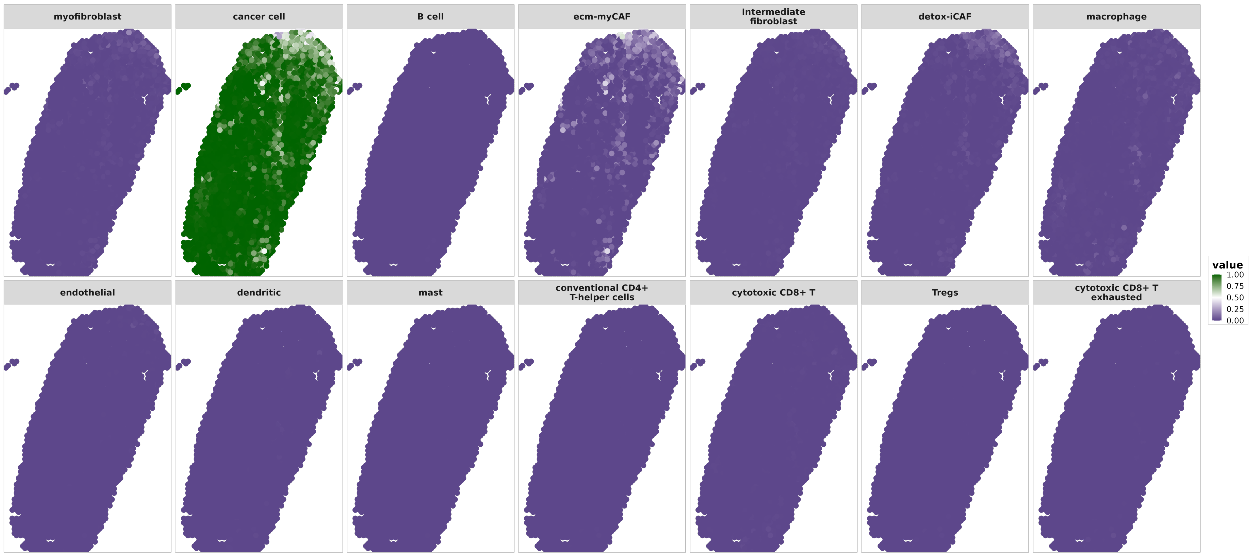


(a) Cell-Type Proportions Estimates

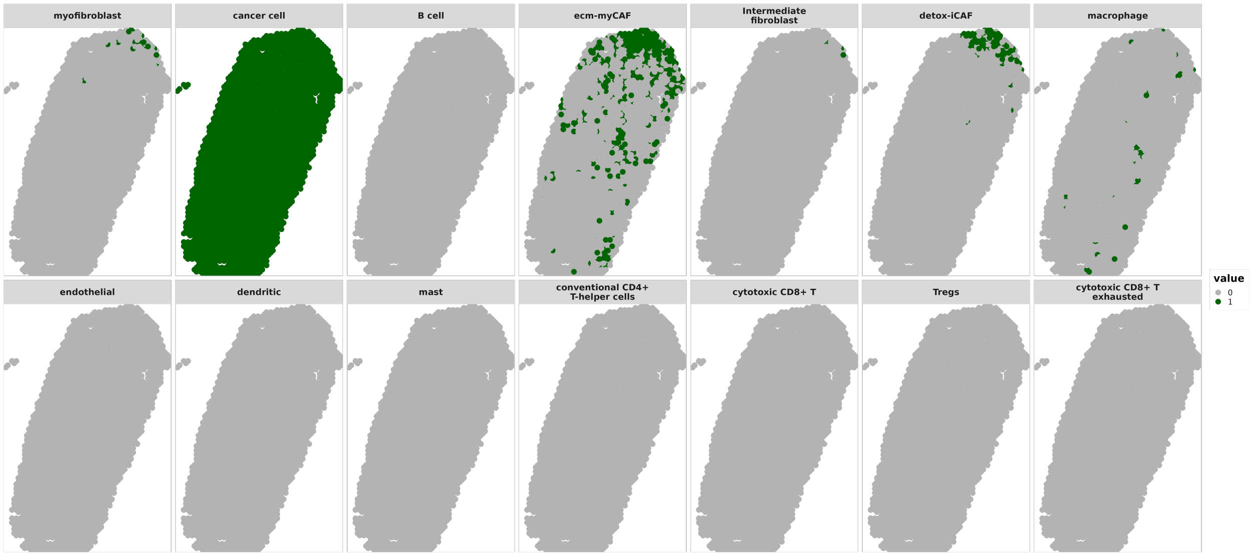


(b) Cell-Type Proportions $> 0.05$

**Figure 26**: OSCC Sample 10 CARD Cell-Type Proportions Estimates. a) Plots of cell-type proportions for all 14 cell types. b) Binary plots of cell-type proportions $> 0.05$ for all 14 cell types.

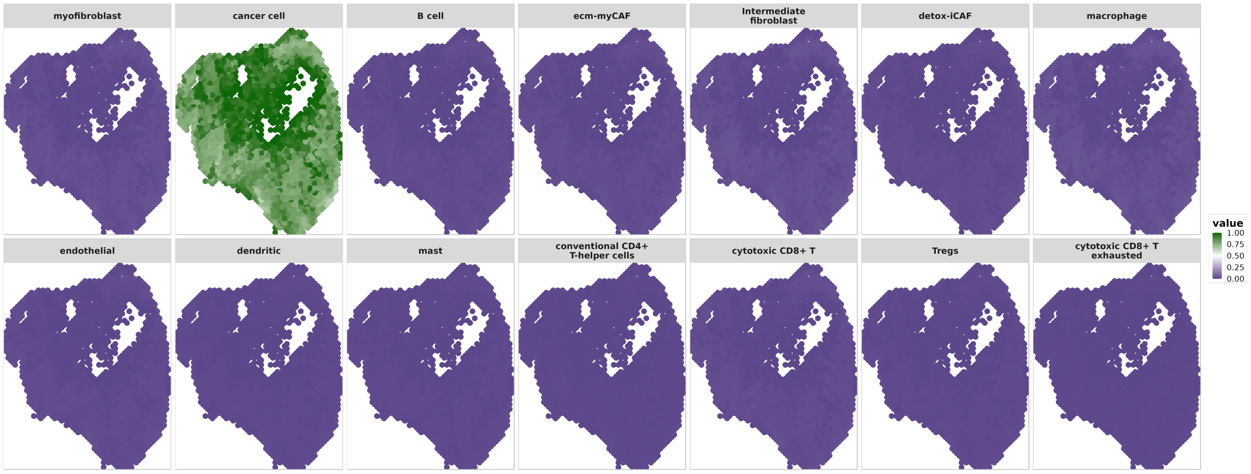


(a) Cell-Type Proportions Estimates

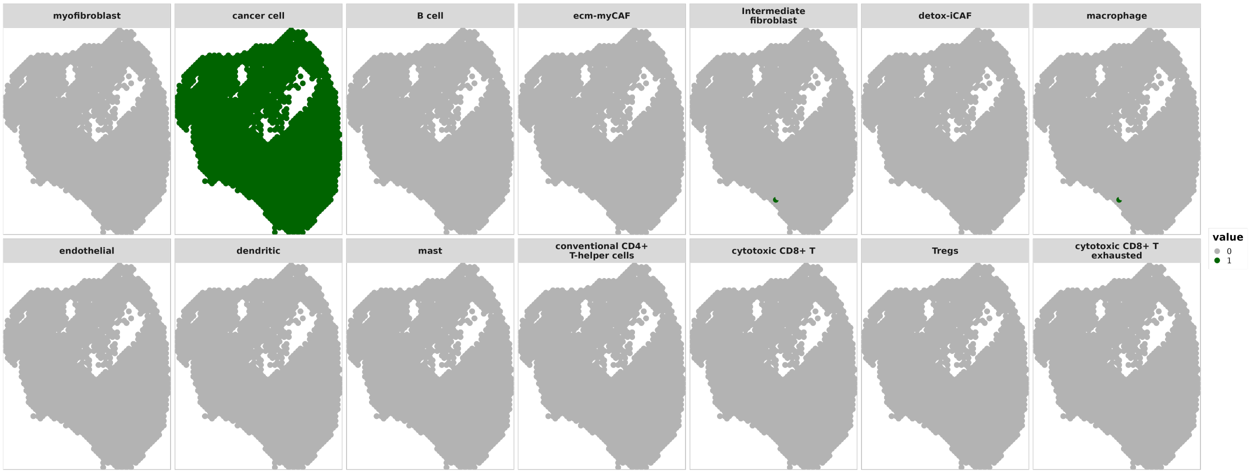


(b) Cell-Type Proportions $> 0.05$

**Figure 27**: OSCC Sample 11 ZI-HGT + CARD Cell-Type Proportions Estimates. a) Plots of cell-type proportions for all 14 cell types. b) Binary plots of cell-type proportions $> 0.05$ for all 14 cell types.

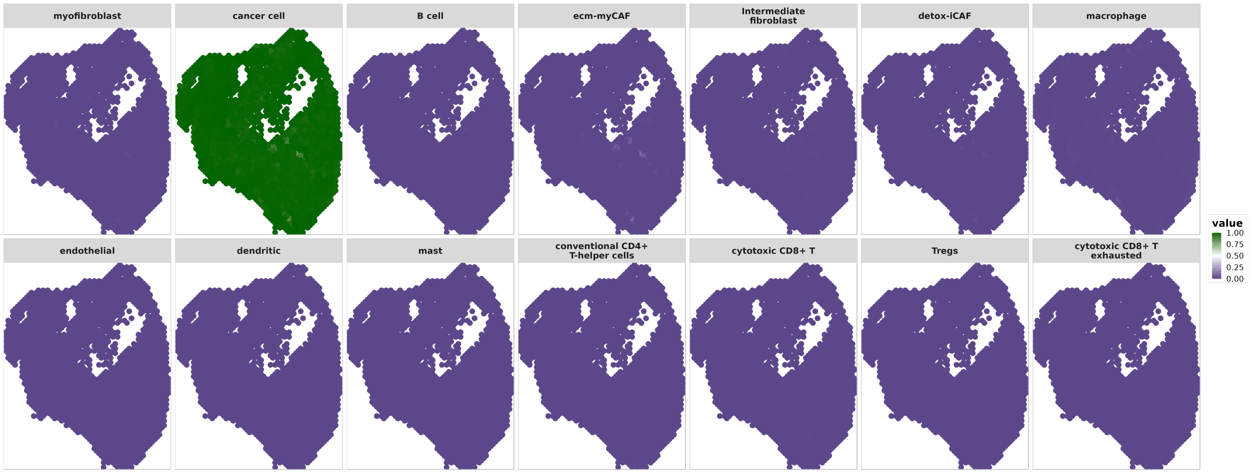


(a) Cell-Type Proportions Estimates

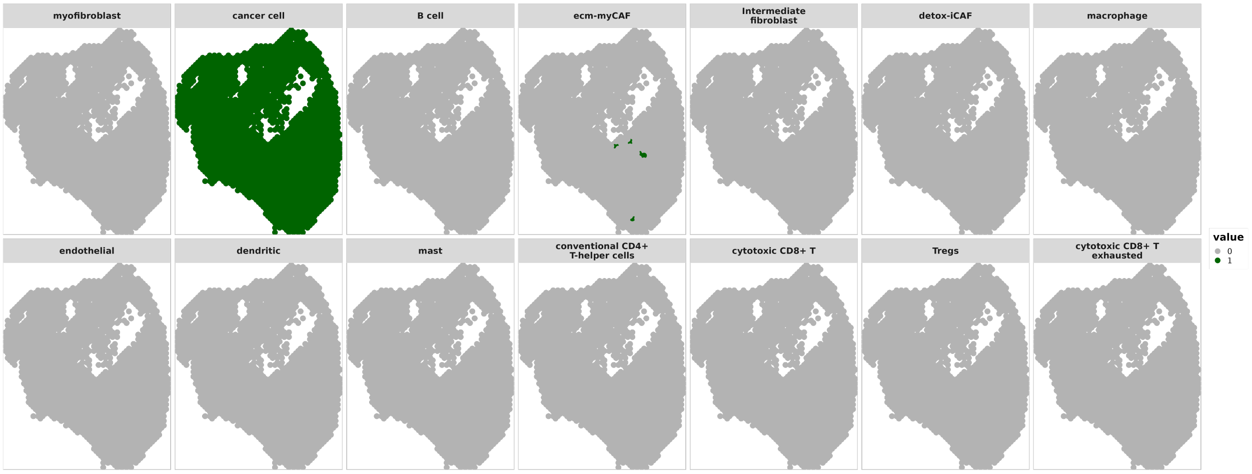


(b) Cell-Type Proportions > 0.05

**Figure 28**: OSCC Sample 11 CARD Cell-Type Proportions Estimates. a) Plots of cell-type proportions for all 14 cell types. b) Binary plots of cell-type proportions > 0.05 for all 14 cell types.

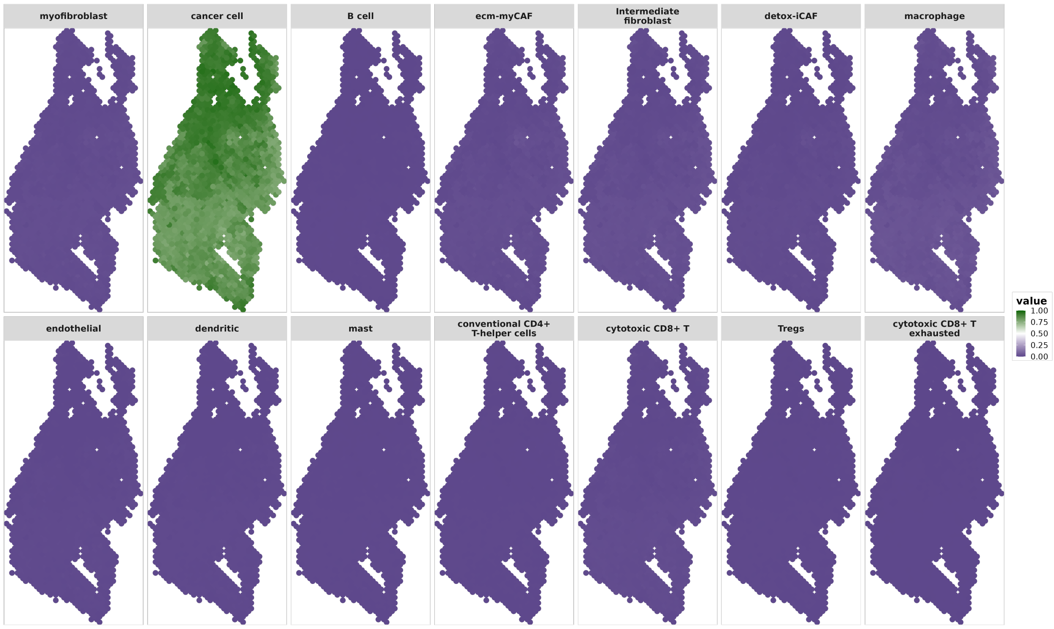


(a) Cell-Type Proportions Estimates

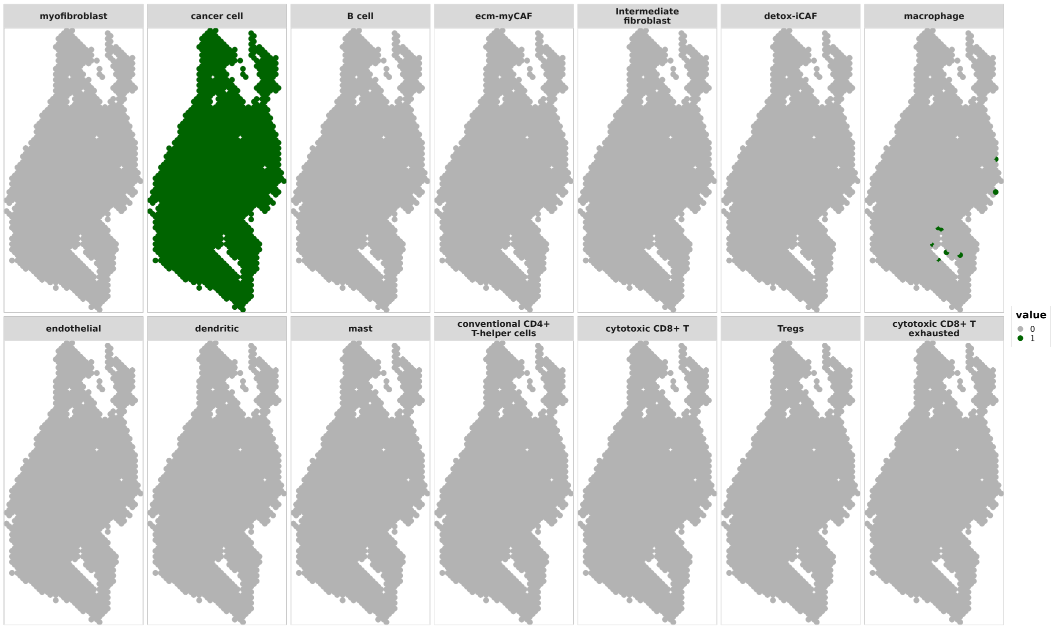


(b) Cell-Type Proportions $> 0.05$

**Figure 29**: OSCC Sample 12 ZI-HGT + CARD Cell-Type Proportions Estimates. a) Plots of cell-type proportions for all 14 cell types. b) Binary plots of cell-type proportions $> 0.05$ for all 14 cell types.

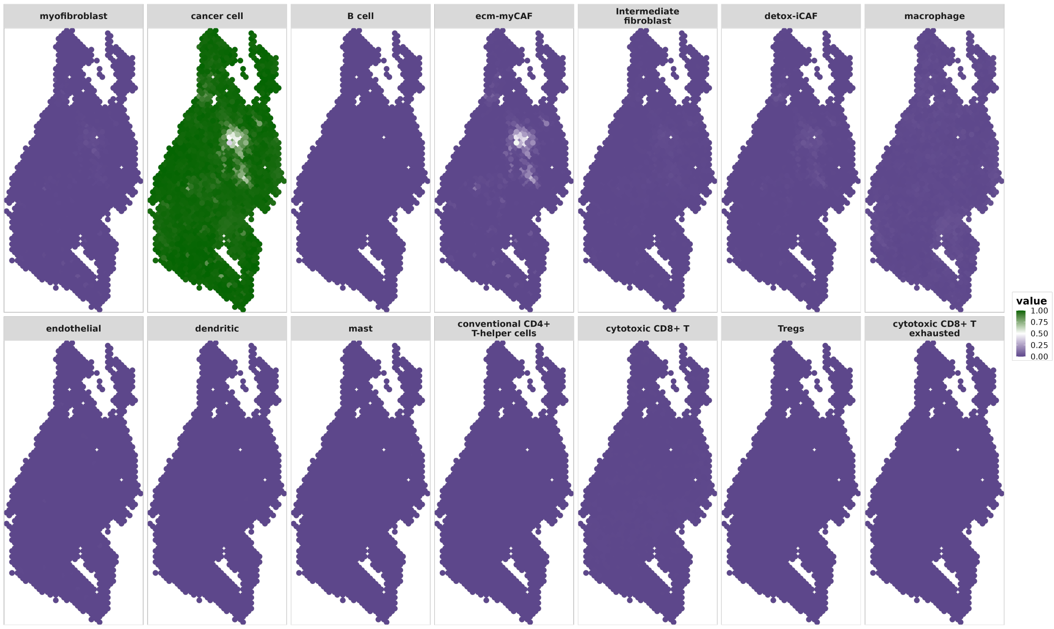


(a) Cell-Type Proportions Estimates

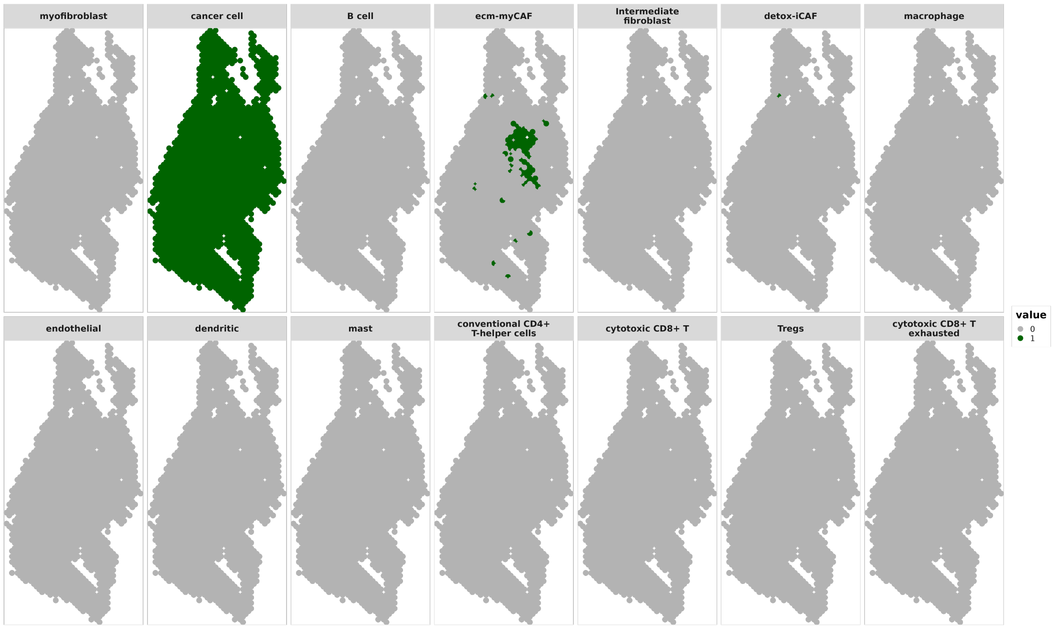


(b) Cell-Type Proportions $> 0.05$

**Figure 30**: OSCC Sample 12 CARD Cell-Type Proportions Estimates. a) Plots of cell-type proportions for all 14 cell types. b) Binary plots of cell-type proportions $> 0.05$ for all 14 cell types.

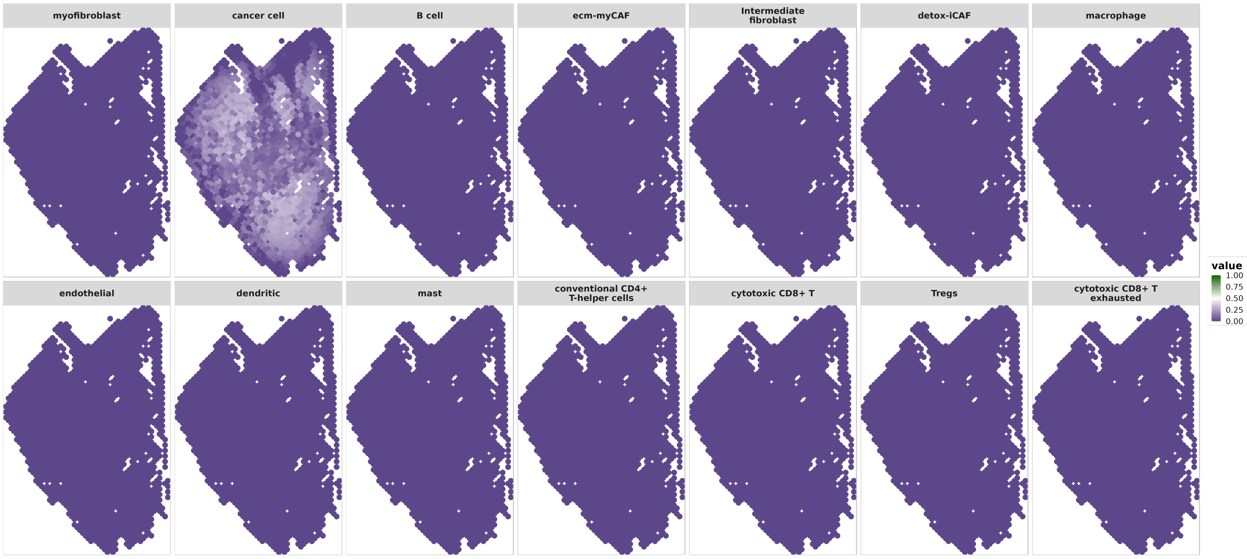


(a) Lower Bound

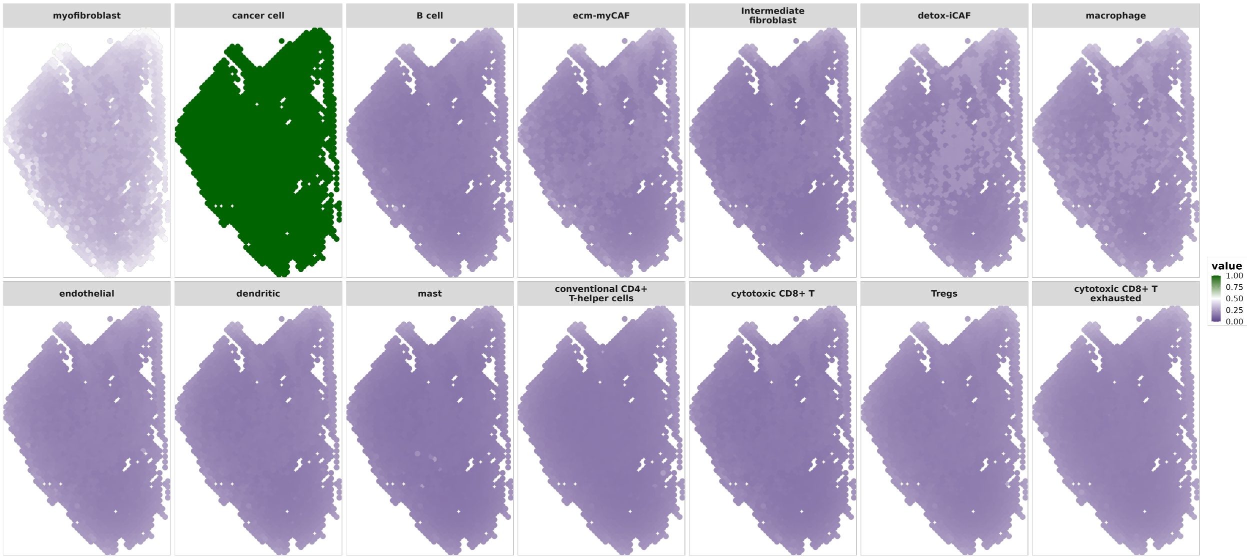


(b) Upper Bound

**Figure 31**: OSCC Sample 2 Pointwise 95% Credible Intervals. Pointwise 95% credible intervals for all cell-type proportions at all locations.

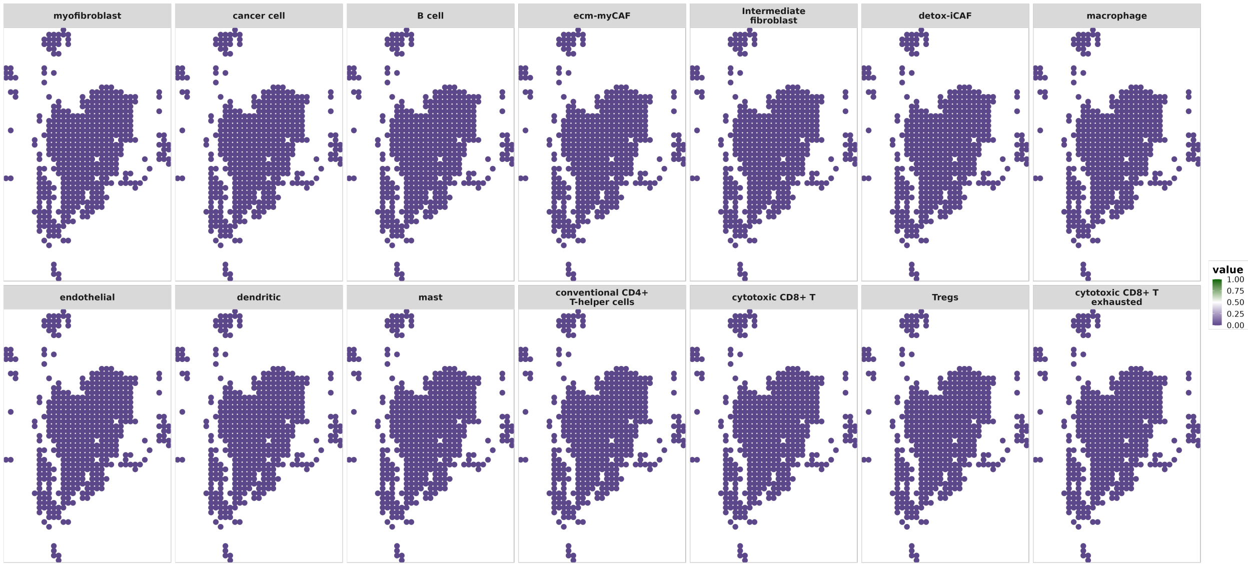


(a) Lower Bound

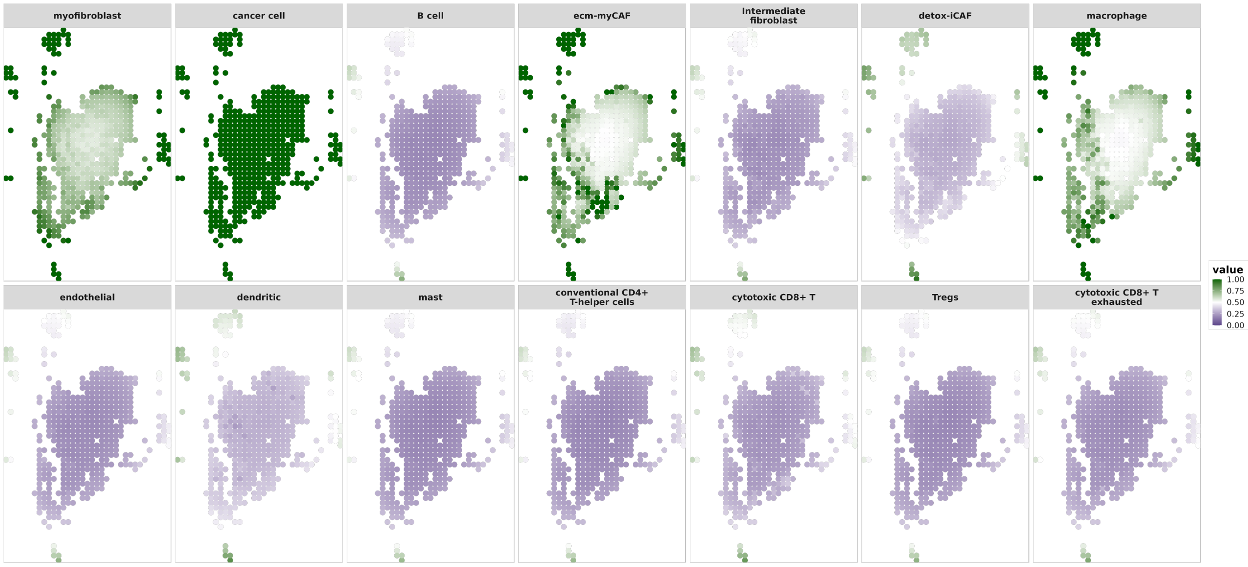


(b) Upper Bound

**Figure 32**: OSCC Sample 3 Pointwise 95% Credible Intervals. Pointwise 95% credible intervals for all cell-type proportions at all locations.

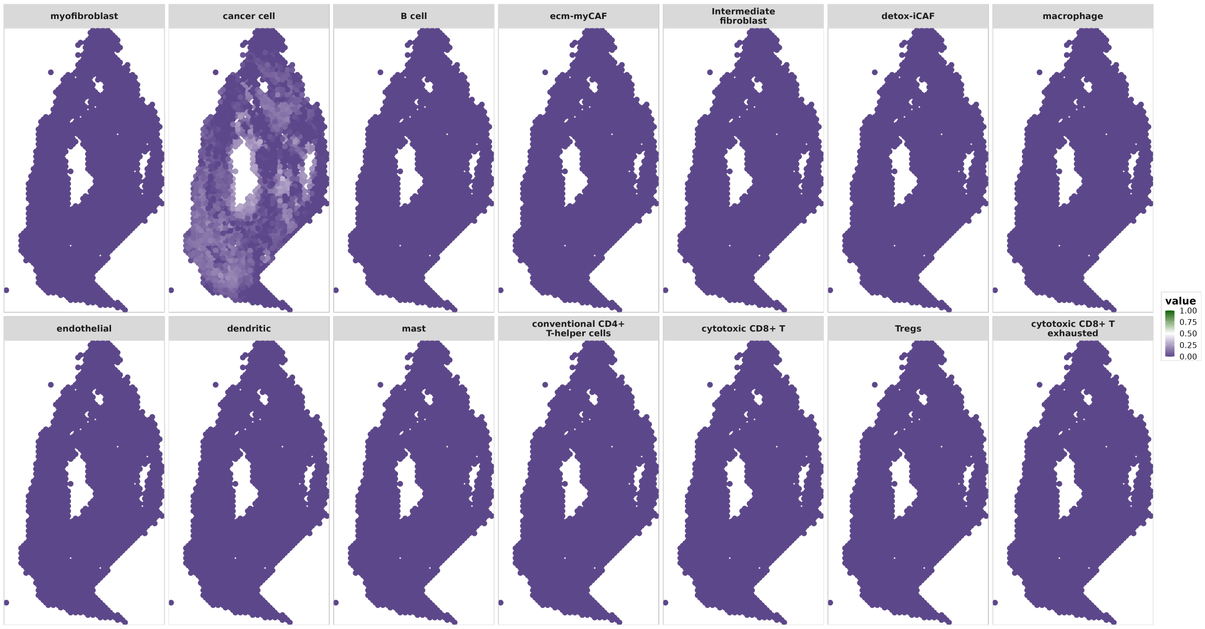


(a) Lower Bound

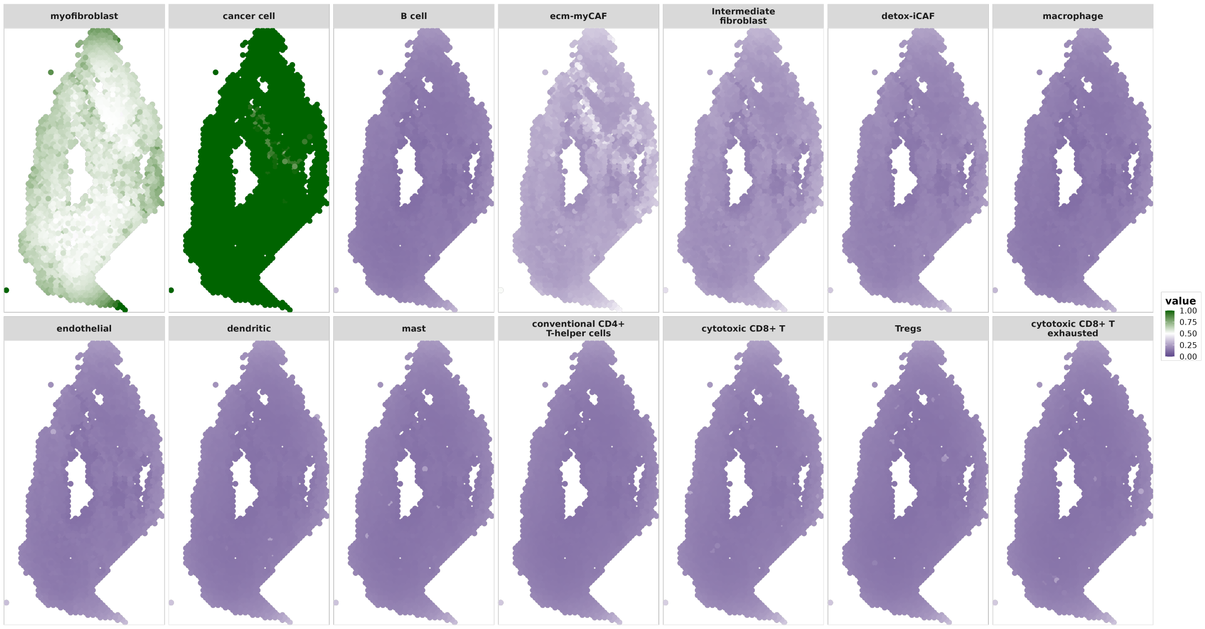


(b) Upper Bound

**Figure 33**: OSCC Sample 4 Pointwise 95% Credible Intervals. Pointwise 95% credible intervals for all cell-type proportions at all locations.

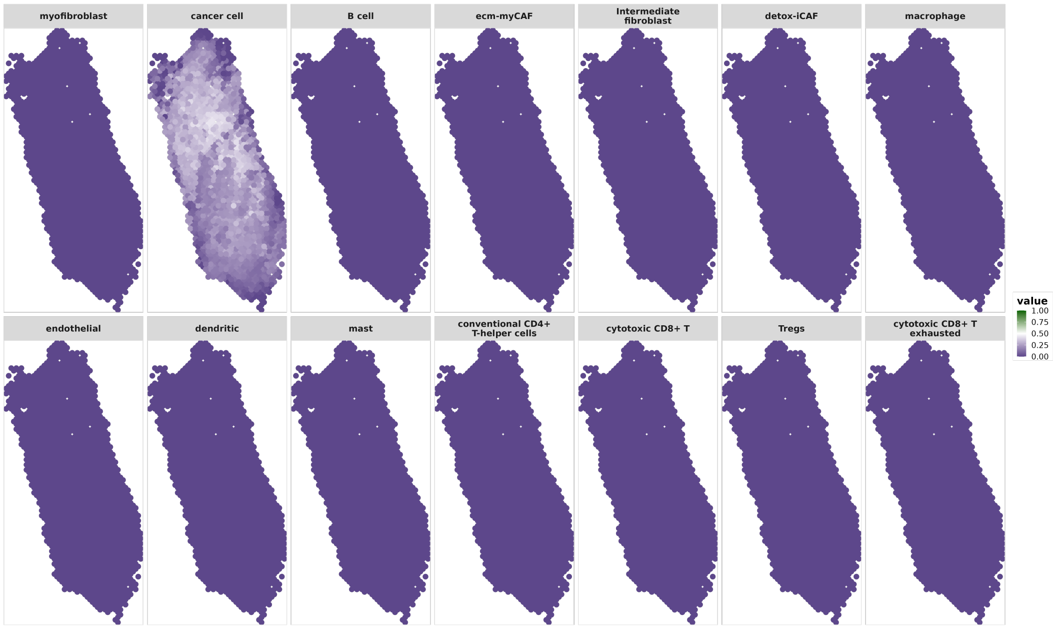


(a) Lower Bound

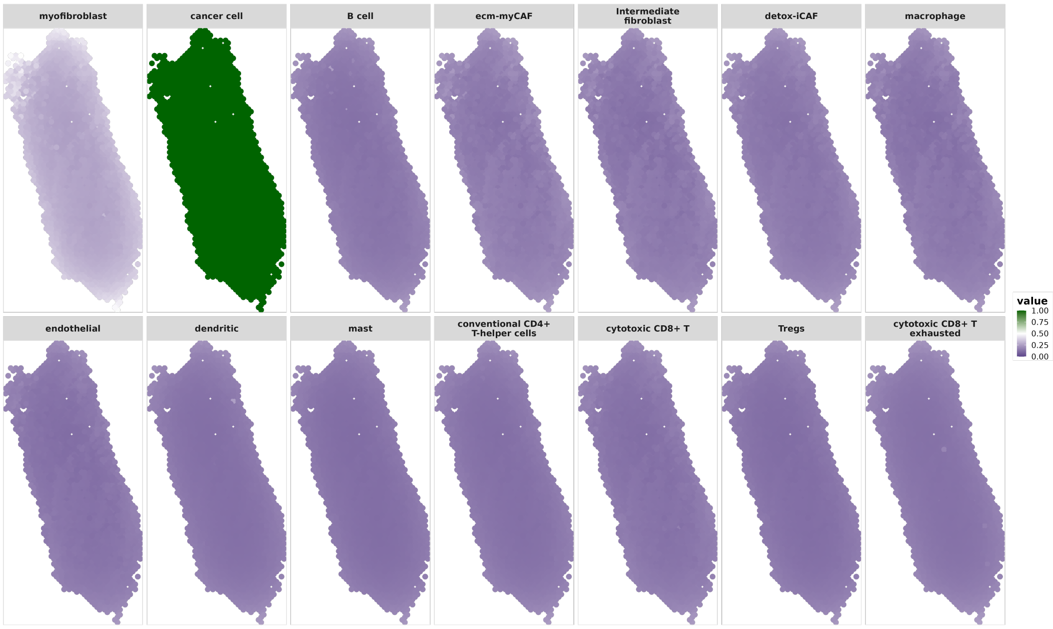


(b) Upper Bound

**Figure 34**: OSCC Sample 5 Pointwise 95% Credible Intervals. Pointwise 95% credible intervals for all cell-type proportions at all locations.

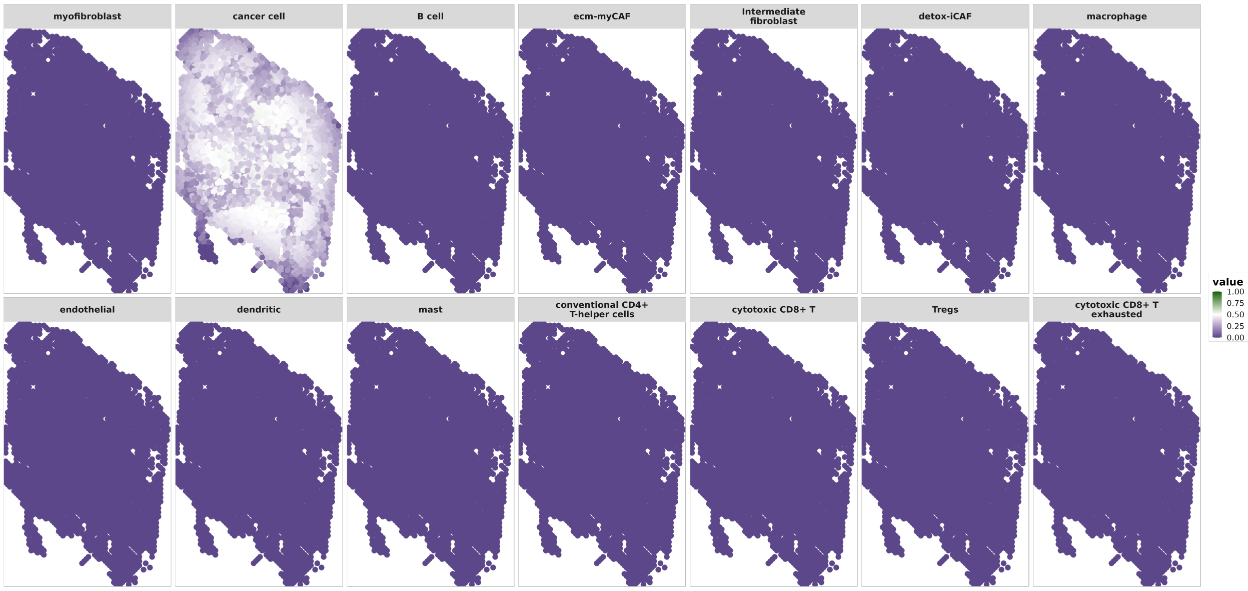


(a) Lower Bound

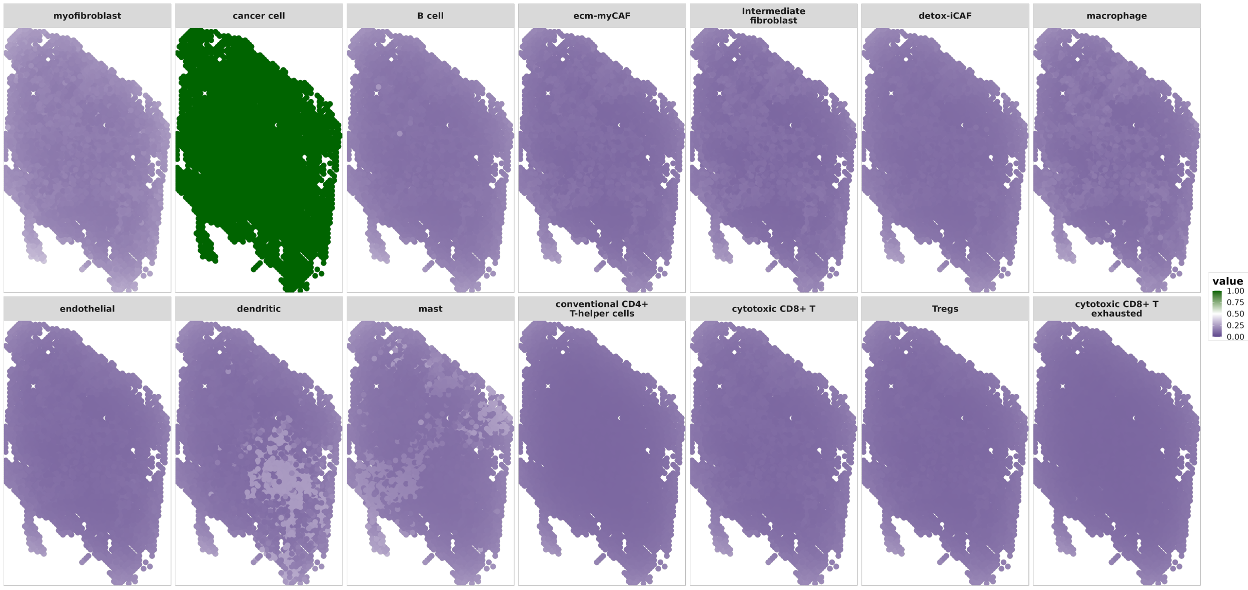


(b) Upper Bound

**Figure 35**: OSCC Sample 6 Pointwise 95% Credible Intervals. Pointwise 95% credible intervals for all cell-type proportions at all locations.

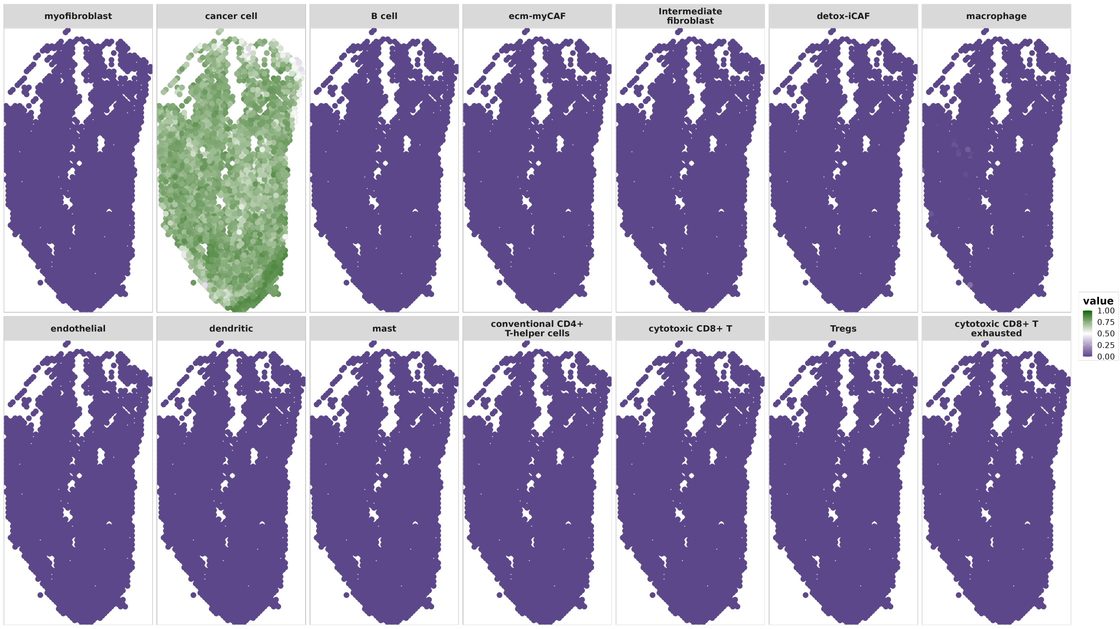


(a) Lower Bound

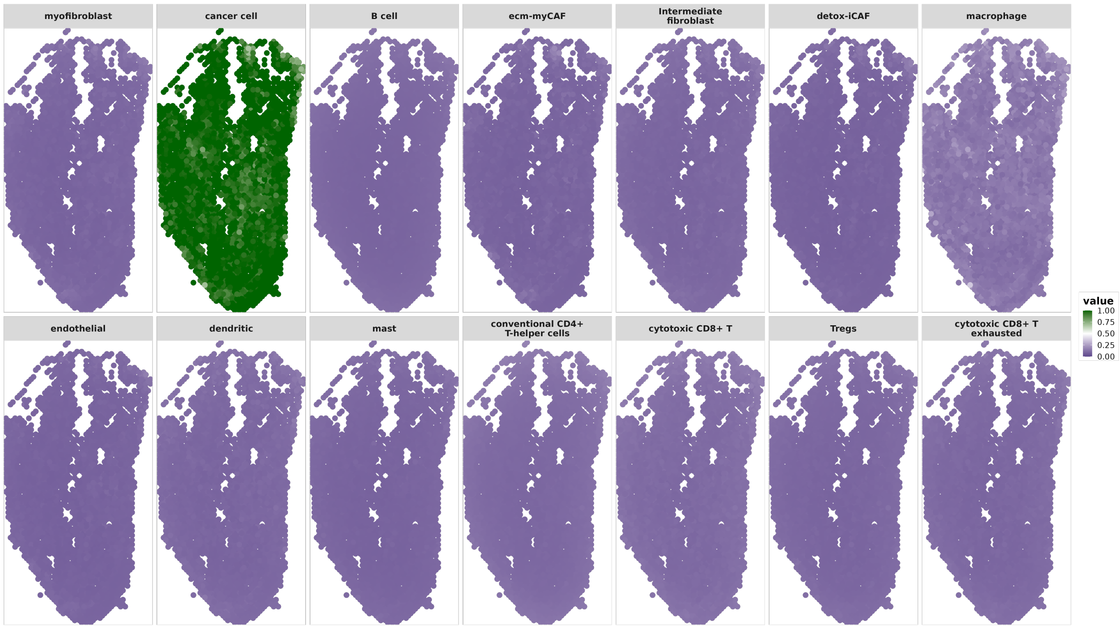


(b) Upper Bound

**Figure 36**: OSCC Sample 7 Pointwise 95% Credible Intervals. Pointwise 95% credible intervals for all cell-type proportions at all locations.

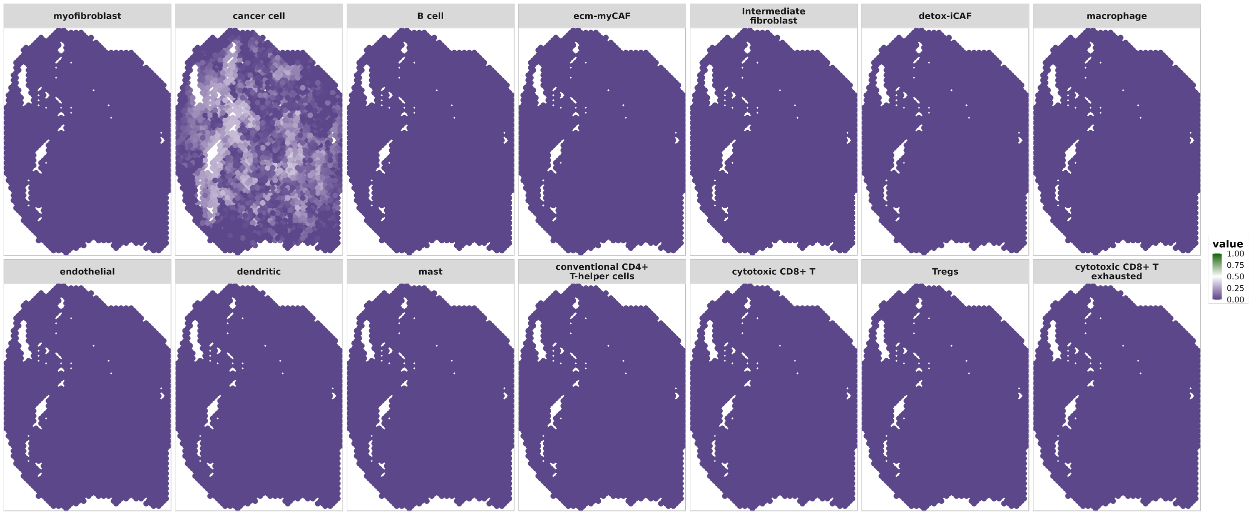


(a) Lower Bound

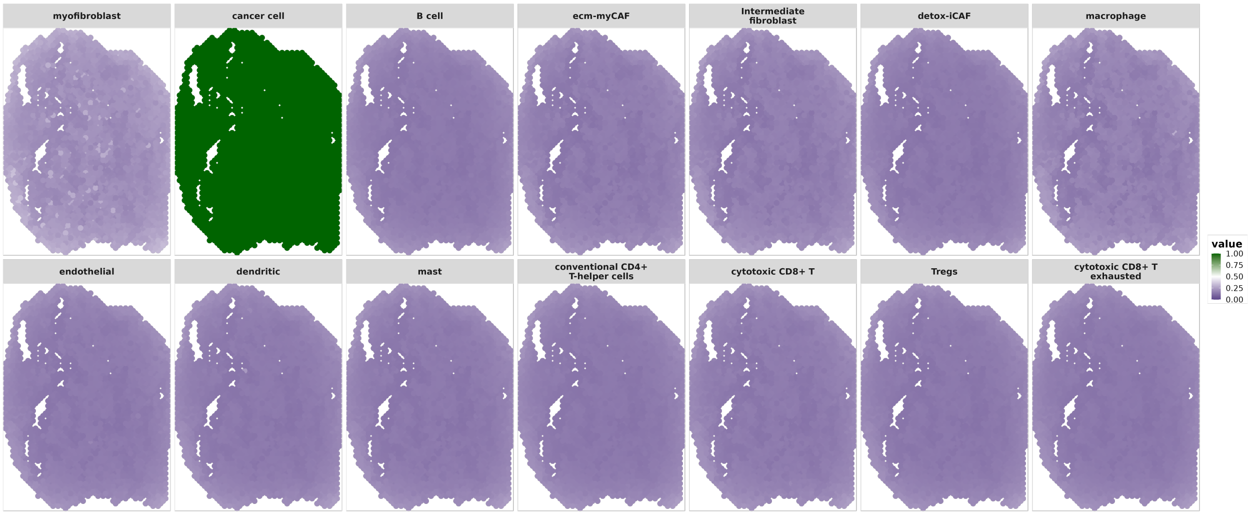


(b) Upper Bound

**Figure 37**: OSCC Sample 8 Pointwise 95% Credible Intervals. Pointwise 95% credible intervals for all cell-type proportions at all locations.

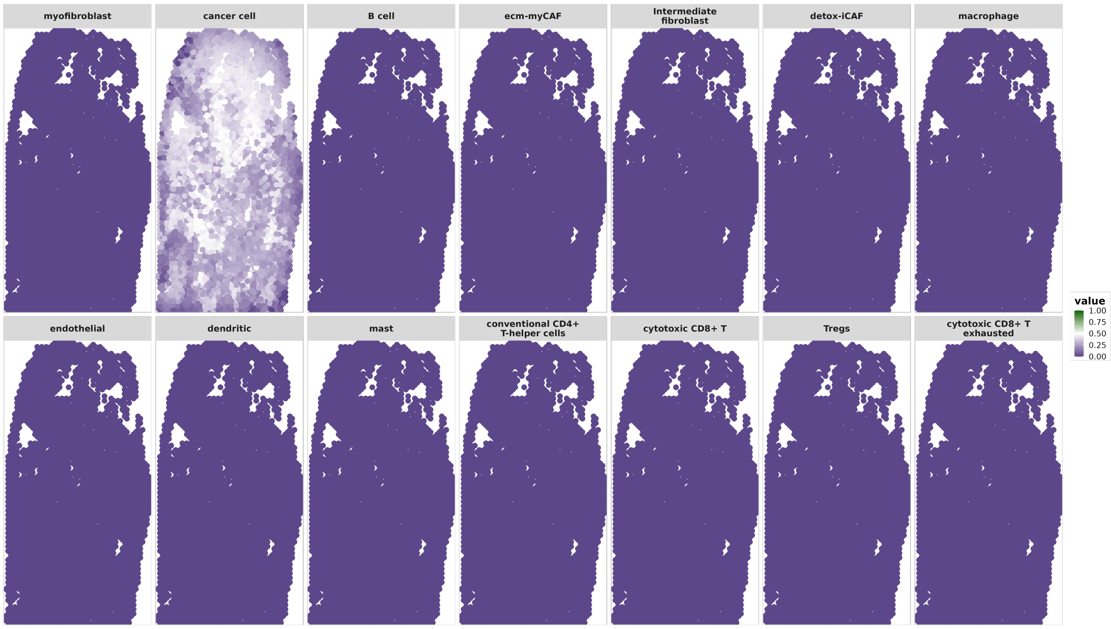


(a) Lower Bound

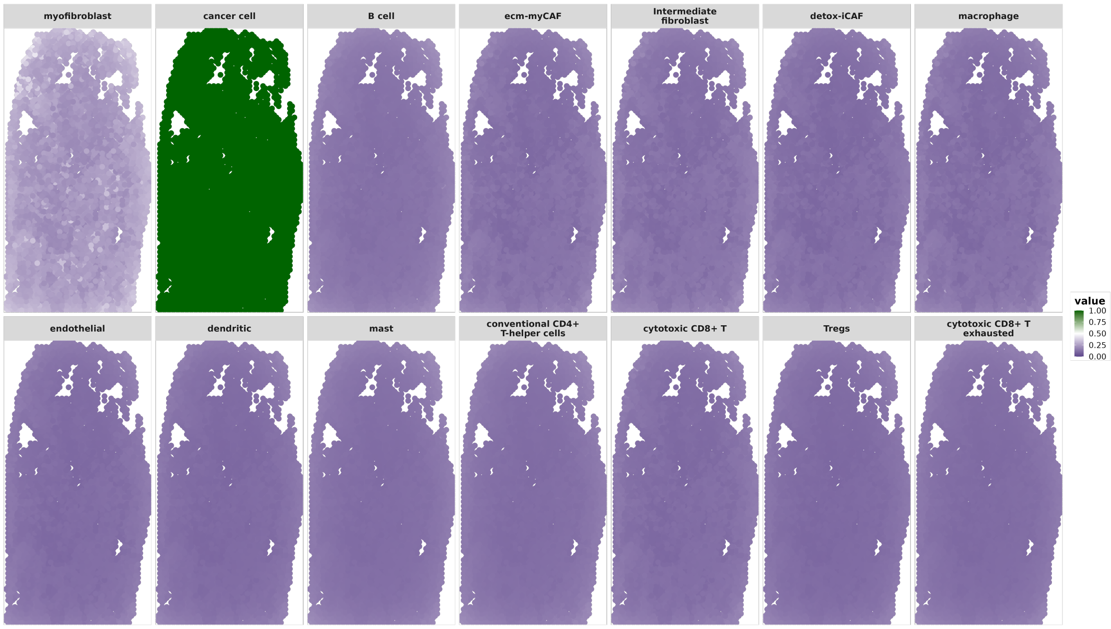


(b) Upper Bound

**Figure 38**: OSCC Sample 9 Pointwise 95% Credible Intervals. Pointwise 95% credible intervals for all cell-type proportions at all locations.

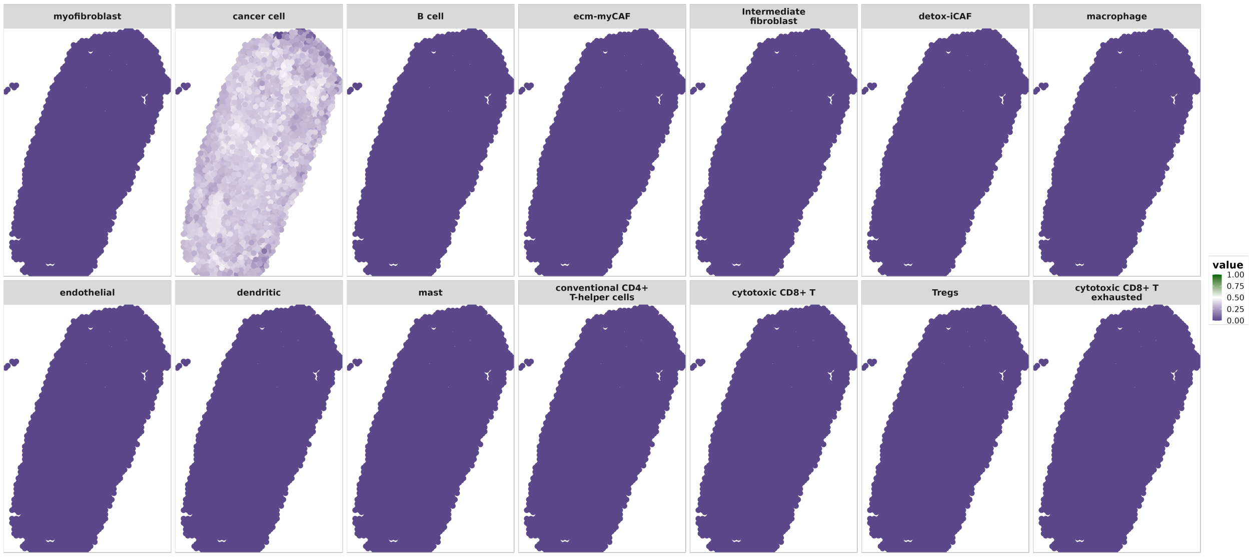


(a) Lower Bound

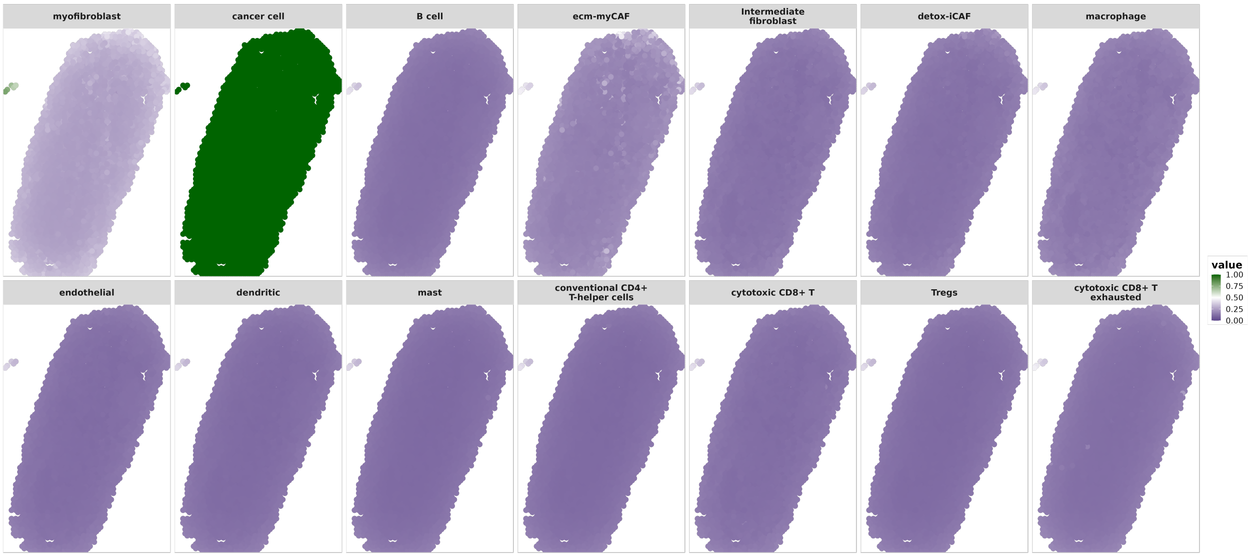


(b) Upper Bound

**Figure 39**: OSCC Sample 10 Pointwise 95% Credible Intervals. Pointwise 95% credible intervals for all cell-type proportions at all locations.

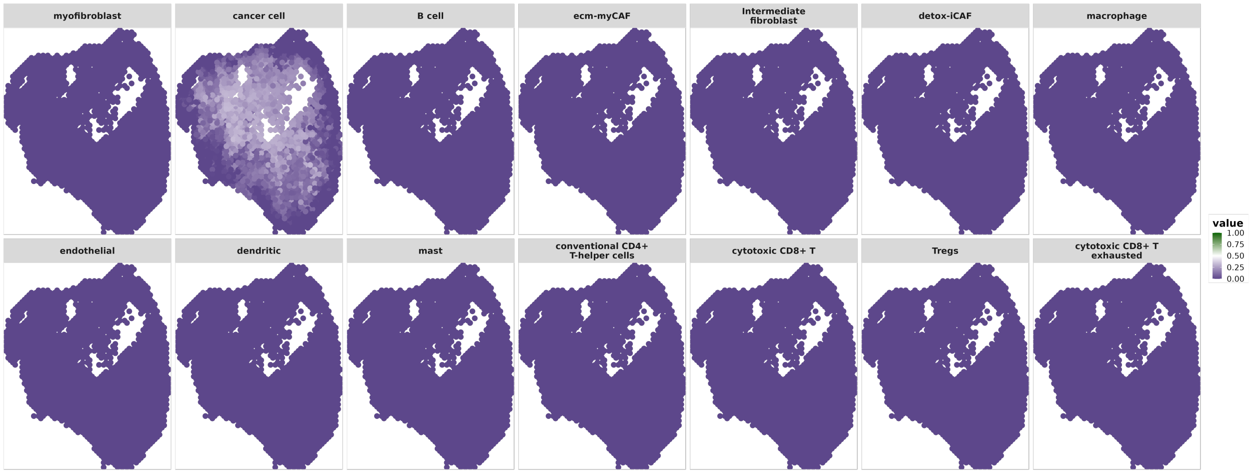


(a) Lower Bound

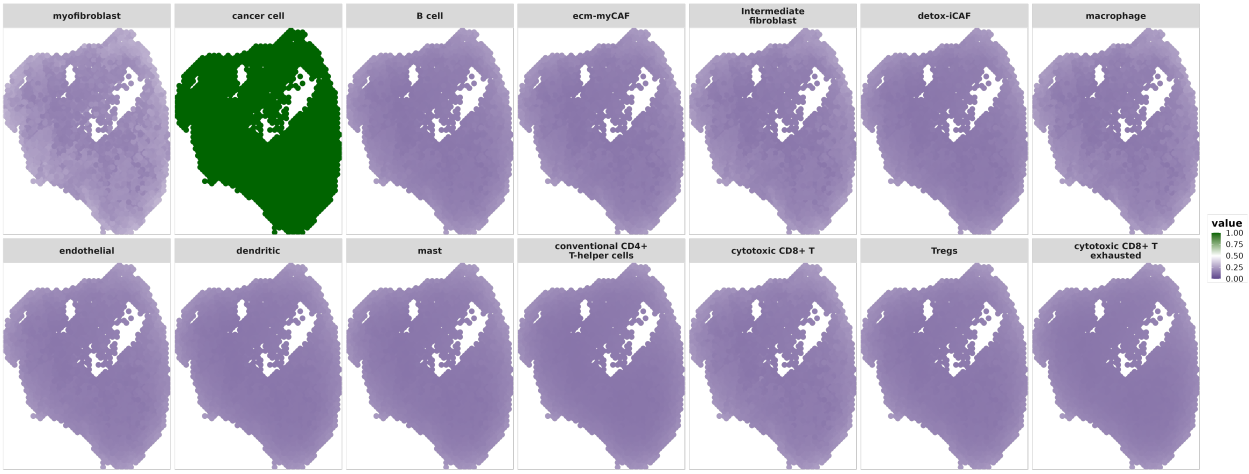


(b) Upper Bound

**Figure 40**: OSCC Sample 11 Pointwise 95% Credible Intervals. Pointwise 95% credible intervals for all cell-type proportions at all locations.

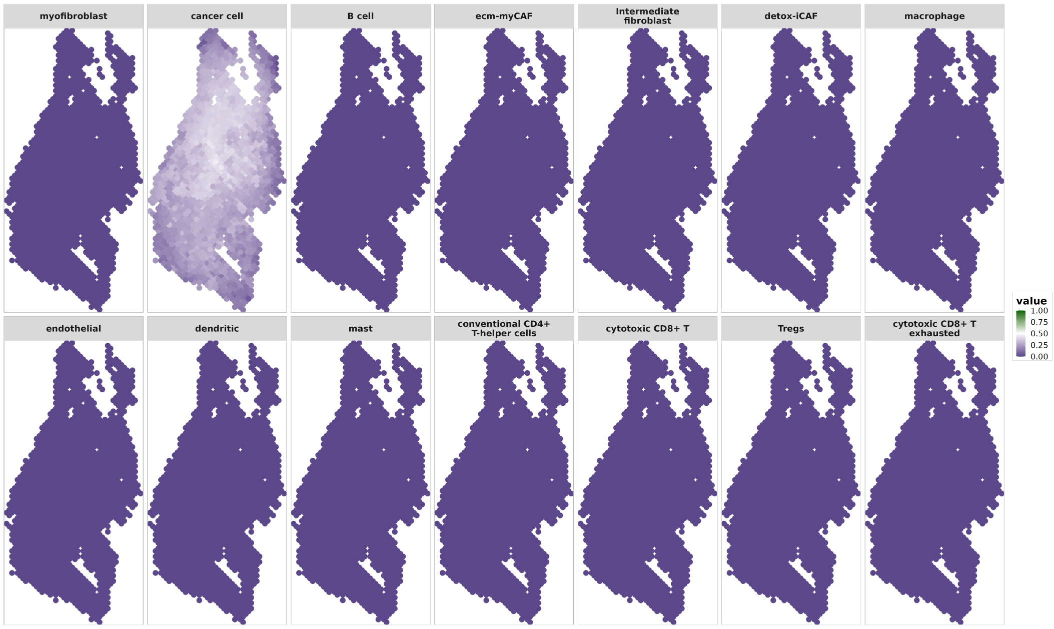


(a) Lower Bound

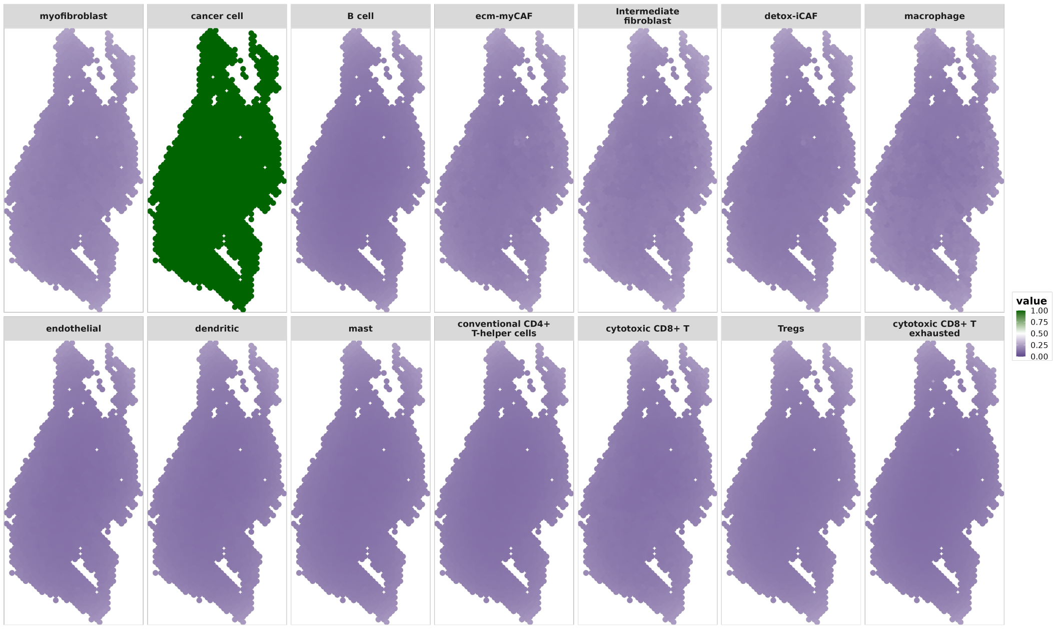


(b) Upper Bound

**Figure 41**: OSCC Sample 12 Pointwise 95% Credible Intervals. Pointwise 95% credible intervals for all cell-type proportions at all locations.

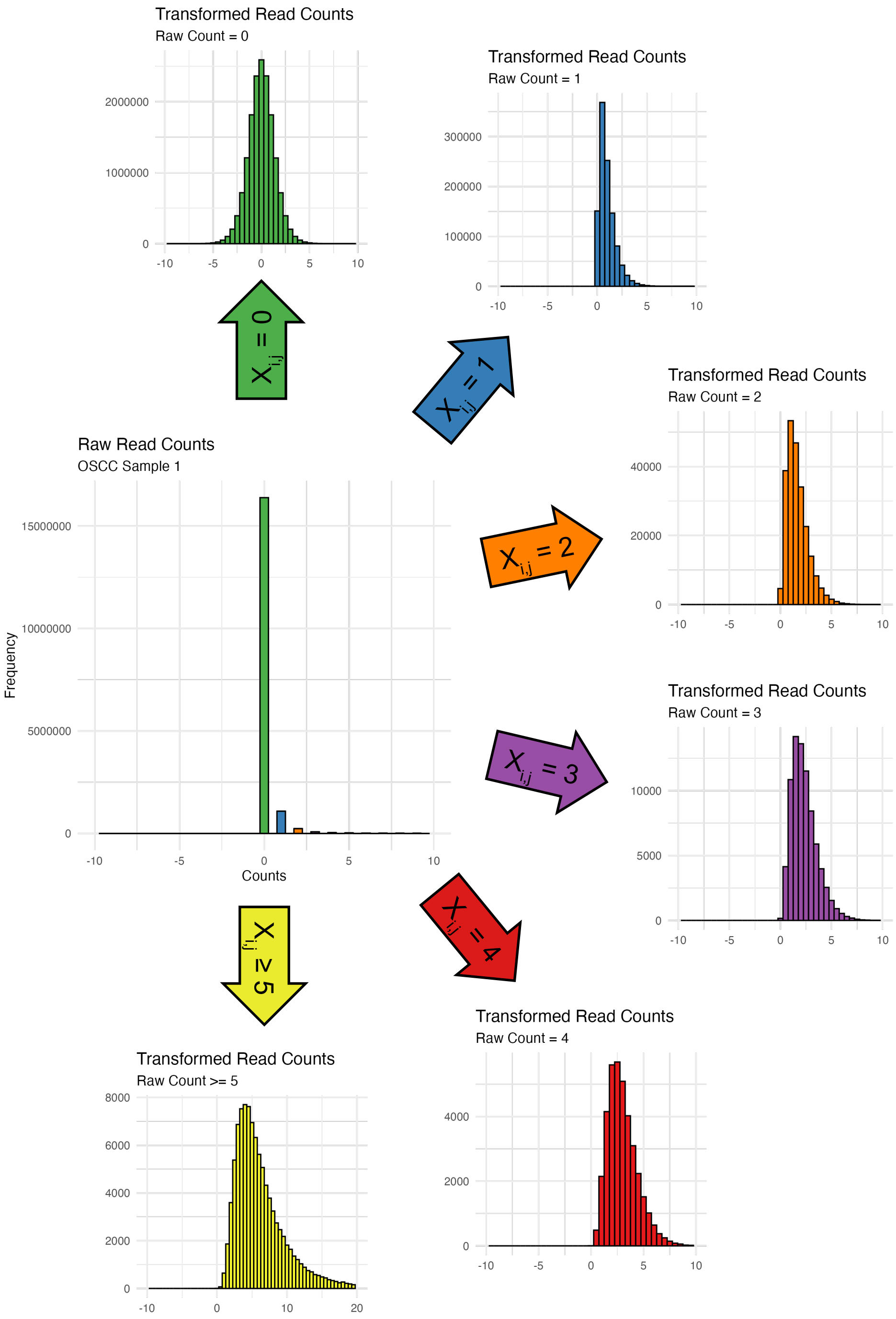


**Figure 42**: Distribution of the gene expression read counts for OSCC Sample 1. Raw read counts are displayed in the central figure. A single posterior sample, split by raw value, is represented on the edges.

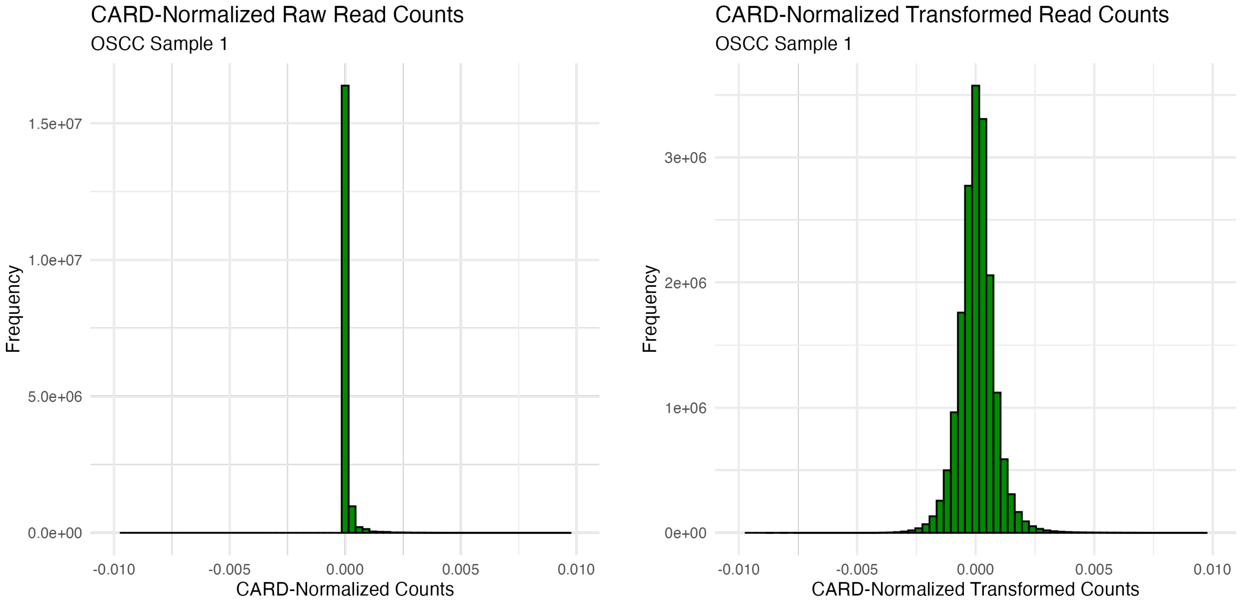


(a) CARD-Normalized Raw Read Counts (b) CARD-Normalized Transformed Read Counts

**Figure 43**: Distribution of the gene expression read counts for OSCC Sample 1 after normalization across spots. a) Raw read counts. b) A single posterior replicate of the transformed read counts.

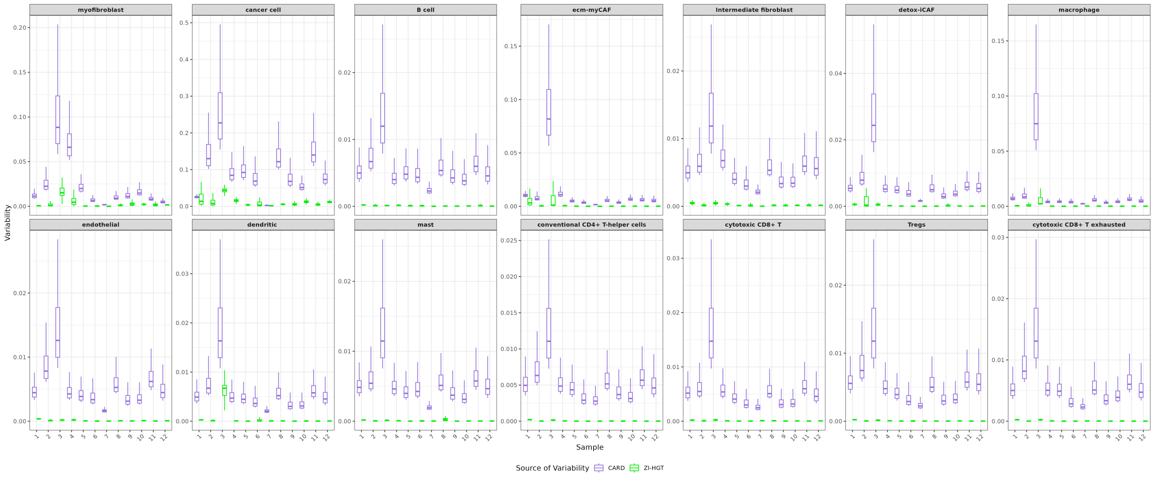


**Figure 44**: Comparison of variability in cell-type proportions estimates from the ZI-HGT and CARD.

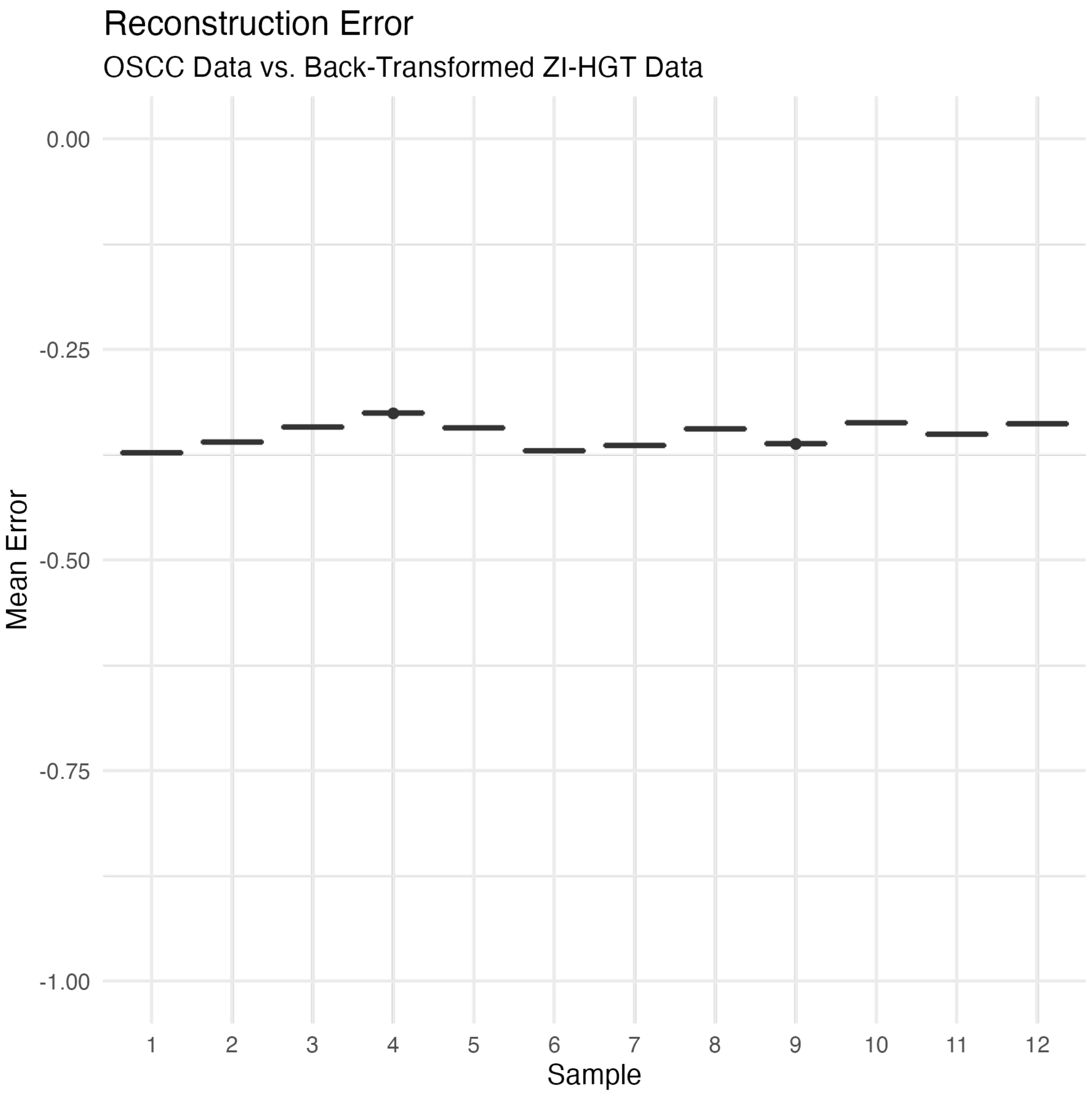


**Figure 45**: Mean reconstruction error per sample between original OSCC data and 10 ZI-HGT–transformed and back-transformed datasets.

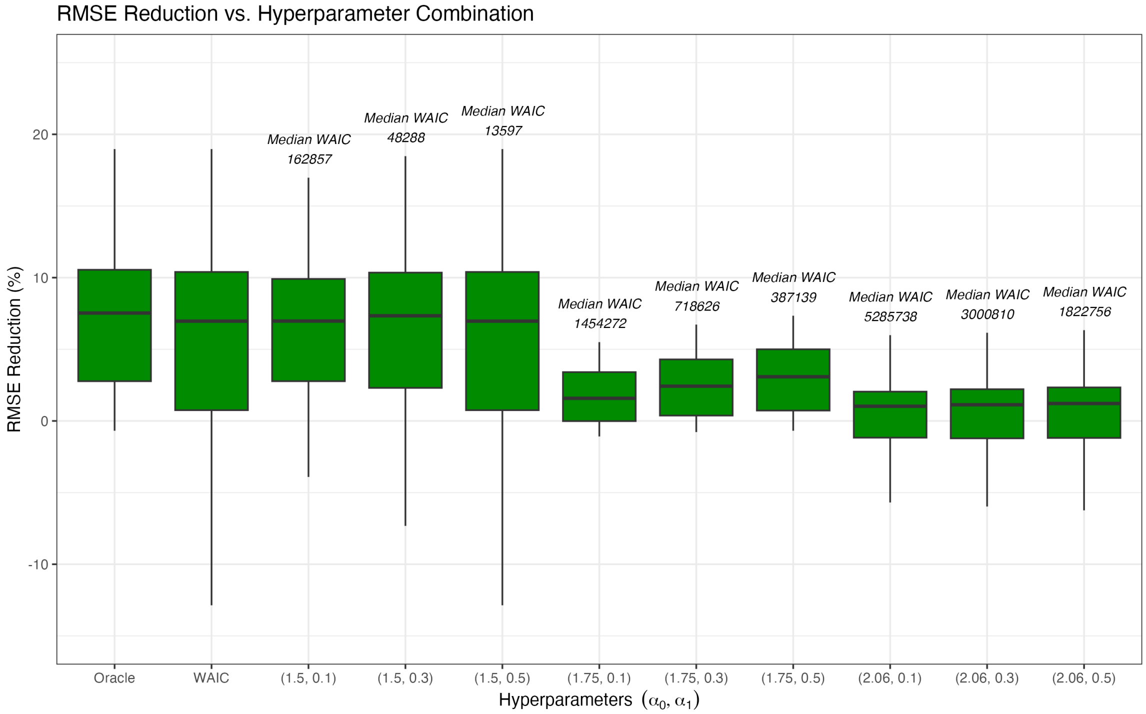


**Figure 46**: RMSE reduction comparing different combinations of hyperparameters. RMSE reduction for the ZI-HGT + CARD across 100 simulated ST datasets for OSCC sample 2. The SPARSim scRNA-seq hyperparameters were set to 0.05 and 50 for the library factor and $\Phi$ factor respectively. The first two boxes show the RMSE reduction for the oracle-chosen (to minimize RMSE) and WAIC-chosen hyperparameters. For all pairs of hyperparameters $(\alpha_0, \alpha_1)$, the median WAIC across all 100 simulated datasets is displayed abovee the boxplot.

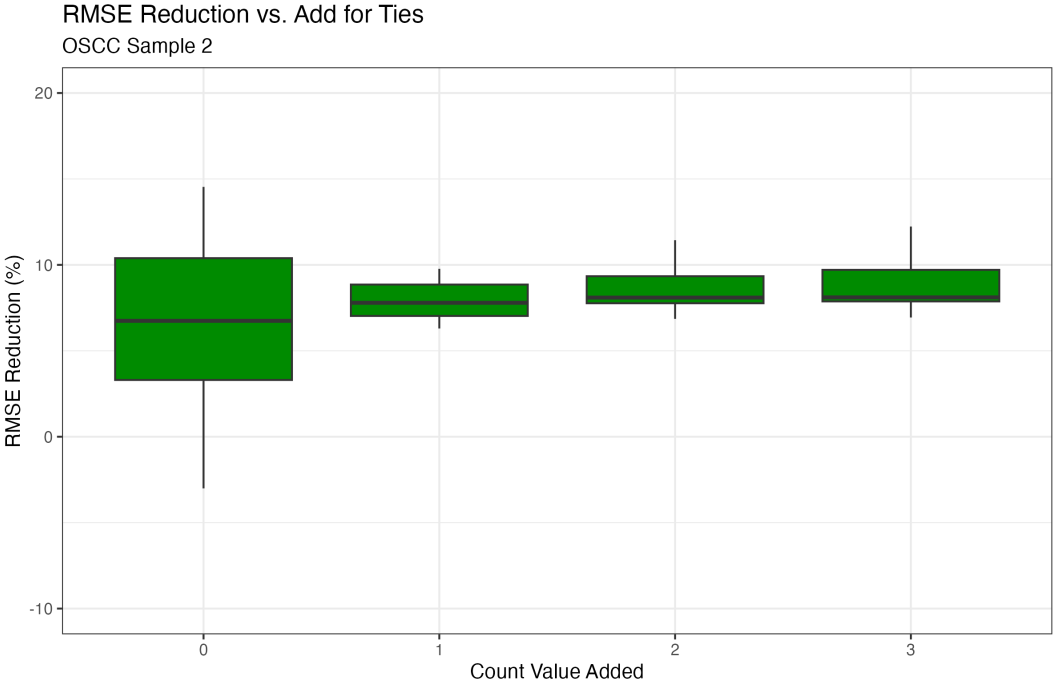


| Count Value Added | Ties at 0 | Ties at 1 | Ties at 2 | Ties at 3 | Ties at 4 | Ties at 5 |
|---|---|---|---|---|---|---|
| 0 | 89.2% | 5.0% | 2.1% | 1.2% | 0.7% | 0.4% |
| 1 | 71.9% | 17.3% | 5.9% | 1.9% | 1.1% | 0.6% |
| 2 | 72.0% | 3.7% | 15.0% | 1.6% | 4.4% | 0.7% |
| 3 | 71.9% | 3.8% | 1.5% | 14.3% | 1.3% | 0.6% |

**Figure 47**: RMSE reduction for the ZI-HGT + CARD when applied to simulated ST data with varied levels of ties at low, nonzero values. Underlying simulated scRNA-seq was constructed such that, for each simulated dataset, one cell type was randomly selected and a randomly selected subset of 50% of all genes was increased by the *Count Value Added* for all cells of that type. The mean proportions of the simulated ST datasets that are equal to 1, 2, 3, 4, or 5 are given in the table, broken down by the count value added. The median improvement for the ZI-HGT + CARD vs. CARD alone ranged from 6.7% for the standard simulated ST data to 8.1% if the count value added was 2 or 3.

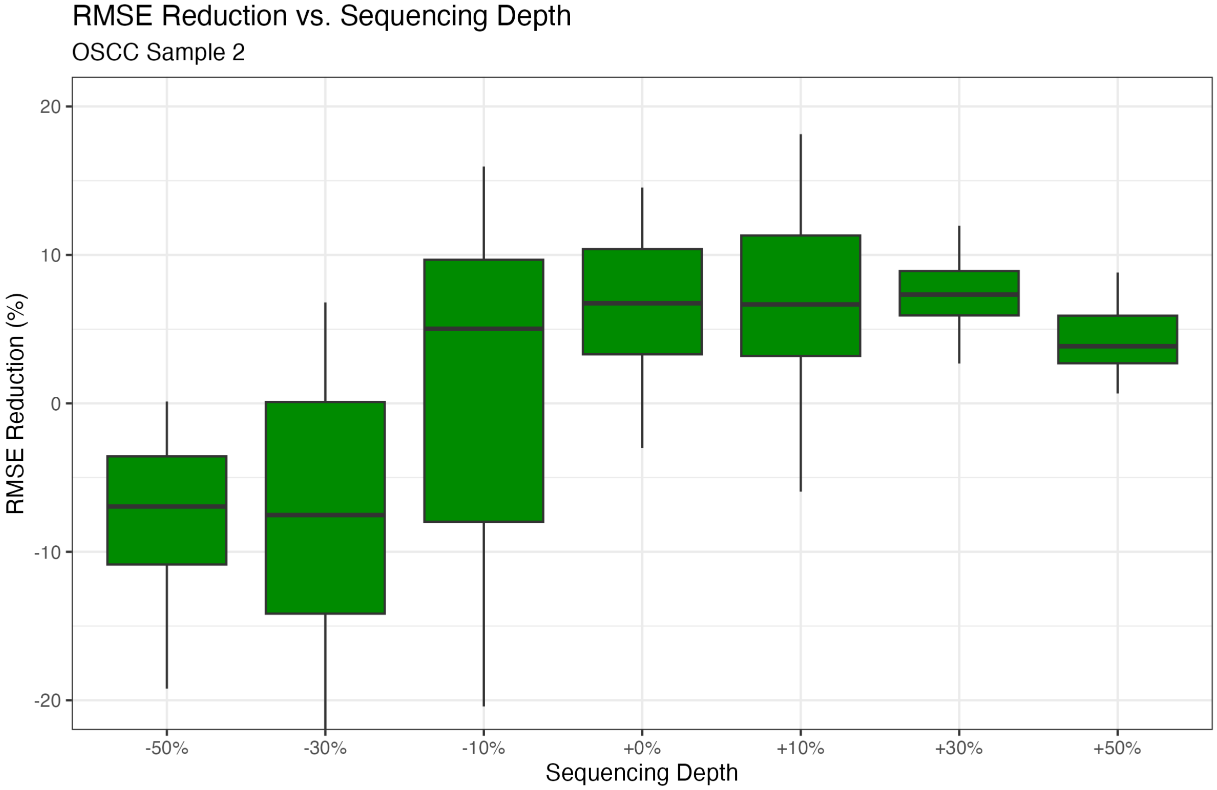


**Figure 48**: reduction vs. sequencing depth. RMSE reduction for the ZI-HGT + CARD across 25 simulated ST datasets for OSCC Sample 2 with -50%, -30%, -10%, +0%, +10%, +30%, and +50% sequencing depth. The SPARSim scRNA-seq hyperparameters were 0.05 and 50 for the library factor and Φ factor respectively. The ZI-HGT + CARD consistently improves performance when sequencing depth is added, but fails to do so when > 10% of sequencing depth is removed.
